# NMR structures of small molecules bound to a model of an RNA CUG repeat expansion

**DOI:** 10.1101/2024.06.21.600119

**Authors:** Jonathan L. Chen, Amirhossein Taghavi, Alexander J. Frank, Matthew A. Fountain, Shruti Choudhary, Soma Roy, Jessica L. Childs-Disney, Matthew D. Disney

## Abstract

Trinucleotide repeat expansions fold into long, stable hairpins and cause a variety of incurable RNA gain-of-function diseases such as Huntington’s disease, the myotonic dystrophies, and spinocerebellar ataxias. One approach for treating these diseases is to bind small molecules to the structured RNAs. Both Huntington’s disease-like 2 (HDL2) and myotonic dystrophy type 1 (DM1) are caused by a r(CUG) repeat expansion, or r(CUG)^exp^. The RNA folds into a hairpin structure with a periodic array of 1×1 nucleotide UU loops (5’C<u>U</u>G/3’G<u>U</u>C; where the underlined nucleotides indicate the Us in the internal loop) that sequester various RNA-binding proteins (RBP) and hence the source of its gain-of-function. Here, we report NMR-refined structures of single 5’C<u>U</u>G/3’G<u>U</u>C motifs in complex with three different small molecules, a diguandinobenzoate (**1**), a derivative of **1** where the guanidino groups have been exchanged for imidazole (**2**), and a quinoline with improved drug-like properties (**3**). These structures were determined using nuclear magnetic resonance (NMR) spectroscopy and simulated annealing with restrained molecular dynamics (MD). Compounds **1**, **2**, and **3** formed stacking and hydrogen bonding interactions with the 5’C<u>U</u>G/3’G<u>U</u>C motif. Compound **3** also formed van der Waals interactions with the internal loop. The global structure of each RNA-small molecule complexes retains an A-form conformation, while the internal loops are still dynamic but to a lesser extent compared to the unbound form. These results aid our understanding of ligand-RNA interactions and enable structure-based design of small molecules with improved binding affinity for and biological activity against r(CUG)^exp^. As the first ever reported structures of RNA r(CUG) repeats bound to ligands, these structures can enable virtual screening campaigns combined with machine learning assisted *de novo* design.

## INTRODUCTION

The vast majority of the RNA that is transcribed encodes for noncoding (nc)RNAs.^1, 2^ NcRNAs function in a variety of normal cellular processes, such as splicing, transcription, translation, degradation, and RNA transport.^3^ However, these RNAs also cause or contribute to various diseases, such as cancers, viral diseases, and neurodegenerative diseases, including those that are caused by RNA repeat expansions.^4^ RNA repeat expansion diseases include amyotrophic lateral sclerosis (ALS), Huntington’s Disease (HD), and myotonic dystrophy types 1 (DM1) and 2 (DM2). DM1 is trinucleotide repeat disorder where an expanded r(CUG) repeat [r(CUG)^exp^] is present in the 3′ UTR of the dystrophia myotonia protein kinase (*DMPK*) gene.^4, 5^ When the number of repeats exceeds 50,^4, 5^ the RNA folds into a long hairpin structure that contains a periodic array of 1×1 UU mismatches.^4, 5^ These motifs bind and sequester various RNA-binding proteins (RBPs) such as the alternative pre-mRNA splicing regulator muscleblind-like splicing regulator 1 (MBNL1).^4, 5^ The resulting r(CUG)^exp^-RBP complexes form foci in the nucleus, leading to poor nucleocytoplasmic transport of *DMPK* mRNA.^6^ Loss of functional MBNL1 leads to aberrant pre-mRNA splicing of its substrates, a hallmark of DM1.^4^ Huntington’s disease-like 2 is also caused by r(CUG)^exp^, which are sequestered into foci containing RBPs.^7, 8^ The result is loss of functional JPH3 protein and pre-mRNA splicing defects that cause disease.

Various modalities have been explored as potential treatments for DM1, including antisense oligonucleotides (ASOs),^9^ CRISPR/Cas9 gene editing,^10^ peptides,^11^ and small molecules. Several small molecule scaffolds have been reported to bind to r(CUG)^exp^ and inhibit formation of r(CUG)^exp^-MBNL1 complexes *in vitro* and in cellular models, including *bis*- benzimidazole derivatives,^6, 12, 13^ a triaminotriazine-acridine conjugate,^14^ pentamidine,^15^ neomycin B,^15^ benzo[g]quinoline,^16^ a perimidine derivative,^17^ and others.^17^ To date, however, there have been no reports of the structures of a small molecule bound to r(CUG) repeats, whether by X-ray crystallography or NMR spectroscopy, although structures of the apo form have been reported.^18–21^ The crystal structure of an RNA is influenced by packing forces and represent a single state of the RNA.^22^ In contrast, RNA structures are intrinsically dynamic and undergo large conformational changes when binding to small molecules.^23^ While NMR spectroscopy can detect solution state dynamics, NMR spectroscopy of RNA is often complicated by spectral overlap due to the small chemical shift dispersion of RNA signals.^24^ Further, the physicochemical properties of the compound, particularly solubility, can confound structural studies.

Our laboratory has constructed a database of small molecules that bind to RNA structures, derived from selection-based experiments or reported in the literature.^25, 26^ Some of the molecules therein elicit effects against their biological targets in patient-derived cells. Three of these molecules were selected that might be amendable to structure determination using NMR spectrometry based on their solubilities. Two-dimensional (2D) NMR spectroscopy and restrained molecular dynamics (MD) were employed to determine the structures of the complexes formed between all three small molecules and a model of r(CUG) repeats.

## METHODS

### Calculation of small molecule physicochemical properties

SwissADME^27^ was used to calculate physicochemical properties such as molecular weight, number of hydrogen bond donors and acceptors, number of rotatable bonds and molar refractivity. Physicochemical properties as well as druglikeness of **1**, **2** and **3** showed the compounds have good druglike properties with no PAINS (pan assay interference structures) alerts (**Table S1**).

### Small molecule inhibition of r(CUG)n-MBNL1 complex formation *in vitro*

A previously reported TR-FRET assay^28^ was used to study the inhibitory effect of **1**-**3** on formation of the r(CUG)12–MBNL1 complex. In summary, 5′-biotinylated r(CUG)12 (80 nM final concentration) was folded in 1× Folding Buffer (20 mM HEPES, pH 7.5, 110 mM KCl, and 10 mM NaCl) by heating at 60 °C followed by slowly cooling to room temperature on the benchtop. The buffer was adjusted to 1× Assay Buffer (20 mM HEPES, pH 7.5, 110 mM KCl, 10 mM NaCl, 2 mM MgCl2, 2 mM CaCl2, 5 mM DTT, 0.1% BSA, and 0.5% Tween-20) and MBNL1-His6 (60 nM final concentration) was then added. The samples were equilibrated at room temperature for 5 min, and then the compound (**1**, **2**, or **3**) was added. After incubating for 15 min at room temperature, streptavidin-XL665 and anti-His6-Tb antibody were added to final concentrations of 40 nM and 0.44 ng/μL, respectively, with a final volume of 10 μL. Samples were incubated for 1 h at room temperature and then transferred to a 96-well plate. Time-resolved fluorescence was measured on a Molecular Devices SpectraMax M5 plate reader. Fluorescence was first measured using an excitation wavelength of 345 nm and an emission wavelength of 545 nm (fluorescence due to Tb). TR-FRET was then measured by using an excitation wavelength of 345 nm, an emission wavelength of 665 nm, a 200 μs evolution time, and a 1.5 ms integration time. The ratio of fluorescence intensity of 545 and 665 nm as compared to the ratios in the absence of ligand and in the absence of RNA were used to determine IC50s.

### Preparation of NMR samples

The oligoribonucleotide r(GACAGC<u>U</u>GCUGUC) was purchased from Dharmacon and de-protected per the manufacturer’s recommended protocol. The oligonucleotide was then de-salted using a PD-10 desalting column containing Sephadex G- 25 resin (Cytiva, catalog #17085101), per the manufacturer’s recommended protocol. After lyophilization to dryness, the RNA was dissolved in Nanopure water. The concentration of the RNA was determined by its absorbance at 260 nm, measured at 90 °C using a Beckman Coulter DU800 UV/Vis spectrophotometer, and the corresponding coextinction coefficient provided by Dharmacon. This stock was then lyophilized to dryness for NMR experiments to afford the desired final concentration, 0.3 mM for samples containing unbound RNA or 0.3 mM and 0.4 mM for samples containing ligand-bound RNA as indicated. The lyophilized RNA was then dissolved in NMR Buffer [5 mM KH2PO4/K2HPO4, pH 6.0, and 0.25 mM EDTA] and folded by heating to 95 °C for 3 min, then slowly cooling the sample to room temperature. D2O was added to the sample to 5% (v/v) to provide a lock signal. For samples collected in D2O, the sample was dissolved in NMR Buffer that was exchanged into 100% D2O and folded by heating to 95 °C for 3 min, and then slowly cooling the sample to room temperature. To prepare ligand-RNA samples for WaterLOGSY (water-ligand observed via gradient spectroscopy) experiments, 15 μM of RNA was mixed with 300 μM of compound to afford a 1:20 RNA:compound ratio.^29–31^ To prepare ligand-RNA samples for 2D NMR spectroscopy, **1**, **2**, and **3** were added to final concentrations of 0.7 mM, 0.6 mM, and 0.6 mM, respectively, where the final concentration of DMSO-d6 was no greater than 5% (v/v). The concentration of the studied RNA:compound complexes was increased to >300 μM for 2D NMR experiments.

### NMR spectroscopy

NMR spectra of samples in Shigemi tubes (Shigemi, Inc.) were acquired on Bruker Avance III 600, 700, 850, and 900 MHz spectrometers. 1D and 2D spectra were acquired on the apo and ligand-bound forms at 5 °C, 6 °C, 15 °C, 25 °C, 30 °C, and 35 °C. NMR experiments with RNA samples in 95% H2O/5% D2O were carried out at 5 °C, 6 °C, and/or 15 °C. NMR experiments in with RNA samples in 100% D2O were carried out at 25 °C, 30 °C, and/or 35 °C. Excitation sculpting was applied during acquisition of 1D NMR spectra to suppress the water signal.^32^ WaterLOGSY experiments were carried out on samples containing the r(CUG) duplex construct and each compound.^29–31^ WaterLOGSY signals were phased to give negative signals for negative NOEs relative to water. Proton chemical shifts were referenced to water. 2D ^1^H-^1^H NOESY spectra of samples in 95% H2O/5% D2O were collected with a 125 ms mixing time. 2D ^1^H-^1^H NOESY spectra of samples in 100% D2O were collected with 100, 170, and/or 175 ms mixing times and, separately, at a 400 ms mixing time. 1D NMR spectra were processed using Bruker TopSpin® and 2D NMR spectra were processed with TopSpin or NMRPipe^33^ and assigned with NMRFAM-SPARKY.^34^ The solubility of small molecules in complex with RNA was measured under NMR conditions by titrating compound into a solution containing 50 μM of RNA and NMR Buffer using up to a 6:1 molar ratio of compound:RNA. After addition of each aliquot of compound, a 1D NMR spectrum was acquired. If the RNA peaks remained visible and compound peaks appeared in the aromatic region, then a higher concentration sample of the RNA-compound complex was prepared for 2D NMR spectroscopy.

### Methods for obtaining distance and dihedral restraints

Spectral peaks (NOEs) for **1**-, **2**-, and **3**-bound r(CUG) complexes were assigned using an NMR solution structure for the apo RNA by comparing spectra of free form and ligand-bound RNA. Distance restraints for pairs of protons were calculated by integrating NOE volumes in SPARKY^35^ or manually assigned a range of distances based on relative NOE intensities in 100 to 200 ms mixing time NOESY spectra to reduce the influence of spin diffusion.^36^ Distance restraints for ligand-RNA NOEs, which were weak in shorter mixing time spectra, were assigned in the same manner using 400 ms mixing time NOESY spectra. NOE volumes were referenced to those calculated from fixed distances: H2′-H1′ (2.75 Å); cytosine or uracil H5-H6 (2.45 Å).^37^ Hydrogen bonds in canonical AU, GC, and GU pairs were assigned to distances of 2.1 ± 0.3 Å. NOE intensities between H1′ and H6/H8 indicate that all residues adopted an *anti* conformation. Therefore, the *χ* dihedral angle was constrained between 170° and 340° (*anti*) for all residues except terminal residues and uridine nucleotides comprising the internal loop in the apo structure.

### Modeling methods

Structures were calculated with a simulated annealing protocol, using the lowest free energy structure of NMR-restrained apo-r(CUG) structure to which compounds were added to the binding site in an arbitrary orientation using UCSF Chimera.^38^ LEaP was then used to generate topology and coordinate files.^39^ The starting structure was then energy minimized using AMBER.^39^ Restrained molecular dynamics simulations were carried out with AMBER^39^ using the parm99χ_YIL force field.^40^ All distance restraints, both inter- and intra- molecular, for each RNA-small molecule complex can be found in **Tables S2 – S7**. Solvation was simulated with the general Born implicit model and 0.1 M NaCl.^41^ The system was heated from 0 to 1,000 K in 5 ps, cooled to 100 K in 13 ps, and then to 0 K in 2 ps. Force constants were 20 kcal mol^−1^ Å^−2^ for NOE restraints during the simulated annealing protocol. The calculation was repeated 100 times. The structures generated from the calculations were sorted based on NOE distance violations, and the 20 structures with the fewest violations were selected. The top twenty structures were superimposed and their RMSD determined using VMD.^42^

### Calculation of global distortions of RNA (curvature)

In order to measure the global deformation of the RNA before and after ligand binding, the overall bend angle (global curvature) was calculated using *Curves+*.^43^ The global curvilinear helical axis was calculated from the average local axes applying a polynomial smoothing. NMR ensemble with 20 structures was used for this calculation.

## RESULTS & DISCUSSION

### Compound selection

To begin to uncover the principles of small molecule recognition of RNA structures, our laboratory has employed a selection-based method to identify selective RNA motif-small molecule interactions, which are collected into a database along with interactions reported in the literature.^25, 26^ Indeed, these interactions have guided design of molecules that modulate cellular RNAs in patient-derived cells. Previously, **1** was discovered to bind the 1×1 nucleotide AA internal loops formed by expanded r(CAG) repeats, or r(CAG)^exp^ and inhibit complex formation of the RNA with MBNL1,^44^ the cause of spinocerebellar ataxia type 3 (SCA3).^4^ To improve the binding properties of **1** and perhaps direct its recognition to r(CUG)^exp^, a derivative of the compound was synthesized, or **2**. In **2**, the imidazolines replaced the guanidines of **1** to reduce the polarity of the compound while substitution of the ester linkage with an amide linkage increases compound stability and imparts conformational rigidity on the compound.^45^ Preorganization of the ligand chemically constrains its structure, reducing the entropic cost of binding to the target RNA.^46, 47^ Compound **3** was a screening hit that bind to the A-bulge present in a mutant splicing regulatory element (SRE) that controls *MAPT* pre-mRNA alternative splicing, the presence of which causes frontotemporal dementia with parkinsonism-17 (FTDP-17). The small molecule was lead optimized by via pharmacophore modeling and chemical similarity searching.^48^

1D NMR experiments were then carried out on these compounds to determine if their solubilities were sufficient under NMR conditions for structure determination. At 50 μM of RNA and up to 300 μM of compound, resonances for each compound appeared in the aromatic proton region while imino proton resonances shifted or broadened. Appearance of the aromatic proton resonances combined with observed changes (broadening and shifting) of resonances in 1D spectra upon addition of compounds **1** to **3** showed that these compounds were amenable to structural studies using NMR spectroscopy.

### Small molecules 1, 2, and 3 inhibit r(CUG) repeat-MBNL1 complex formation

The interaction of r(CUG)^exp^ with MBNL1 and other RBPs triggers the formation of nuclear foci in disease-affected cells.^49^ Therefore, as a secondary validation of compound binding to the RNA from 1D NMR spectra, the ability of **1**, **2**, and **3** to disrupt the r(CUG)^exp^-MBNL1 complex *in vitro* was measured by using a previously reported time-resolved fluorescence resonance energy transfer (TR-FRET) assay.^28^ In this workflow, the RNA–protein complex was pre-formed followed by addition of the compound of interest at various concentrations to afford an IC50. The IC50s for **1**, **2**, and **3** in this assay were 7.9 ± 0.3, 18.9 ± 0.5 and 100.4 ± 6.8 μM, respectively (**Figure S1**). Of note, the affinity of MBNL1 for r(CUG) repeats, for which the small molecules compete, is 7.2 ± 1 nM.^50^

### NMR analysis of the apo structure of a model of r(CUG) repeats

Previous NMR and modeling studies with RNA constructs containing one and three tandem 5′C<u>U</u>G/3′G<u>U</u>C motifs show that the 1×1 UU loops are dynamic.^21, 51, 52^ In this study, we designed a self-complementary RNA construct (duplex) containing one 5′C<u>U</u>G/3′G<u>U</u>C motif and five base pairs on either side to stabilize the helical region. The RNA sequence was optimized to reduce overlap of NMR resonances.

NMR spectra of the free, unbound form of the r(CUG) duplex indicated formation of the 5’C<u>U</u>G/3’G<u>U</u>C internal loop (where the underlined Us indicate loop nucleotides) flanked by helical regions. In particular, a sequential H6/H8-H1′ walk, in addition to inter-residue H6/H8-H2′ NOEs, were assigned for all residues in NOESY spectra, indicating that the duplex adopted an overall A-form structure (**Figures S2 – S3** and **Tables S2 and S8**). Exchangeable protons were assigned in a spectrum collected in 95% H2O/5% D2O at 5 °C. Cross-peaks between adenine H2 protons and uracil imino protons were assigned for all AU pairs while cross-peaks between guanine imino and cytosine amino protons were assigned for all GC pairs except for the terminal GC pair. Intrastrand and interstrand NOEs from guanine H1 to the 3′ H1′ were observed for the same GC pairs. Specifically, NOEs were observed from G5H1 to C6H1′ and U10H1′ and G11H1 to U12H1′ and A4H1′ in both constructs. NOEs from U7H3 to C6H1′, G8H1′, and G8H1, C6H2′ to U7H6, and U7H1′ and U7H2′ to G8H8 suggest that the loop uridines stack in the helix. This is also supported by the observation of the upfield shifted U7H3 proton in the 10-12 ppm range in the 1D and 2D water experiments, in agreement with chemical shifts of imino protons in 1×1 UU mismatches.^53^ These data are consistent with previously reported structures.^22, 47, 48^ The presence of the U7H3 proton can be used to evaluate the binding of compounds to the r(CUG) loop. Retention of the U7H3 resonance would be an indicator that the small molecule binds in a way that does not interfere with the UU mismatch and or stacking. Loss of the U7H3 resonance would indicate disruption of the UU mismatch with possible displacement of one or both Us.

### Solution structure of the apo RNA

Structures of apo r(CUG) motif were calculated using 246 distance restraints to generate a final ensemble of 20 structures (see Methods section). All 20 structures in the final ensemble adopted an A-form geometry, in agreement with the 2D NMR spectra described above (**Figures 1** **and S4** and **Table S8**). The average RMSD for the ensemble of 20 structures was 1.25 ± 0.33 Å (**Table 1**), indicating good convergence. The average helical rise and twist were 2.8 ± 0.3 Å and 30.9 ± 2.0°, respectively. Overall, the helix is slightly under-twisted compared to A-form RNA, which had an average helical rise and twist of 2.8 Å and 32°, respectively.^21, 54^ The average χ dihedral angles ranged from −151 to −167° for all residues, corresponding to *anti* conformations.

**Figure 1:**
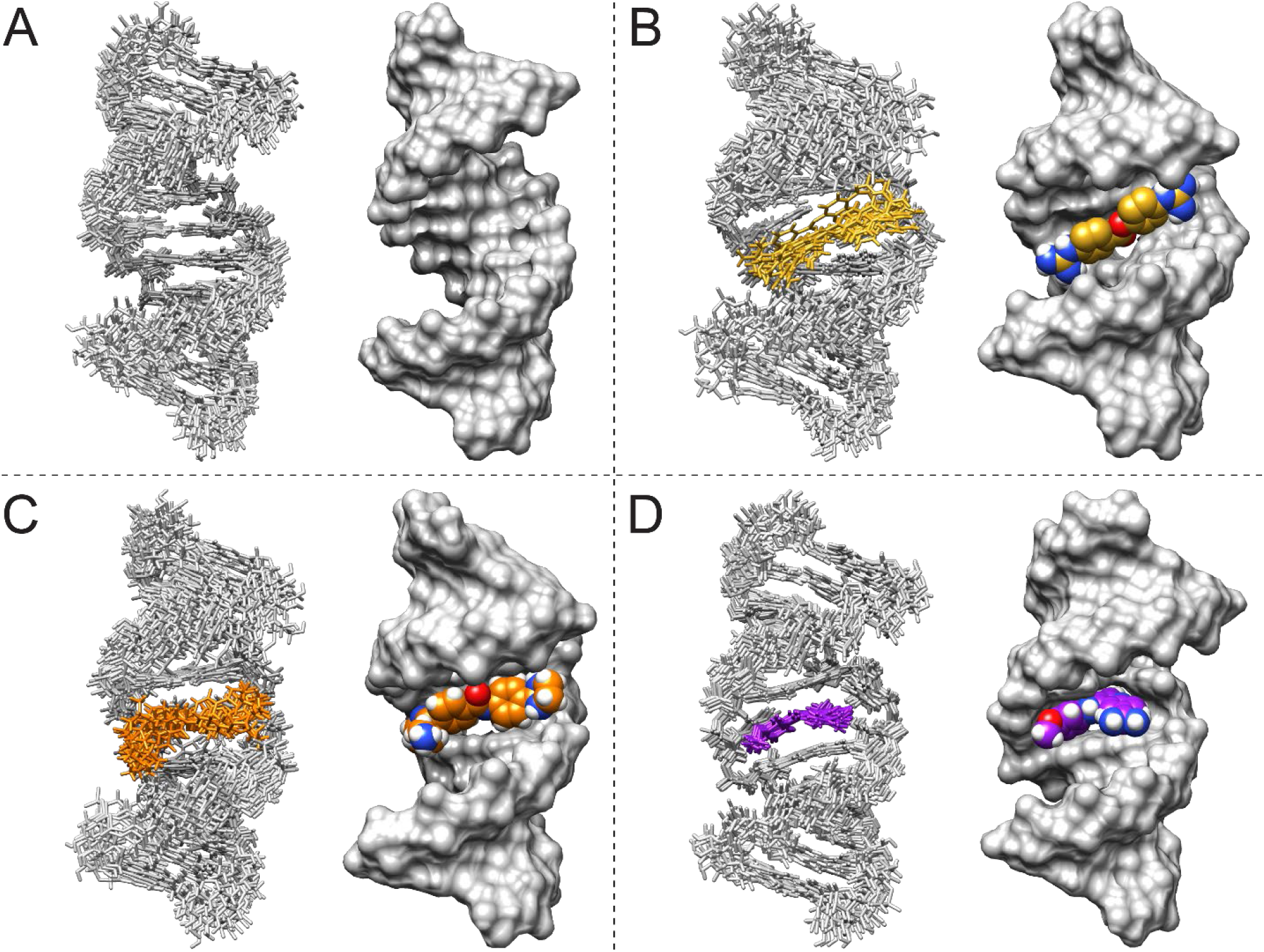
NMR solution structures of unbound r(CUG) duplex and r(CUG) duplex bound to compounds 1-3. (A) Structure of unbound r(CUG). (B) Structure of the r(CUG)-**1** complex. (C) Structure of the r(CUG)-**2** complex. (D) Structure of the r(CAG)-**3** complex. For each model, an overlay of the 10 structures with the fewest distance restraint violations in stick representation (left) and a surface representation of the structure with the fewest distance restraint violations (right) are shown.

**Table 1:** Structural refinement statistics for the average of 20 structures for unbound and ligand bound RNAs.

|  | Unbound<br>r(CUG) | r(CUG)-1 | r(CUG)-2 | r(CUG)-3 |
| --- | --- | --- | --- | --- |
| No. of restraints |  |  |  |  |
| All NOE restraints and hydrogen bonds | 246 | 336 | 292 | 358 |
| All NOE restraints | 214 | 284 | 266 | 326 |
| Intraresidue/compound intramolecular | 118 | 134 | 122 | 161 |
| Sequential residues | 88 | 126 | 104 | 133 |
| Long range, excluding RNA-compound | 8 | 0 | 17 | 5 |
| RNA-compound | N/A | 24 | 23 | 27 |
| Hydrogen bond | 32 | 52 | 26 | 32 |
| Dihedral/chiral restraints | 130 | 0 | 0 | 0 |
| RMSD of experimental restraints |  |  |  |  |
| Distance (Å) | $7.9 \times 10^{-4}$ | $1.0 \times 10^{-3}$ | $9.3 \times 10^{-4}$ | $2.8 \times 10^{-3}$ |
| Dihedral (deg) | 0.3 | 0.0 | 0.0 | 0.0 |
| RMSD of structures for heavy atoms (Å) |  |  |  |  |
| All residues, excluding compound | $1.25 \pm 0.33$ | $1.81 \pm 0.43$ | $1.87 \pm 0.48$ | $1.01 \pm 0.19$ |
| All residues and compound | N/A | $1.86 \pm 0.42$ | $1.89 \pm 0.48$ | $1.02 \pm 0.20$ |
| Outer helices (residues 1-5 and 9-13) | $1.22 \pm 0.36$ | $1.88 \pm 0.49$ | $1.75 \pm 0.53$ | $0.93 \pm 0.22$ |
| Compound | N/A | $1.02 \pm 0.25$ | $0.94 \pm 0.30$ | $0.46 \pm 0.29$ |
| Internal loop and closing pairs (residues 6-8), excluding compound | $0.78 \pm 0.30$ | $0.94 \pm 0.28$ | $1.65 \pm 0.62$ | $0.79 \pm 0.18$ |
| Internal loop and closing pairs (residues 6-8) and compound | N/A | $1.18 \pm 0.31$ | $1.65 \pm 0.59$ | $0.80 \pm 0.15$ |

Among the 20 ensemble structures, interbase hydrogen bonding of the UU mismatch caused the helix to be under-twisted at one step (5’C<u>U</u>/3’G<u>U</u>) and over-twisted at the adjacent step (5’<u>U</u>G/3’<u>U</u>C) within the 5’C<u>U</u>G/3’G<u>U</u>C motif. In particular, the helical twist at the 5′C<u>U</u>/3′G<u>U</u> and 5′<u>U</u>G/3′<u>U</u>C steps were 27.7 ± 11.1° and 33.2 ± 11.7°, respectively. The UU loop contained a *cis*-Watson Crick/Watson Crick UU pair^55^ closed by canonical GC pairs (**Figures 1** and **S4**), stabilized by stacking and hydrogen bonding interactions. The average number of hydrogen bonds in the UU pair among all 20 ensemble structures was 1.4 ± 0.5. A one hydrogen bonded UU pair was observed in eight of the structures (40%), and a two hydrogen bonded UU pair was observed in 12 of the structures (60%), indicating that the loop’s structure is dynamic and interconverting between these hydrogen bonding structures. Hydrogen bonding results in a small reduction of the average C1′-C1′ distance for the UU mismatch to 10.0 ± 0.5 Å from an average of ∼10.5 Å for A-form RNA, although the distance is still within range of that observed in A-form RNA.^20, 56^ The UO4 to UH3 distance was 1.8-1.9 Å in 19 of the 20 structures and >3.4 Å in the remaining structure. The average buckle, an indication of cusp formation,^57^ for the UU mismatch was 1.3 ± 6.2°, which is less than values of 5.8 ± 6.4° and −4.2 ± 4.8° calculated for the flanking GC pairs. The opening, or angle between two bases,^57^ and stretch, an indication of separation of two bases,^57^ of the UU mismatch (−10.3 ± 9.6° and −1.6 ± 0.1°, respectively) were of the greatest magnitude among all paired nucleotides in the helix. The relatively large variation for the opening in the UU mismatch is consistent with interconversion of the UU pair between one and two hydrogen bond states. In summary, the unbound r(CUG) motif contains a UU pair, stabilized by one or two hydrogen bonds, that induces distortions in the helix.

The exchange between one- and two-hydrogen bond states is consistent with that observed in r(3×CUG).^21^ The average number of hydrogen bonds in UU pairs calculated by X3DNA is 1.4 ± 0.5 and 1.5 ± 0.5 for the single r(CUG) and r(3×CUG) constructs, respectively. In both constructs, hydrogen bonding in UU mismatches was facilitated by shorter C1′-C1′ distances. As in the single r(CUG) construct, the helix is over-twisted and under-twisted at adjacent steps of the outer 5’C<u>U</u>G/3’G<u>U</u>C motifs in the r(3×CUG). Both base pair steps of the middle 5′C<u>U</u>G/3′G<u>U</u>C motif of r(3×CUG) are under-twisted; overtwisting instead occurs at the flanking 5′GC/3′CG motifs. In comparison, Parkesh *et al*. observed exchange between zero, one, and two hydrogen bond states in their structure of r(1×CUG), with a shortened C1′-C1′ distance in the two hydrogen bond state. DSSR analysis of their structures showed that for the one hydrogen bond state, slight undertwisting occurs at the 5′C<u>U</u>/3′G<u>U</u> and 5′<u>U</u>G/3′<u>U</u>C steps and overtwisting occurs at the flanking 5′GC/3′CG motifs.^58^ In the two hydrogen bond state, the helical twist (31.5°) at the 5′C<u>U</u>/3′G<u>U</u> step is close to average but the helix is over-twisted (36.7°) at the 5′<u>U</u>G/3′<u>U</u>C step and under-twisted at the flanking 5′GC/3′CG steps (28.3° and 25.0°).

Backbone distortion was not observed in a de-twinned crystal structure of r(6×CUG) that contained a one hydrogen bond stretched UU wobble pair.^59^ Separately, a construct containing r(2×CUG) contains a stretched wobble pair, with one hydrogen bond in the upper 5′C<u>U</u>G/3′G<u>U</u>C motif and two hydrogen bonds in the lower 5′C<u>U</u>G/3′G<u>U</u>C motif. A shortened C1′-C1′ distance of 8.59 Å was observed in the lower 5′C<u>U</u>G/3′G<u>U</u>C motif to accommodate the two hydrogen bond state 5’C<u>U</u>G/3’G<u>U</u>C motif.^60^ In the X-ray structure of another r(3×CUG) construct, the outer UU mismatches each contained one direct hydrogen bond, which was the predominant conformation, and the central UU mismatch contained a water-bridged hydrogen bond.^61^ The C1′-C1′ distance was 10.8 Å in each of the outer UU mismatches and 10.3 Å in the central UU mismatch.^61^ In summary, X-ray and NMR structures of 5′C<u>U</u>G/3′G<u>U</u>C motifs show that the UU mismatches adopt one or two hydrogen bond states with helical distortion around the r(CUG) motif. The structural features of r(CUG) repeats could allow small molecules to recognize and form specific contacts with r(CUG) motifs.

Analysis of ensembles of NMR structures of fully base paired duplexes using X3DNA^62^ showed no significant differences in the values of helical parameters compared to those of apo- r(CUG) or r(3×CUG). The average helical rise and twist values for a dodecamer, r(CGCAAAUUUGCG)2 (PDB 1AL5) are 2.5 ± 0.6 Å and 33.6 ± 3.1°, respectively.^63^ For the decamer, r(5′-GAAGAGAAGC/3′CUUCUCUUCG) (PDB 1RRR),^64^ the average helical rise and twist are 2.5 ± 0.3 Å and 32.0 ± 2.2°, respectively. All of these values are within error of those observed for apo-r(CUG) (see above) and r(3×CUG) (2.7 ± 0.5 Å for helical rise and 30.8 ± 3.7° for helical twist).^21^ For the decamer, r(5′-GCAGAGAGCG/3′-CGUCUCUCGC) (PDB 2JXQ),^65^ the average helical rise and twist are 2.4 ± 0.3 Å and 33.7 ± 2.4°, respectively. The average C1′-C1′ distance for each base pair was in the 10.7-10.8 Å range in PDB 1AL5,^63^ and the 10.7-10.9 Å range for PDB 1RRR,^64^ and 10.4-10.6 Å for PDB 2JXQ, which as expected for A-form RNA.^65^ As noted earlier, r(CUG)-containing RNAs typically have shorter C1′-C1′ distances, particularly for two hydrogen bond UU pairs. Average major groove width values ranged from 18.4-19.7 Å for PDB 1AL5, 19.3-21.8 Å for PDB 1RRR, and 16.4-16.9 Å for PDB 2JXQ. In comparison, these values ranged from 18.9-19.6 Å for apo-r(CUG) and from 17.8-20.6 Å for r(3×CUG). Thus, the helical structures of apo-r(CUG) and r(3×CUG) appear to be consistent with those of fully base paired duplexes. That said, larger variations in the C1′-C1′ distances and number of hydrogen bonds are observed for mismatched UU pairs in the apo-r(CUG) and r(3×CUG) RNAs than for canonical base pairs in the same constructs and the fully base paired constructs. Thus, small molecules could selectively recognize the localized dynamics in the UU pairs, rather than the helical structure it shares with fully paired RNAs, and form favorable interactions that compete with MBNL1 binding.

### NMR analysis of 1D and WaterLOGSY spectra

As a first step, the molecules were studied in a WaterLOGSY experiment. WaterLOGSY spectra of **1**, **2**, and **3** in the absence of RNA gave positive NOEs, indicating no aggregation of the compounds. Addition of r(CUG) duplex to each compound gave negative NOEs (positive WaterLOGSY signals) for the compounds (**Figures S5 – S7)**. WaterLOGSY spectra of **1**, **2**, and **3** in the presence of RNA also gave RNA signals with negative intensity. Thus, these negative NOEs resulted from small molecule binding to the RNA, rather than aggregation of the compounds. Thus, these spectra indicate that the RNA-small molecule complexes were soluble.

Complementarily, imino proton spectra of the duplex model were collected as a function of small molecule concentration. For **1**, addition of compound to 1:1 **1**:RNA ratio caused minor shifting of the U10H3 and G5H1 resonances, suggesting that the compound is binding to the r(CUG) motif (**Figure S8**). At this ratio, U7 broadened significantly while G8H1 was mostly unchanged. At 4:1 **1**:RNA, U7H3 was broadened while G8H1 continued to shift upfield, indicating improved stacking interactions in the r(CUG) motif and slight displacement of the uracils from the helix. At this ratio, G1H1 also began to shift upfield, indicating stacking of the compound at the end of the helix. Addition of the small molecule to 6:1 **1**:RNA molar ratio caused U10H3 to shift further downfield and U12H3 and G5H1 to shift upfield, respectively, relative to the unbound state. Within the CUG motif, G8H1 continued to shift upfield from its position at the 4:1 **1**:RNA ratio and broadened. Although these shifts may be minor, the shifting and broadening of peaks, particularly around the r(CUG) motif, collectively indicate a structural shift upon compound binding. These results also show that the compound selectively binds to r(CUG) over base paired regions.

Addition of **2** to 2:1 **2**:RNA resulted in broadening of G5H1, U7H3 and G8H1, but the resonances did not otherwise shift. At 4:1 **2**:RNA, U7H3 and G8H1 broadened further and G8H1 shifts downfield, towards G5H1, indicating binding of the compound to the r(CUG) motif. Meanwhile, U10H3 shifted downfield relative to the unbound duplex. At 6:1 **2**:RNA molar ratio, addition of **2** resulted in a further downfield shift of U10H3 relative to unbound but G5H1, U7H3, and G8H1 do not further broaden. This indicates saturation of compound at the r(CUG) motif at the 4:1 **2**:RNA molar ratio. G1H1 and G11H1 remained overlapped upon addition of compound to 6:1 **2**:RNA ratio (**Figure S9**). The shifting and broadening of peaks suggest that the compound binds to the r(CUG) motif.

Addition of **3** at a 1:1 ratio of compound:RNA shifted G5H1 and G8H1 upfield and U7H3 and U10H3 downfield. At 4:1 **3**:RNA ratio, G5H1 shifted further downfield, G8H1 shifted further upfield, and U7H3 broadened. This suggests that **3** disrupts the UU base pair at this ratio. At 6:1 compound:RNA ratio, G8H1 continued to shift upfield while U7H3 remained broadened.

Otherwise, resonances in the r(CUG) motif do not differ much at the 4:1 and 6:1 **1**:RNA ratios, indicating saturation of binding at that site. Away from the r(CUG) motif, shifts in G11H1 and U12H3 were less pronounced throughout the titration. However, G1H1 is distinguishable from G11H1 4:1 **3**:RNA ratio, which may result from stacking of the compound at the end of the helix (**Figure S10**).

Collectively, both the WaterLOGSY and imino proton spectra suggest that the small molecule binds to the RNA with sufficient affinity and selectivity to determine the structures of the complexes.

### NMR analysis of r(CUG) bound to 1 and 2

As discussed above, an uninterrupted NOESY walk with sharp cross-peaks was observed for the apo form of the r(CUG) repeat duplex model. Interestingly, an uninterrupted NOESY walk was also observed in the spectra collected for the duplex in the presence of related small molecules **1** and **2** (**Figures 2** **– 3** and **S11 – S12**). Further, these cross-peaks remained sharp (similar to those observed in the apo spectra), suggesting fast exchange among multiple states of the complexes. A total of 24 NOEs between the RNA and **1** were observed for the r(CUG)-**1** complex (**Figure 2** and **Table S9**). Among them, NOEs were observed from H1 and H3 aromatic protons of **1** to sugar protons of C6, and G8 and to aromatic protons of C6, U7, and G8. On the opposite side of the **1**, NOEs were observed from H5 and H7 aromatic protons of 1 to aromatic and sugar protons of U20 and G21. However, no NOEs between **1** and the loop uracils were observed, Thus, **1** appears to bind to the major groove of the RNA, between the closing GC pairs of r(CUG) motif and disrupts base pairing and/or stacking interactions of the UU mismatch.

**Figure 2:**
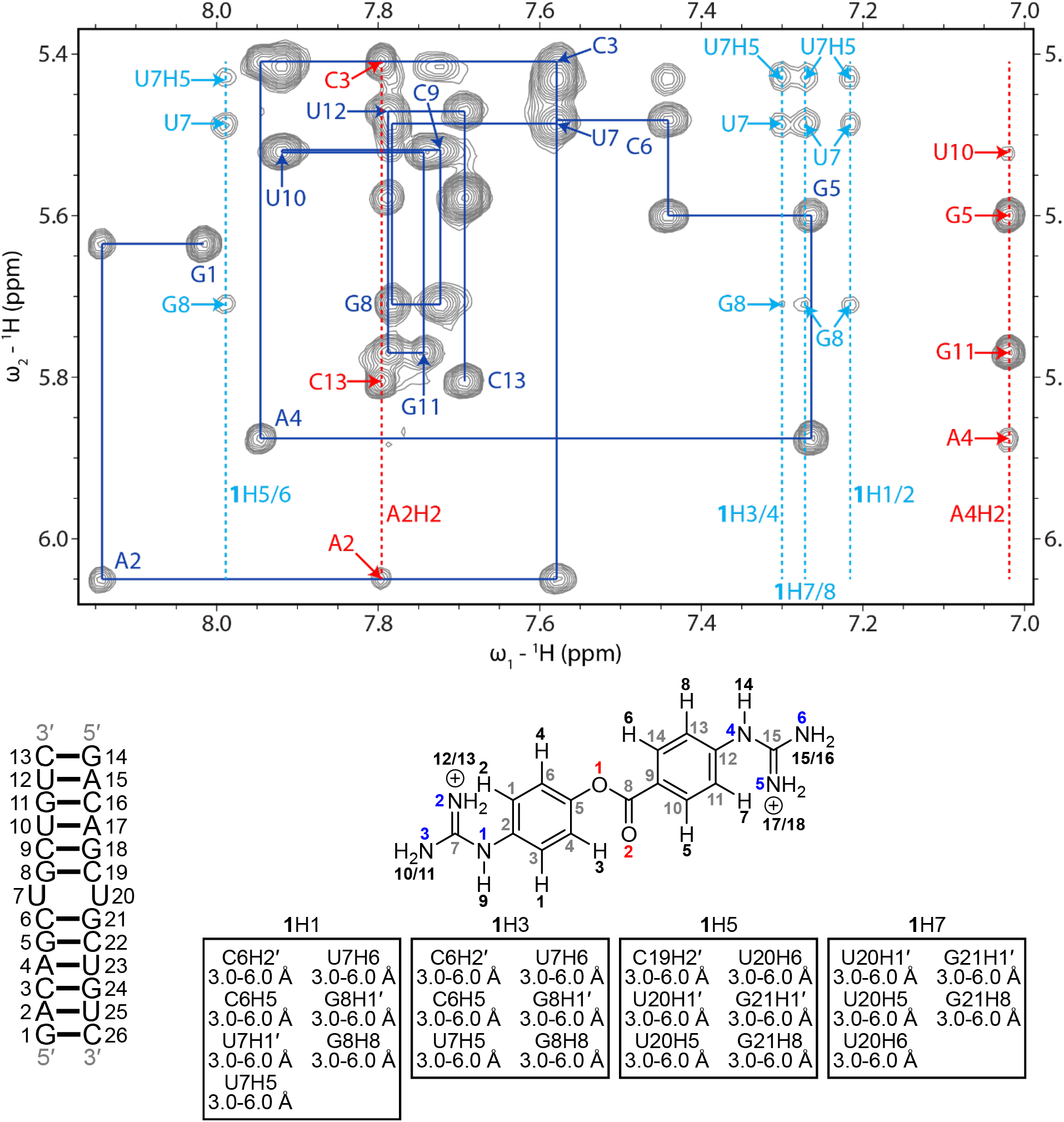
H6/H8-H1′ region of a 2D ^1^H-^1^H NOESY spectrum of the r(CUG)-1 complex. Blue lines represent a sequential H6/H8-H1′ walk, and blue labels represent intraresidue H6/H8 to H1′ NOEs. Dashed red lines represent adenine H2 resonances, and red labels represent interresidue and intraresidue NOEs between adenine H2 and H1′ of nearby residues. Dashed light blue lines represent aromatic proton resonances of **1**, and light blue labels correspond to NOEs between H1′of r(CUG) residues, unless otherwise noted, and **1**. RNA residues are only numbered from 1 to 13 in the NMR spectrum, and residues 14 to 26 correspond to residues 1 to 13 due to the symmetry of the RNA construct. In the chemical structure of **1**, black numbers represent hydrogens, gray numbers represent carbons, blue numbers represent nitrogens, and red numbers represent oxygen. NOEs between the RNA and **1** and their corresponding distance restraints used for modeling are colored according to the atoms of **1**. The spectrum was acquired at 25 °C with 400 ms mixing time and 0.3 mM of RNA and 0.6 mM of **1**.

In the r(CUG)-**2** complex, 23 NOEs between r(CUG) and **2** were observed (**Figure 3** **and Table S10**), six of which are also observed between r(CUG) and structurally similar compound **1**: H1 of **2** to U7H5 and U7H6; H5 of **2** to C19H2′, U20H5, U20H6; H7 of **2** to U20H5. In addition to these and other NOEs between **2** and sugar and aromatic protons of r(CUG), **2** has NOEs from the aromatic H4 and H6 protons of **2** to U7H3 and U20H3, respectively, from one of the imidazoline groups of **2** to U20H6 and G21H5″, and from the H26 amide proton of **2** to U7H3. These NOEs between the compound and the imino protons of the loop uracils laid within the 10- 12 ppm imino proton range, where U7/U20H3 of the apo structure resonates, suggesting that the loop uracils remain base paired. That said, broadening of the U7 and G8 imino protons upon addition of **2** as observed in the 1D titration indicates that the complex is dynamic. In summary, the NMR data suggests that **2** adopts a similar binding mode to **1**, where the compound laid in the major groove of the RNA and between the closing GC pairs of r(CUG), but with a different conformation of the UU mismatch. As with **1**, each of the aromatic rings appeared to be in close proximity to one, but not both strands of the RNA.

**Figure 3:**
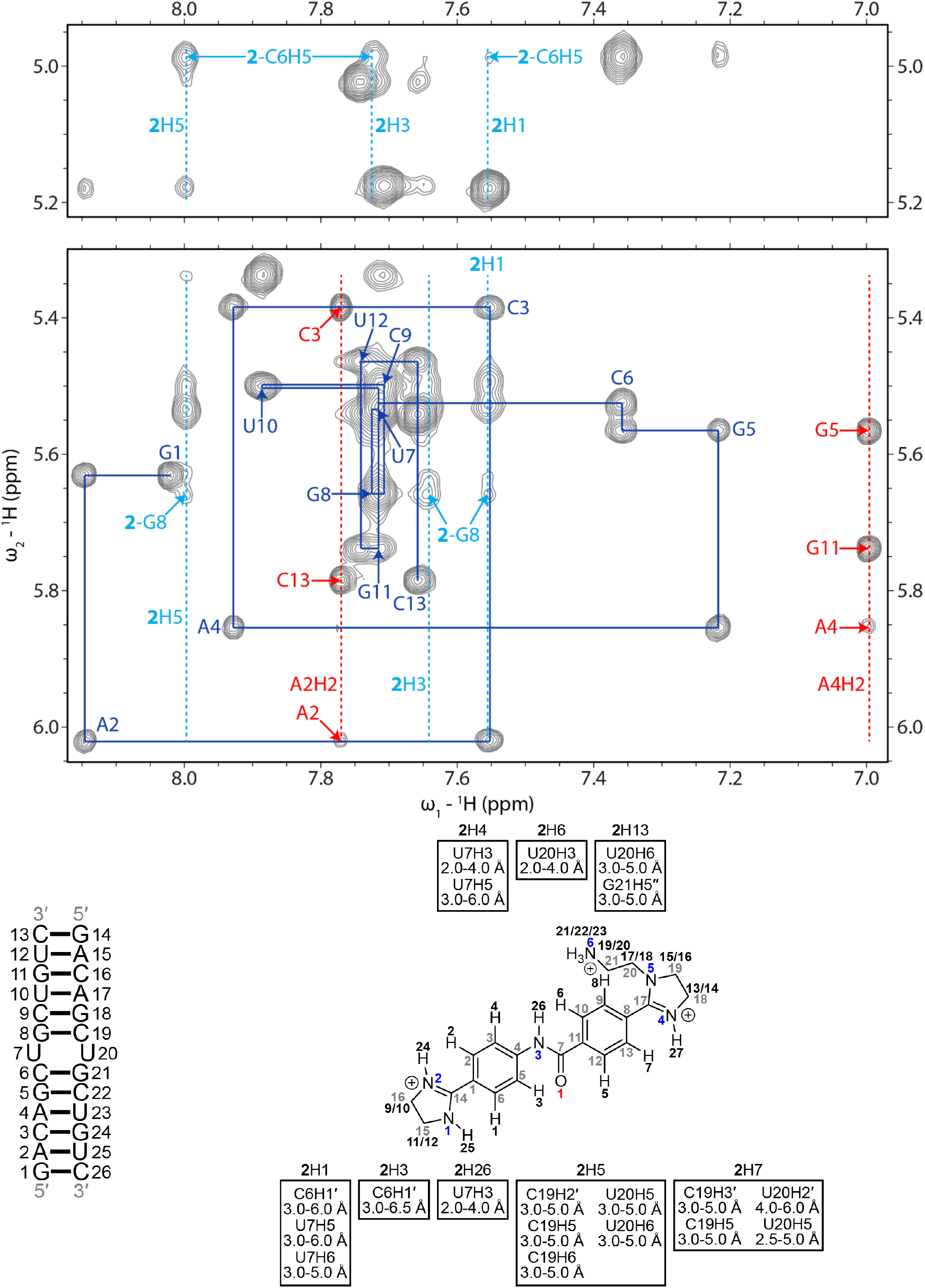
H6/H8-H1′ region of a 2D ^1^H-^1^H NOESY spectrum of the r(CUG)-2 complex. Blue lines represent a sequential H6/H8-H1′ walk, and blue labels represent intraresidue H6/H8 to H1′ NOEs. Dashed red lines represent adenine H2 resonances, and red labels represent interresidue and intraresidue NOEs between adenine H2 and H1′ of nearby residues. Dashed light blue lines represent aromatic proton resonances of **2**, and light blue labels correspond to NOEs between H1′of r(CUG) residues, unless otherwise noted, and **2**. RNA residues are only numbered from 1 to 13 in the NMR spectrum, and residues 14 to 26 correspond to residues 1 to 13 due to the symmetry of the RNA construct. In the chemical structure of **2**, black numbers represent hydrogens, gray numbers represent carbons, blue numbers represent nitrogens, and red numbers represent oxygen. NOEs between the RNA and **2** and their corresponding distance restraints used for modeling are colored according to the atoms of **2**. The spectrum was acquired at 35 °C with 400 ms mixing time and 0.3 mM of RNA and 0.6 mM of **2**.

### Structures of r(CUG) repeat-1 and r(CUG) repeat-2 complexes

As expected from the uninterrupted NOESY walk observed for both small molecules (**Figures 2** **– 3**), the r(CUG) RNA retained A-form geometry when bound to **1** or **2**. A total of 24 and 23 intermolecular NOEs were used as distance restraints to model the r(CUG)-**1** and -**2** complexes, respectively (**Tables 1, S4 – S6**). The average RMSD of all heavy atoms for the ensemble of structures from restrained MD was 1.81 ± 0.43 Å for r(CUG)-**1** and 1.87 ± 0.48 for r(CUG)-**2**, both slightly higher than that of the apo structure (1.25 ± 0.33 Å; **Table 1**).

Although small molecules **1** and **2** both laid in the major groove of the 5’CUG/3’GUC internal loop motif, different intermolecular interactions afford stability to the complexes (**Figures 1** and **4**). Small molecule **1** formed stacking interactions with the closing GC pair(s), but not the uracil loops. Both uracil loops were stacked in the helix in 19 of the 20 structures; in the remaining structure one uracil was flipped into the major groove and formed one hydrogen bond with a guanidine of **1**. However, the uracils were shifted away from each other in base pair plane, disrupting their interbase hydrogen bonding interactions. Additionally, they did not form contacts with **1**, consistent with the absence of NOEs between **1** and the uracils. The positively charged guanidines formed electrostatic interactions (5 Å distance or less)^66, 67^ with the phosphate backbone at G8 and G21. In all structures that comprise the ensemble formed by the r(CUG) repeat-**1** complex, the carbonyl oxygen of **1** was oriented towards the minor groove. The major and minor grooves within the 5’C<u>U</u>G/3’G<u>U</u>C motif were widened in the r(CUG)-**1** structures, and the loop’s closing GC pairs had the largest changes in buckle of the C6G21 (14.2 ± 6.7° vs. 5.8 ± 6.4° in the apo structure) and G8C19 pairs (−15.8 ± 7.1° vs. −4.2 ± 4.8° in the apo structure).

**Figure 4:**
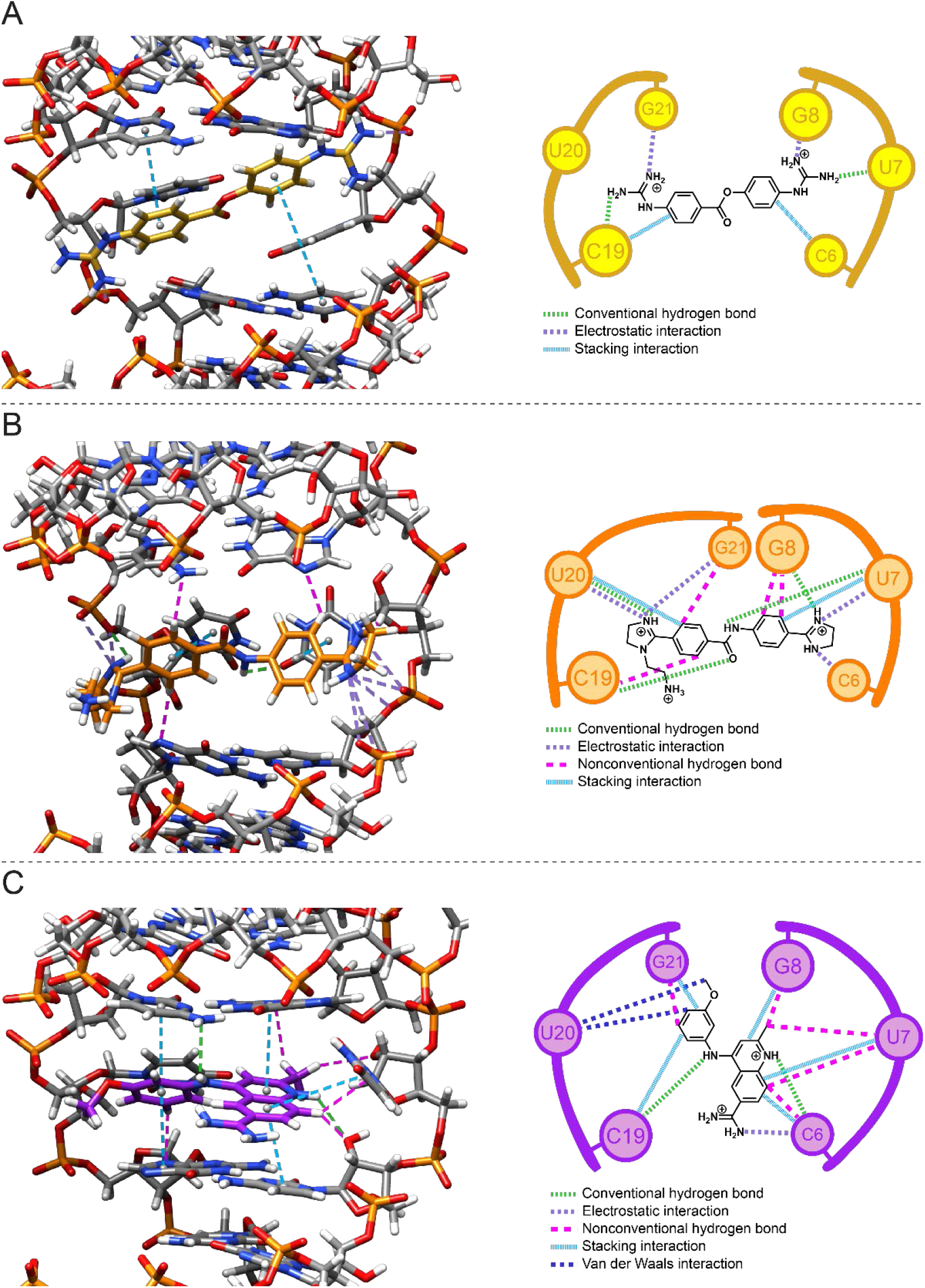
3D and schematic diagrams of ligand-r(CUG) interactions. (A) Interactions in the r(CUG)-**1** complex. (B) Interactions in the r(CUG)-**2** complex. (C) Interactions in the r(CUG)-**3** complex. In the 3D and schematic diagrams, light green dashed lines represent conventional hydrogen bonds, purple dashed lines represent electrostatic interactions, magenta dashed lines represent nonconventional hydrogen bonds, and light blue dashed lines represent stacking interactions. In the schematic diagrams, dark blue lines represent van der Waals interactions (not shown in the 3D diagrams).

Not surprisingly, addition of **1** to r(CUG) disrupted hydrogen bonding within the UU mismatch and resulted in increases in helical twist at 5′CU/3′GU and 5′UG/3′UC steps within the 5’C<u>U</u>G/3’G<u>U</u>C motif. At one of the 5′GC/3′CG steps, the helical twist increases from 31.7 ± 1.5° to 35.3 ± 3.8° while at the other 5′GC/3′CG step, the helical twist remains relatively unchanged (33.2 ± 11.7°) in the apo structure and 33.5 ± 3.1° in the **1**-bound structure). Changes in values of 5′GC/3′CG parameters also suggest that helical regions upstream of the UU mismatch undergo rearrangement to accommodate binding of the compound. The bend angle of the helix decreased from 11.88 ± 5.01° for the apo structure to 9.53 ± 4.26° for the **1**-bound structure. The major groove width at the 5′CU/3′GU and 5′UG/3′UC steps and flanking 5′GC/3′CG steps increase upon binding of **1**. These results suggest that the compound affects the helix away from the r(CUG) motif, consistent with shifting of imino proton resonances in base pairs away from the r(CUG) motif. Large variation in these values suggest that the r(CUG) motif in the apo and bound structures are dynamic and that binding of **1** disrupts localized interactions that stabilize the r(CUG) motif.

The overall average helical rise and twist of all nucleotides bound RNA were similar to that of the apo structure, 2.2 ± 1.0 Å and 31.8 ± 3.0° (2.8 ± 0.3 Å and 30.9 ± 2.0°, respectively, for the apo structure). However, the increased errors in these values as compared to those of apo- r(CUG) suggest that the r(CUG)-**1** complex is dynamic. An increase in the C1′-C1′ distance at the UU mismatch in the bound structure was also observed, as compared to the apo structure, 12.8 ± 0.7 Å and 10.0 ± 0.5 Å, respectively. Large propeller twist values were also observed for the UU pair in the r(CUG)-**1** complex and were significantly larger than observed in the unbound r(CUG) repeat structure. However, the values for shear (a measure of sliding of one base in relation to the other)^57^ were not significantly different between the two structures; rather, base non-planarity is represented in the stagger values of the r(CUG) repeat-**1** structures. Altogether, the r(CUG)-**1** complex is stabilized by stacking and hydrogen bonding interactions and remodels the RNA structure around the UU loop, by decreasing the bend angle of the helix, increasing the major groove width within the r(CUG) motif, disrupting hydrogen bonding in the UU mismatch, and lengthening the C1′-C1′ distance of the UU mismatch.

As mentioned above, the binding of **2** to the r(CUG) repeat did not disrupt the overall A-form geometry of the RNA, as evidenced by the average helical rise (2.8 ± 0.5 Å) and helical twist (30.6 ± 19.9°) for the ensemble of 20 r(CUG) repeat-**2** structures generated by restrained MD (**Figure 1**). Although these values are similar for the apo structure, the large error in helical twist suggests that the r(CUG)-**2** complex is dynamic. Although **2** also formed stacking interactions the 5’CUG/3’G<u>U</u>C internal loop motif as **1** did, **2** stacked on the mismatched uracils, rather than the GC closing base pair (**Figure 4**). In further contrast to **1**, in most of the r(CUG)-**2** structures (14/20), either one or both UU mismatch bases were not stacked in the helix and displaced towards the minor groove. However, the uracils remained base paired, as evident from an average of 1.6 ± 0.5 hydrogen bonds vs. 0.2 ± 0.4 for the U7U20 pair in the **1**-bound structure. The mismatched uracils were also in close proximity to **2**, consistent with NOEs between U7/U20 imino proton and **2**. The interactions between **2** and the UU loop were also stabilized by a hydrogen bond (3 Å distance or less) between an uracil of 5’C<u>U</u>G/3’G<u>U</u>C internal loop and the amide H26 of the small molecule (12/20 structures) and a hydrogen bond between the phosphate backbone at U20 and H27 of **2** (16/20 structures). The imidazolines form electrostatic interactions with the phosphate backbone at C6 in 10 of the 20 structures, at U7 in all of the structures, at U20 in 19 of the 20 structures, and at G21 in six of the 20 structures. Aromatic protons of the benzylimidazoles formed van der Waals and/or nonconventional hydrogen bonding interactions with G8, C19, and/or G21G21.

A closer inspection of the structure of the internal loop bound to **2** indicates that the 5′CU/3′GU step is under-twisted and the 5′UG/3′UC step is over-twisted, concomitant with a much shorter C1′-C1′ distance, 9.6 ± 1.0 Å. Farther away from the r(CUG) motif, the 5′GC/3′CG steps appear to be over-twisted. This under- and over-twisting and the shortening of C1′-C1′ distance is more severe in the r(CUG)-**2** complex than that observed in both apo structure or in the structure of **1** complex. Akin to that observed in the structure of the r(CUG) repeat-**1** complex, a large propeller twist value, 25.8 ± 29.0°, was observed for the UU pair in the r(CUG) repeat-**2** structure vs. −9.8 ± 3.6° in the unbound structure while the shear values were not significantly different. Rather, stagger values for r(CUG) repeat-**2** structures (−1.1 ± 1.8° for C6G21) indicated non-planarity of the RNA bases. Collectively, these changes upon **2** binding of the internal loop slightly widened. The large error values in the parameters for the **2**-bound structure indicate that the complex is dynamic as the helix accommodates binding of the compound. The dynamics observed in the r(CUG) motif are also evident from breathing of the closing GC pairs, which have average numbers of hydrogen bonds of 2.2 ± 1.3 for the C6-G21 and 2.8 ± 0.5 for the G8-C19 pair. The changes in the structure of r(CUG) are also consistent with shifting of imino resonances upon titration of **2** into r(CUG). In summary, the binding of **2** to r(CUG) repeat distorts the RNA helix by increasing the bend angle by 2.68° (average of 14.81 ± 2.68°) and compensates for those distortions by forming intermolecular stacking and hydrogen bonds.

### Comparison of the r(CUG)-1 and r(CUG)-2 structures

The Carloni group predicted the structure of **1** bound to r(CAG) using docking and molecular dynamics simulations.^68^ In their structures, **1** binds to the major groove of r(CAG) and interacts with both strands of the RNA. The hydrogen bonding interactions of one of the AA pairs (A7-A14) appears to be altered from the free state when bound to **1**. In the r(CUG)-**1** and -**2** NMR structures, the compounds also bind in the major groove, but **1** disrupted the hydrogen bonding interactions in the UU mismatch whereas **2** did not. In the Carloni group’s structure of **1** bound to a r(CAG) repeat, hydrogen bonds were observed between the guanidines of **1** and the phosphate backbone of the RNA. In the r(CUG)-**1** and r(CUG)-**2** structures, the guanidines of **1** and imidazolines of **2** also formed hydrogen bonds and electrostatic interactions with the negatively charged phosphates. The modeled r(CAG)-**1** structures have stacking interactions between the rings of **1** and the mismatched bases of the 1×1 AA pairs. In comparison, stacking interactions between the compound and mismatched uracils were observed in the r(CUG)-**2** but not -**1** structures. To facilitate these stacking interactions, the rings in **2** adopted a quasi-perpendicular orientation in some and a planar orientation in the other states. The quasi-perpendicular orientation of the rings in **1** was present in the Carloni group’s r(CAG)-**1** structures, but not the NMR structures of r(CUG)-**1**, where the rings adopted a planar orientation in all states. The mismatched uracils in the r(CUG)-**1** structures formed stacking interactions with the helix, but lost their interbase hydrogen bonding, whereas those in the r(CUG)-**2** structures retained interbase hydrogen bonding but lost their stacking interactions with the helix. The mismatched A4-A17 and A7-A14 bases in the Carloni group’s structures and the U7-U20 bases in the r(CUG)-**1** structures had negative opening values, but the buckle values of both closing GC pairs were negative in the r(CAG)-**1** structure and alternated between negative and positive in the r(CUG)-**1** structures. In the r(CUG)-**2** structures, the opening was positive and the average buckle values did not change significantly from apo-r(CUG). The major groove widened upon binding of compound in the Carloni group’s work and in the r(CUG)- **1** and -**2** structures. Altogether, binding of **1** and **2** to r(CUG) and r(CAG) induces changes in the local structure of the RNAs but differentially affects interactions between the mismatched bases. Furthermore, selectivity of the compounds for r(CUG) motifs can be increased by designing the compounds to interact with the UU mismatches and closing GC pairs.

### NMR analysis of r(CUG) bound to 3

As observed for **1** and **2**, binding of **3** did not disrupt a NOESY walk through the r(CUG) repeat duplex model (**Figure 5**), indicating that an overall A-form geometry was adopted. In total, 32 NOEs were observed from **3** to the RNA (**Figure 5** **and Table S11**), more than that observed for **1** (24) or **2** (23). NOEs were also observed between H5, H6, and H8 of **3** and the RNA, suggesting that the methoxybenzene group rotates about the C10-N3 bond upon binding to the RNA (**Figures 5** and **S13**). Of note, the NOEs between **3**’s H5 and the r(CUG) motif were not used as restraints for modeling because they were close to the level of noise in the 100 ms mixing time NOESY spectrum. The average RMSD for the 20 structures with the fewest distance restraint violations was 1.02 ± 0.20 Å (**Table 1**).

**Figure 5:**
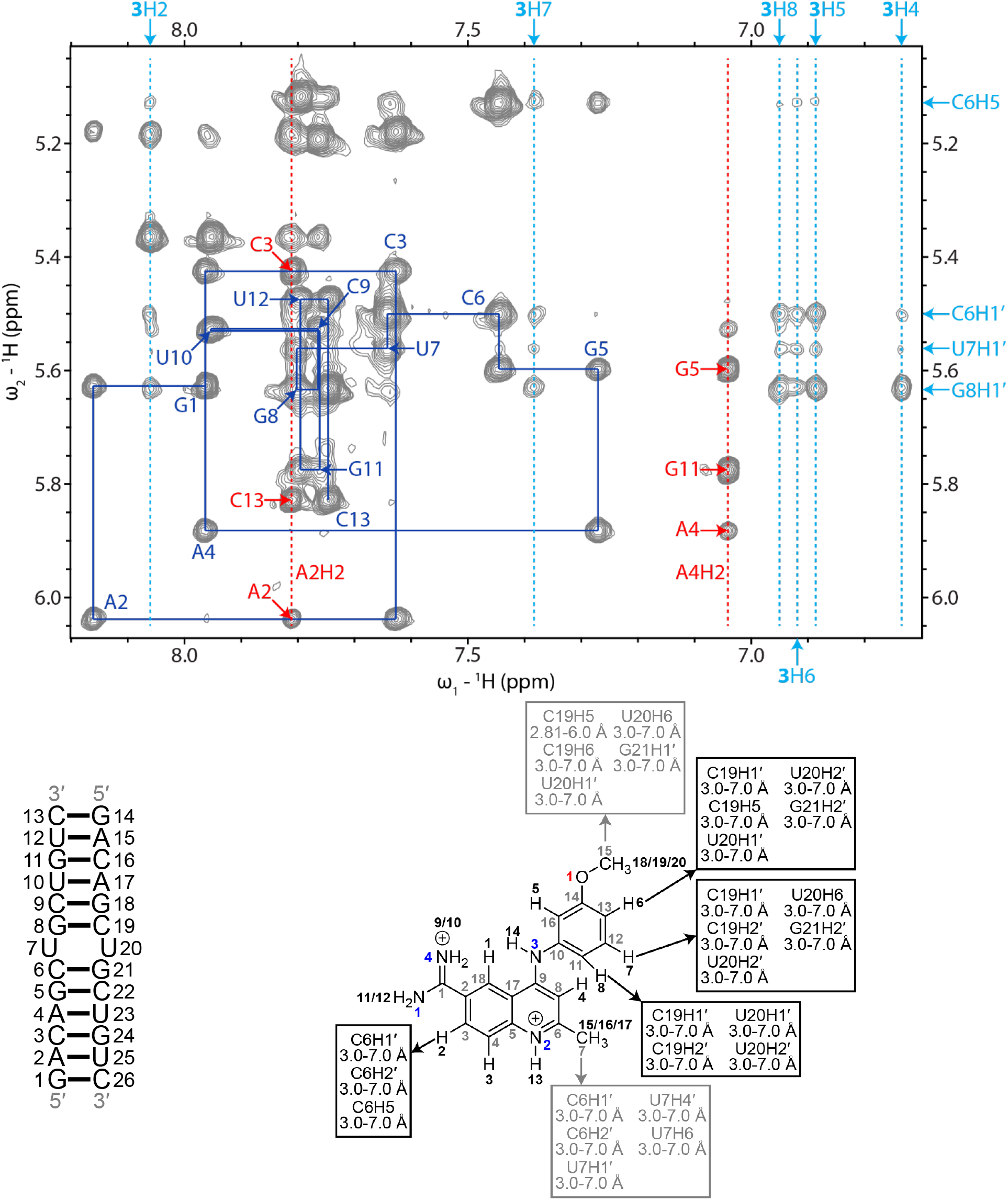
H6/H8-H1′ region of a 2D ^1^H-^1^H NOESY spectrum of the r(CUG)-3 complex. Blue lines represent a sequential H6/H8-H1′ walk, and blue labels represent intraresidue H6/H8 to H1′ NOEs. Dashed red lines represent adenine H2 resonances, and red labels represent interresidue and intraresidue NOEs between adenine H2 and H1′ of nearby residues. Dashed light blue lines represent aromatic proton resonances of **3**. Light blue labels around the edges of the spectrum correspond to r(CUG) and aromatic hydrogens of **3** in RNA-**3** NOEs. RNA residues are only numbered from 1 to 13 in the NMR spectrum, and residues 14 to 26 correspond to residues 1 to 13 due to the symmetry of the RNA construct. In the chemical structure of **3**, black numbers represent hydrogens, gray numbers represent carbons, blue numbers represent nitrogens, and red numbers represent oxygen. NOEs between the RNA and **3** and their corresponding distance restraints used for modeling are colored according to the atoms of **3**. The spectrum was acquired at 35 °C with 400 ms mixing time and 0.4 mM of RNA and 0.6 mM of **3**.

### Structure of the r(CUG) repeat-3 complex

As aforementioned, the overall r(CUG) repeat-**3** complex formed an overall A-form geometry as confirmed by its average helical rise and twist for the ensemble structures, 2.5 ± 0.6 Å and 32.8 ± 11.1°, respectively (**Figure 1**). The large error in helical twist suggests that the complex is dynamic. Perhaps unexpectedly, the binding mode for **3** and its subsequent effect on the RNA’s structure is markedly different than that of **1** and **2**, where stacking interactions, hydrogen bonds, and van der Waals interactions stabilize the r(CUG)-**3** complex (**Figure 4**). The quinoline and methoxybenzene groups of **3** stacked between the closing GC pairs of the 5’C<u>U</u>G/3’G<u>U</u>C motif and displaced one of the uridines of the UU mismatch into the major groove and the other into the minor groove. (Small molecule **1** also stacked on the GC pairs.) The C7 methyl group was oriented towards the minor groove while the C15 methyl group pointed towards the phosphate backbone of the RNA. These groups formed non-conventional hydrogen bonds / van der Waals interactions with the RNA. These types of interactions between the small molecule’s H3 and H7 and the loop nucleotides were also common in the ensemble of 20 r(CUG)-**3** structures. H13 and H14 of the compound formed hydrogen bonds with cytidines forming the closing GC pairs on the opposite strand of the RNA and the amidine lied inside the major groove and forms electrostatic interactions with the phosphate backbone at C6. Taken together, the r(CUG)-**3** complex is stabilized by a combination of stacking, van der Waals, and hydrogen bonding interactions.

Although the overall average twisting of the r(CUG) repeat-**3** complex indicated A-form geometry, the helix appears to be under-twisted at the 5′CU/3′GU step and over-twisted at the 5′UG/3′UC step. At the 5′GC/3′CG steps adjacent to the 5’C<u>U</u>G/3’G<u>U</u>C internal loop, the helix is over-twisted (39.8 ± 2.1° and 35.2 ± 0.9°) while the bend angle of the helix increased to 13.92 ± 5.06°. To accommodate **3**, the major groove was slightly narrower at the 5′CU/3′GU and 5′UG/3′UC steps (17.8 ± 0.9 Å and 17.7 ± 0.9 Å, respectively) and at one of the 5′GC/3′CG steps (17.4 ± 0.7 Å) than that observed for the same steps) in the apo structure (19.6 ± 1.2 Å for 5′CU/3′GU and 5′UG/3′UC steps and 19.6 ± 1.0 Å for the same 5′GC/3′CG step). The changes in the 5′GC/3′CG steps were consistent with shifts in G5H1 and G8H1 resonances in the titration experiment. An increase in C1′-C1′ distance from 10.0 ± 0.5 Å in the unbound structure to 14.6 ± 0.2 Å disrupted the UU base pair and allowed the compound to stack between the closing GC pairs. Nonplanarity of the UU mismatch was also evident from high values for parameters such as buckle (22.5 ± 60.3°), shear (14.5 ± 0.8°) and stagger (4.8 ± 1.5°) for the UU pair, and changes in rise at the 5′CU/3′GU (5.9 ± 0.9 Å vs. 3.7 ± 0.2 Å in apo structure) and 5′UG/3′UC steps (−0.8 ± 1.5 Å vs. 3.7 ± 0.2 Å in the apo structure), roll at the 5′CU/3′GU (−41.5 ± 21.3° vs. 9.2 ± 3.8° in the apo structure) and 5′UG/3′UC steps (13.4 ± 7.5° vs. 7.3 ± 2.5° in the apo structure), and tilt at the 5′CU/3′GU (−73.4 ± 39.5° vs. 0.6 ± 3.4° in the apo structure) and 5′UG/3′UC steps (83.3 ± 43.7° vs. −2.7 ± 3.4° in the apo structure). The large errors in values for the base pair parameters are consistent with dynamics observed in the ensemble of r(CUG)-**3** structures. In summary, binding of **3** to r(CUG) brings about structural changes and dynamics in the helix.

### Comparison of the r(CUG)-3 structure with tau RNA-3 structure

In the r(CUG)-**3** structures and tau RNA-**3** structures, **3** lied in the major groove of the RNA and stacks with the RNA via its quinoline group. The C7 methyl group in these structures was positioned between the closing GC pairs of the A-bulge or UU mismatch and forms van der Waals and/or nonconventional hydrogen bonds with the closing GC pair(s). Similarly, the C15 group interacted with the RNA backbone in the complexes. Compared to the tau RNA-**3** structures, the quinoline core in the r(CUG)-**3** structures is turned slightly towards one of the RNA strands. This orients the amidine away from the RNA backbone of one of the RNA strands and positions the N3 amino group closer to the opposite strand and facilitates a hydrogen bond with the C19 base. Taken together, the r(CUG)-**3** complex resembles the tau RNA-**3** complex and is stabilized by a similar set of interactions.

### Comparison of the r(CUG) repeat structures bound to 3 vs. 1 and 2

The r(CUG)-**3** complex, like r(CUG)-**1** and r(CUG)-**2** complexes, contained the compound in the major groove between the closing GC pairs, with the helices retaining A-form geometry. Whereas **1** and **2** formed stacking interactions with either of the closing GC pairs or the UU mismatch, **3** stacked with both of the GC pairs. To allow for this, the helix was bent to a greater degree in the r(CUG)- **3** structures than the apo or r(CUG)-**1** structures, although to a similar degree as the r(CUG)-**2** structures as a result of stacking of the extrahelical UU pair in most of the r(CUG)-**2** conformations. That said, one of the uracils in the r(CUG)-**3** structures remained in the helix and stacked on the compound while the other was flipped into the major groove. As a result, the C1′-C1′ distance of the mismatched uracils in the r(CUG)-**3** was greater than in r(CUG)-**1** and r(CUG)-**2** structures. Compound **3** also contained methyl groups positioned near the RNA backbone for weak nonconventional hydrogen bonds or van der Waals interactions with the RNA. In comparison, the imidazolines of **2** laid farther in the major groove and did not interact with the RNA. Instead, **2** formed a more extensive hydrogen bonding network with the RNA than **1** or **3**. Altogether, **3** appears to form more stabilizing interactions with r(CUG) than **1** or **2**. In summary, compounds **1**, **2**, and **3** form diverse sets of interactions that aid molecular recognition of r(CUG) repeats.).

### Implications & Conclusions

Structure-based drug design is powerful tool for drug discovery as evidenced by various efforts to identify and optimize small molecules that bind disease-causing proteins.^69^ Unfortunately, such methods are only in their infancy for RNA targets. Herein, we used the physicochemical properties of known small molecules that bind RNA, coupled with solubility measurements to identify three small molecules that bind r(CUG) repeats, amenable to structure determination by NMR spectrometry. NMR spectrometry, has several advantages over other structure determination techniques such as X-ray crystallography and cryo-EM,^70^ particularly its ability to capture conformational dynamics of the target and target-ligand complexes. Indeed, conformational dynamics have been incorporated in a virtual screen to find small molecules that bind HIV TAR RNA.^23^

This study provides the very first examples of small molecules bound to the UU internal loops found in r(CUG)^exp^. Compound **1** was previously identified to bind to 1×1 AA loops and inhibit MBNL1 sequestration in HD.^44^ For this reason, it was used to initiate structural studies of r(CUG)^exp^-binding compounds that inhibit MBNL1 sequestration in HDL2. Using the NMR structure of r(CUG)-**1**, **2** was designed with improved chemical properties and a structure of the compound in complex with r(CUG) was solved. Separately, an NMR structure was determined for r(CUG) in complex with **3**, which was previously identified to bind to an A-bulge and inhibit U1 snRNP binding in FTDP-17.^48^ These compounds each bound to the r(CUG) internal loop via unique sets of interactions. Thus, a structure-based approach to identify RNA-binding compounds may be initiated by searching for compounds that bind to RNA motifs, testing their solubility under NMR conditions, and then analyzing chemical shift perturbations of the RNA upon binding of the compounds.

The selectivity of **1** – **3** for r(CUG)^exp^ could be improved by optimizing their interactions with the 5’C<u>U</u>G/3’G<u>U</u>C internal loop, such as by eliminating steric hindrance or adding new hydrogen bonding and van der Waals interactions. This approach has been used to optimize small molecule drugs that target proteins.^71^ Compounds that selectively bind to RNA can also be identified by docking libraries of compounds to ensembles of RNA conformations generated by NMR studies and computational modeling.^23, 72^ Using ligand-bound receptors usually affords superior results than apo receptors in molecular docking because the binding sites are better defined in the bound structures than unbound ones.^73^ Thus, the results of this work provide invaluable information for future virtual screening campaigns against compounds that target r(CUG)^exp^. These data can also be used for machine learning based methods to improve the binding affinity as well as physicochemical properties through approaches like generative artificial intelligence (AI).^74^ These structures also provide the opportunity to adopt structure-based machine learning assisted drug design approaches as already exemplified in the protein-small molecule field.^75, 76^

## SUPPORTING INFORMATION AVAILABLE

(I) TR-FRET analysis of the inhibition of the r(CUG)^exp^-MBNL1 complex by compounds **1**, **2**, and **3**. (II) H6/H8-H1′ and imino proton regions of 2D ^1^H NOESY spectra of unbound r(CUG). (III) Structures of the unbound r(CUG) motif. (IV) 1D ^1^H and WaterLOGSY NMR spectra of **1**, **2**, and **3** alone and in complex with r(CUG). (V) 1D ^1^H imino region titrations with **1**, **2**, and **3**. (VI) Imino proton regions of 2D ^1^H NOESY spectra of the r(CUG)-**1**, -**2**, and -**3** complexes. (VII) Physicochemical properties of **1**, **2** and **3**. (VIII) NOE and dihedral restraints used for modeling of the unbound r(CUG) duplex. (IX) NOE restraints used for modeling of the r(CUG)-**1**, -**2**, and - **3** complexes. (X) Distance restraint violations greater than 0.1 Å for the r(CUG)-**1** complex. (XI) ^1^H NMR chemical shifts of the unbound r(CUG) and r(CUG)-**1**, -**2**, and -**3** complexes. (XII) Synthetic methods for compounds **1** and **2**.

## ABBREVIATIONS

1D, one-dimensional; 2D, two-dimensional; ALS, amyotrophic lateral sclerosis; ASO, antisense oligonucleotide; CRISPR, clustered regularly interspaced palindromic repeats; CUGBP1, CUG- binding protein 1; DM1, myotonic dystrophy type 1; DMPK, dystrophia myotonica protein kinase; FTDP-17, frontotemporal dementia with parkinsonism-17; HD, Huntington’s disease; HDL2, Huntington’s disease-like 2; IR, insulin receptor; MBNL1, muscleblind-like 1 protein; MD, molecular dynamics; nc, noncoding; NMR, nuclear magnetic resonance; NOE, nuclear Overhauser effect; NOESY, nuclear Overhauser effect spectroscopy; RBP, RNA-binding protein; RMSD, root-mean-square deviation; SCA, spinocerebellar ataxia; SRSF6, serine and arginine rich splicing factor 6; UTR, untranslated region.

## ACKNOWLEDGMENTS

This work was funded by the National Institutes of Health (R35 NS116846 to M.D.D), the Department of Defense (HT94252310336 to M.D.D), the Muscular Dystrophy Association (MDA 1069959 to M.D.D and Development Grant 22 grant 963835 to A.T.), the Huntington’s Disease Society of America (to J.L.C.). This study made use of NMR spectrometers at The Scripps Research Institute, Campus Chemical Instrument Center NMR facility at Ohio State University, Minnesota NMR Center, and the National Magnetic Resonance Facility at Madison (NMRFAM). NMR instrumentation at The Scripps Research Institute was purchased with funds from the NIH S10-OD021550. NMR instrumentation, helium recovery equipment, and computers at NMRFAM were purchased with funds from the University of Wisconsin-Madison, the NIH P41GM136463, R24 GM141526, P41 GM103399, S10 RR023438, S10 RR025062, and S10 RR029220.

