## Supplementary File 1 for "NMR structures of small molecules bound to a model of an RNA CUG repeat expansion"

### Contents Page


| | | |
|---|---|---|
| **Supplementary Figures** | | 4 |
| Figure S1 | In vitro inhibition of the r(CUG)$^{exp}$-MBNL1 complex by compounds **1**, **2**, and **3**, as determined by a TR-FRET assay | 4 |
| Figure S2 | H6/H8-H1′ region of a 2D $^{1}$H NOESY spectrum of a model of apo r(CUG)$^{exp}$ containing a single 5'CUG/3'GUC internal loop | 5 |
| Figure S3 | Imino proton region of a 2D $^{1}$H NOESY spectrum of unbound r(CUG) | 6 |
| Figure S4 | Structures of the unbound r(CUG) motif from restrained MD simulations | 7 |
| Figure S5 | 1D $^{1}$H and WaterLOGSY NMR spectra of **1** alone and in complex with r(CUG) | 8 |
| Figure S6 | 1D $^{1}$H and WaterLOGSY NMR spectra of **2** alone and in complex with r(CUG) | 9 |
| Figure S7 | 1D $^{1}$H and WaterLOGSY NMR spectra of **3** alone and in complex with r(CUG) | 10 |
| Figure S8 | 1D $^{1}$H imino region titration of r(CUG) in the presence of **1** | 11 |
| Figure S9 | 1D 1H imino region titration of r(CUG) in the presence of **2** | 12 |
| Figure S10 | 1D 1H imino region titration of r(CUG) in the presence of **3** | 13 |
| Figure S11 | Imino proton region of a 2D $^{1}$H NOESY spectrum of the r(CUG)-**1** complex | 14 |
| Figure S12 | Imino proton region of a 2D $^{1}$H NOESY spectrum of the r(CUG)-**2** complex | 15 |
| Figure S13 | Imino proton region of a 2D $^{1}$H NOESY spectrum of the r(CUG)-**3** complex | 16 |
| **Supplementary Tables** | | 17 |
| Table S1 | Physicochemical properties of **1**, **2** and **3** | 17 |
| Table S2 | NOE restraints used for modeling of the unbound r(CUG) duplex | 18 |
| Table S3 | Dihedral restraints used for modeling of the unbound r(CUG) duplex | 23 |
| Table S4 | NOE restraints used for modeling of the r(CUG)-**1** complex | 26 |

| | | |
|---|---|---|
| Table S5 | Distance restraint violations greater than 0.1 Å for the r(CUG)-**1** complex | 33 |
| Table S6 | NOE restraints used for modeling of the r(CUG)-**2** complex | 34 |
| Table S7 | NOE restraints used for modeling of the r(CUG)-**3** complex | 40 |
| Table S8 | $^{1}$H NMR chemical shifts of the unbound r(CUG) duplex | 48 |
| Table S9 | $^{1}$H NMR chemical shifts of r(CUG)-**1** complex | 49 |
| Table S10 | $^{1}$H NMR chemical shifts of r(CUG)-**2** complex | 50 |
| Table S11 | $^{1}$H NMR chemical shifts of r(CUG)-**3** complex | 51 |

**Synthetic methods for compounds 1 and 2** 52

**References**

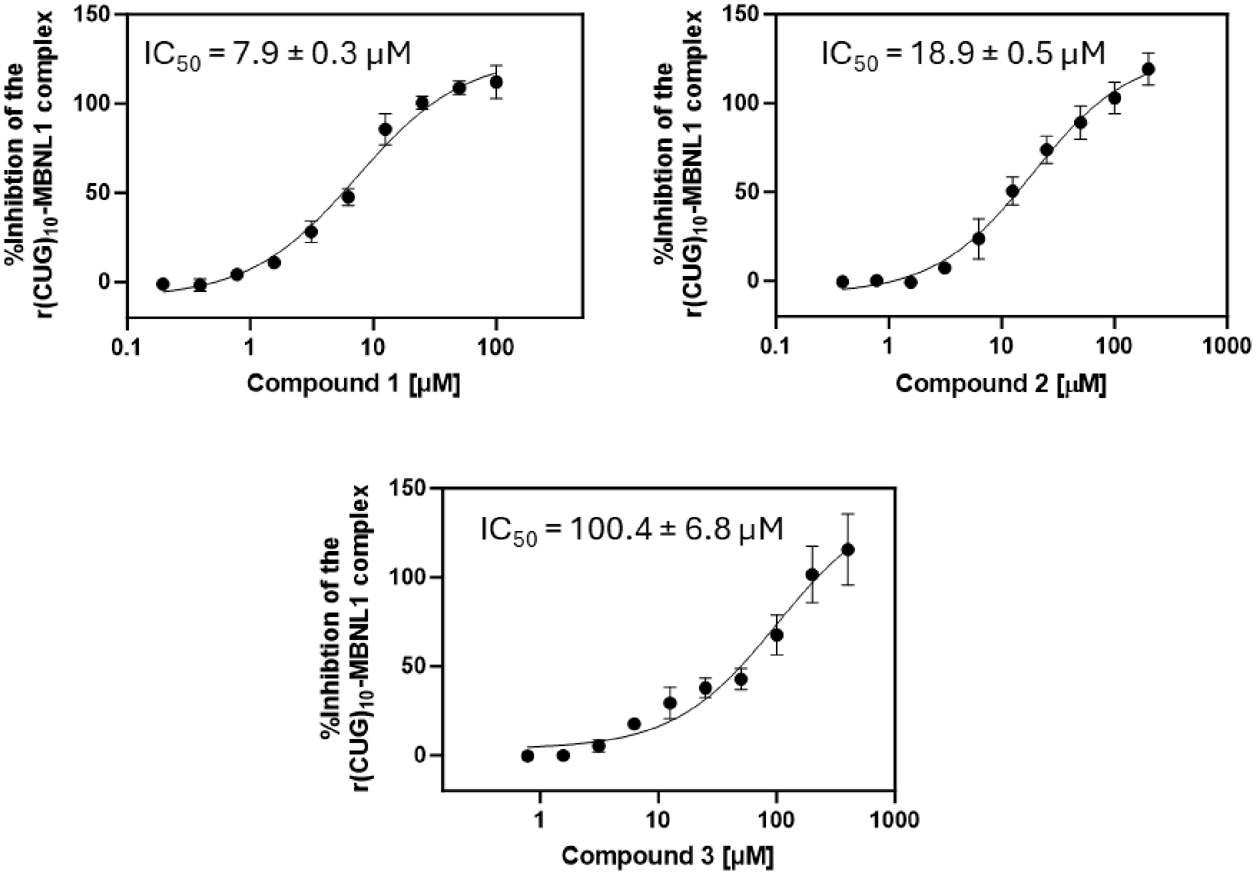


**Figure S1**: **In vitro inhibition of the r(CUG)$^{exp}$-MBNL1 complex by compounds 1, 2, and 3.** A previously reported time-resolved fluorescence resonance energy transfer (FRET) assay[1] was used to measure the percent inhibition of a r(CUG)$_{12}$-MBNL1 complex to generate $IC_{50}$ curves.

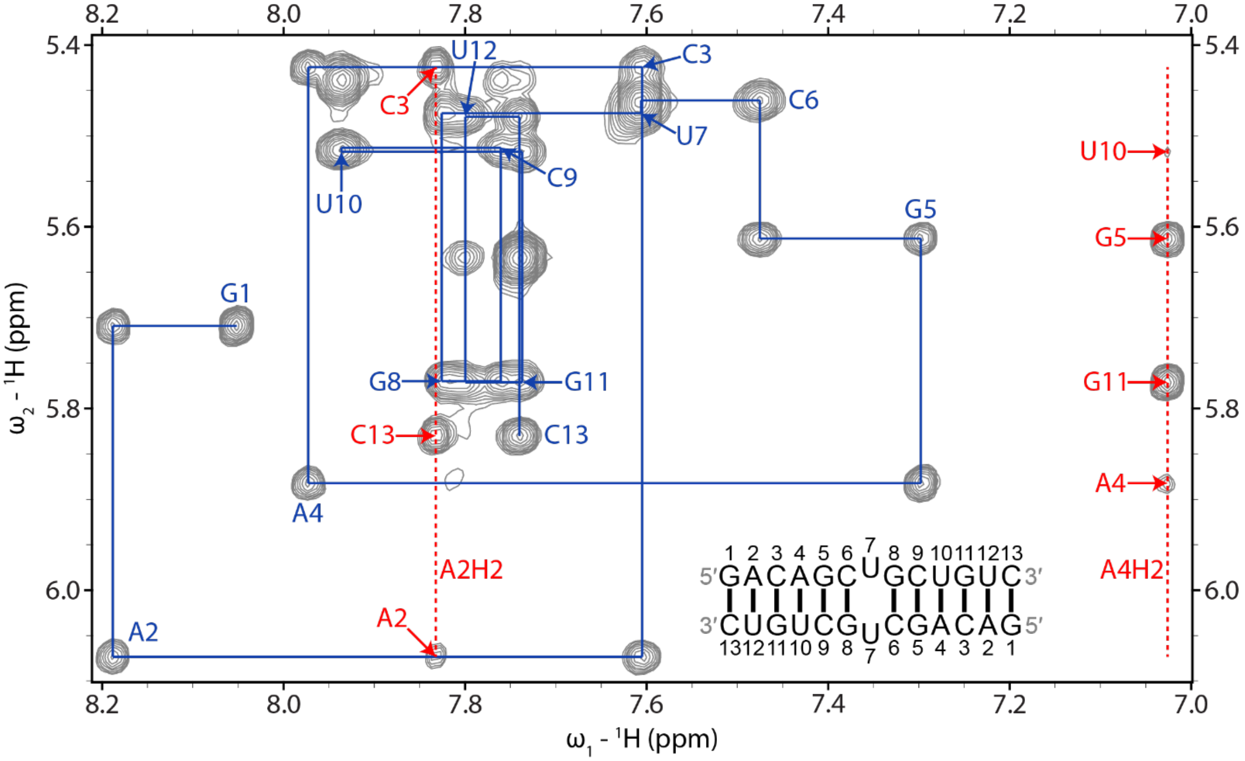


**Figure S2: H6/H8-H1′ region of a 2D $^{1}H$ NOESY spectrum of unbound r(CUG).** Blue lines represent a sequential H6/H8-H1′ walk, and blue labels represent intraresidue H6/H8 to H1′ NOEs. Dashed red lines represent adenine H2 resonances, and red labels represent interresidue and intraresidue NOEs between adenine H2 and H1′ of nearby residues. The spectrum was acquired at 25 °C with 400 ms mixing time and 0.3 mM of RNA.

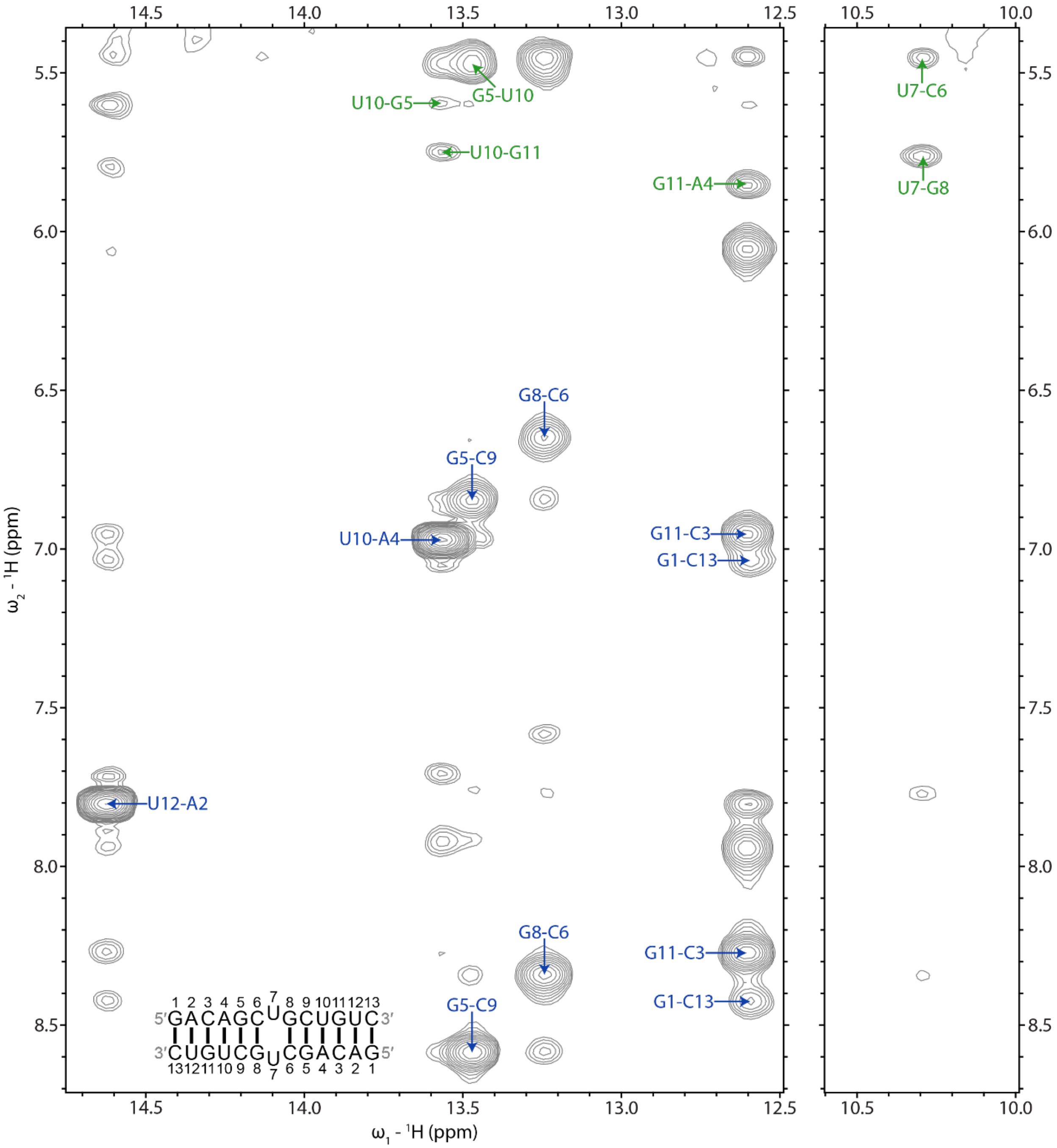


**Figure S3: Imino proton region of a 2D $^1$H NOESY spectrum of unbound r(CUG).** Blue labels represent UH3-AH2 and GH1 to C amino NOEs within base pairs. Green labels represent NOEs between UH3 or GH1 and the H1′ of a 3′ adjacent or cross-strand residue. In each label, the first residue is UH3 or GH1 and the second label is adenine H2, a cytosine amino proton, or H1′ of any residue. The spectrum was acquired at 5 °C with 125 ms mixing time and 0.3 mM of RNA.

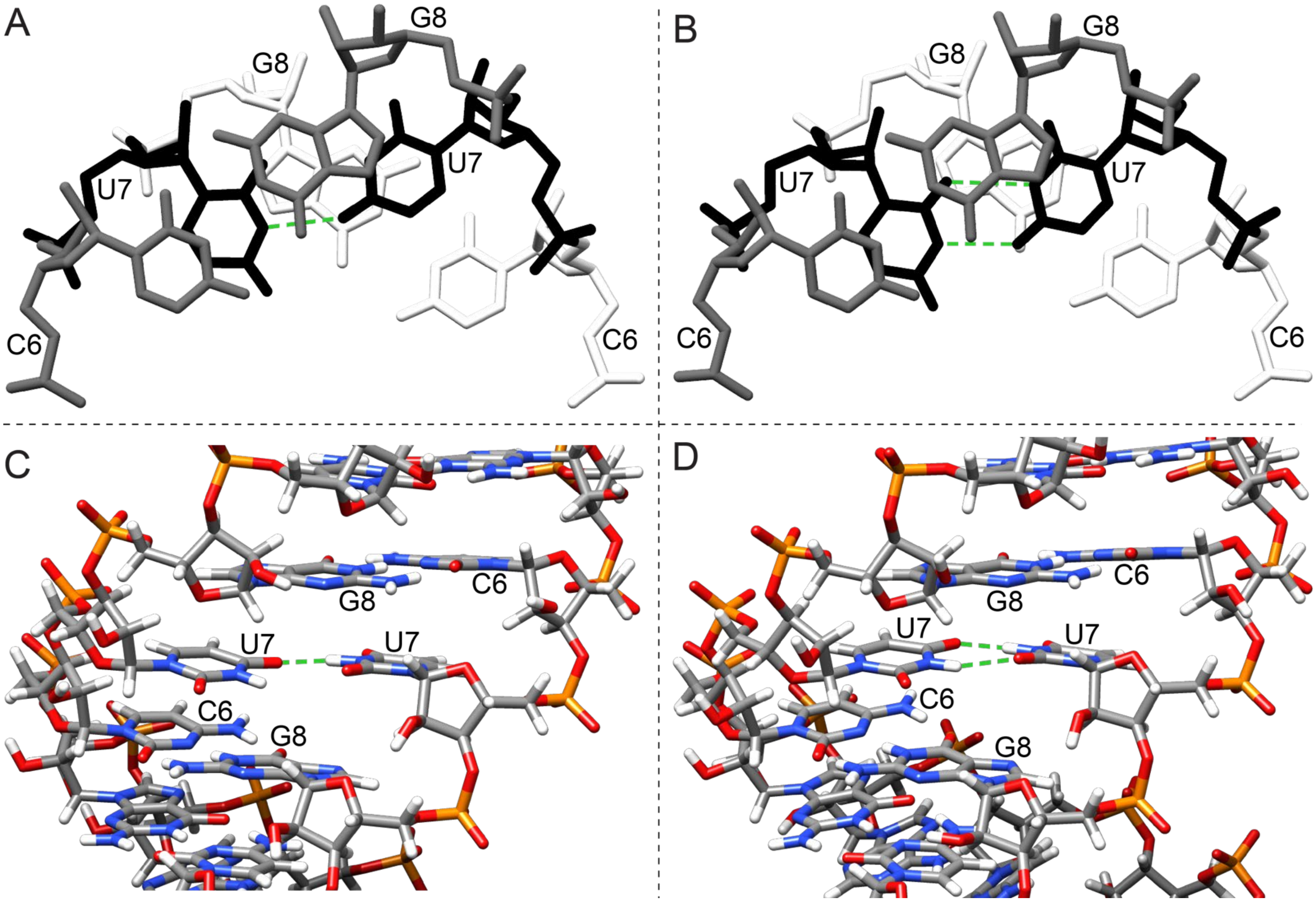


**Figure S4: Structures of the unbound r(CUG) motif (5'CUG/3'GUC).** (A) Major groove view showing overtwisting and undertwisting of the helix at the r(CUG) motif containing one and two hydrogen bond UU pairs where the UU pairs are colored black. (B) Minor groove view showing the stacking of one and two hydrogen bond UU pairs in the helix. Hydrogen bonds in the UU pairs are shown with green dashed lines.

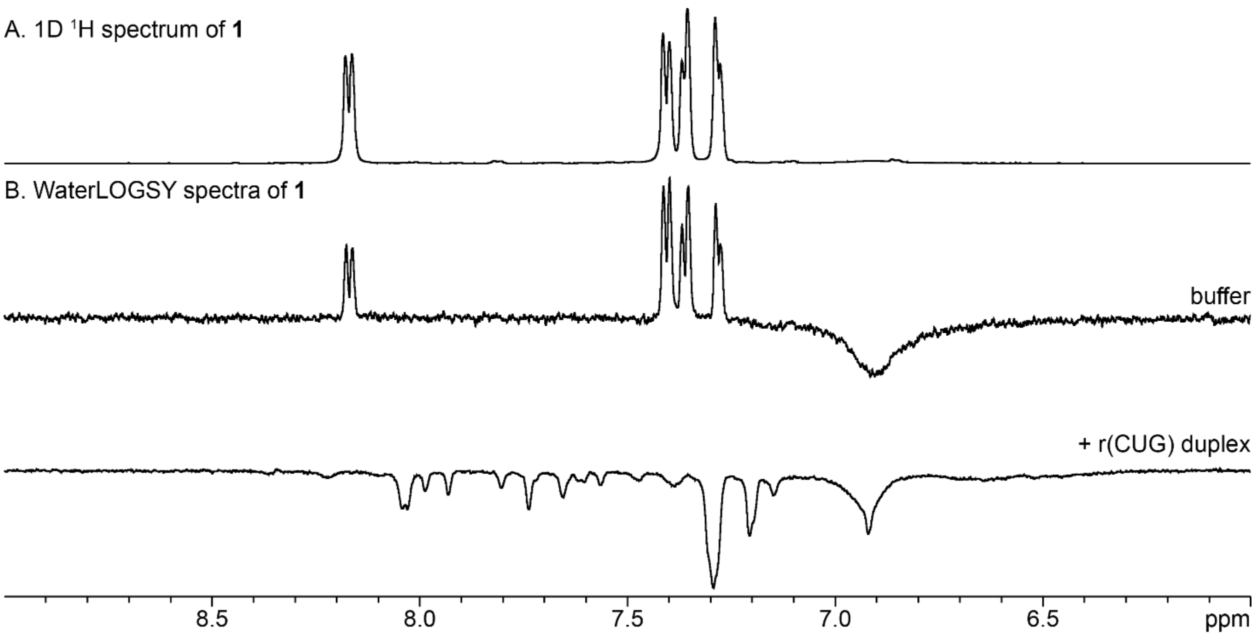


**Figure S5: 1D $^{1}$H and WaterLOGSY NMR spectra of 1 alone and in complex with the model r(CUG) duplex.** (A) 1D $^{1}$H NMR spectrum of 300 μM of **1**. (B) WaterLOGSY spectra of 300 μM of **1** in buffer alone or with 10 μM of the model r(CUG) duplex. Spectra were acquired at 25 °C.

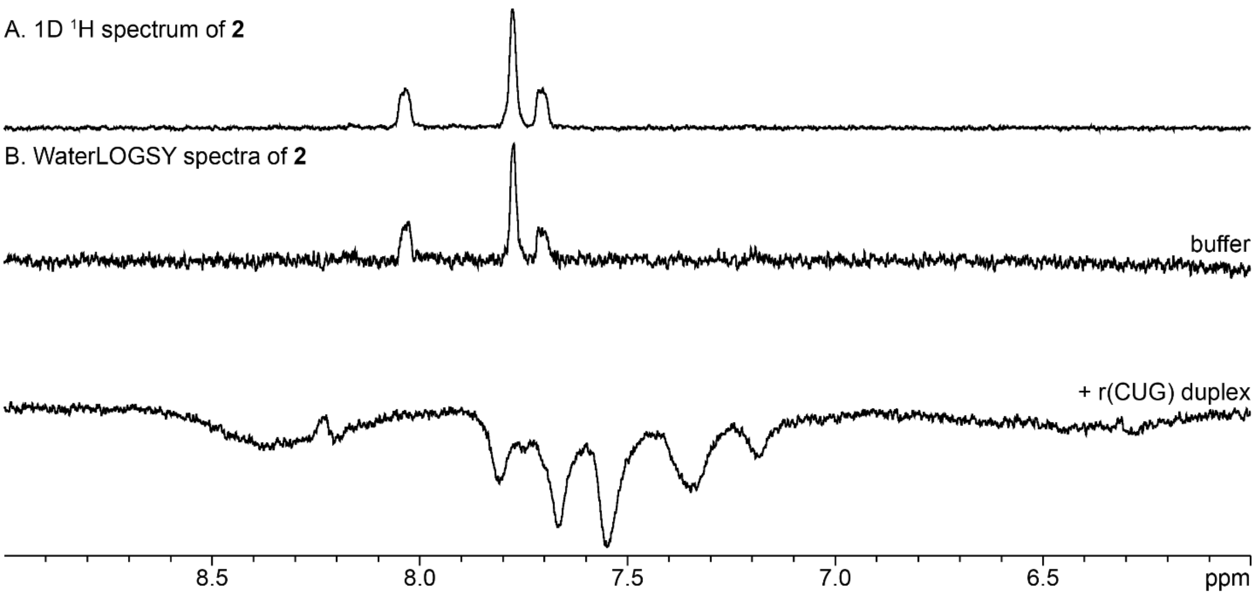


**Figure S6: 1D $^1$H and WaterLOGSY NMR spectra of 2 alone and in complex with the r(CUG) duplex.** (A) 1D $^1$H NMR spectrum of 300 μM of **2**. (B) WaterLOGSY spectra of 300 μM of **2** in buffer alone or with 10 μM of the model r(CUG) duplex. Spectra were acquired at 25 °C.

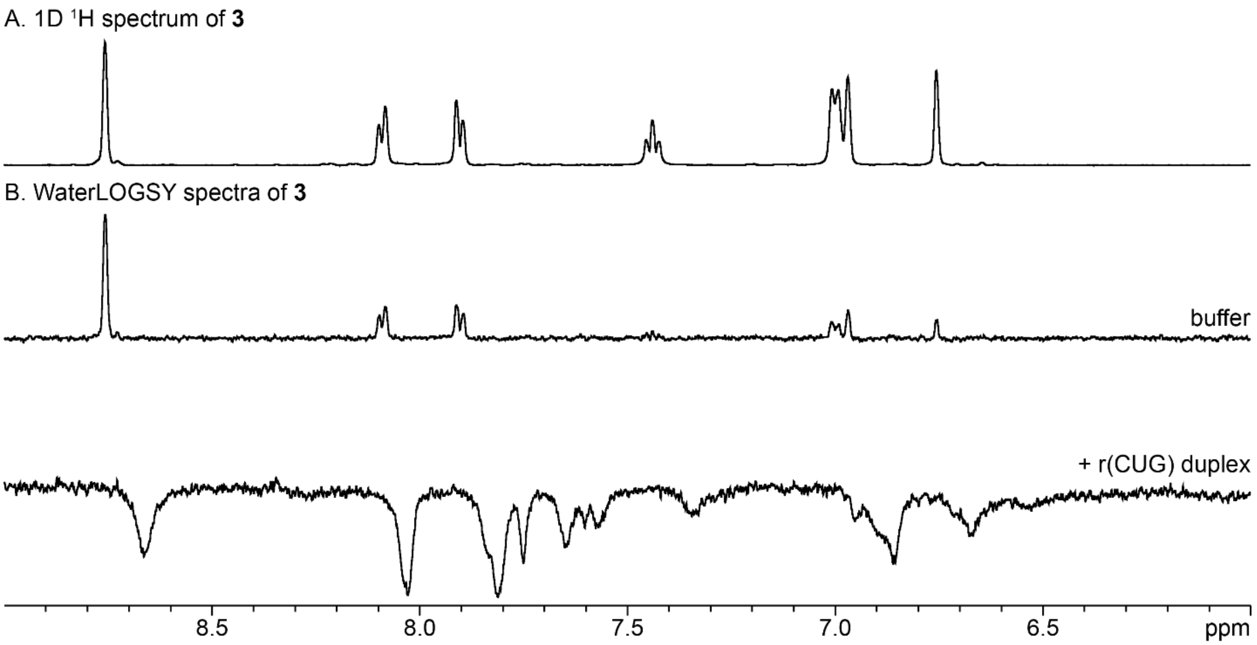


**Figure S7: 1D $^{1}$H and WaterLOGSY NMR spectra of 3 alone and in complex with the model r(CUG) duplex.** (A) 1D 1H NMR spectrum of 300 µM of **3**. (B) WaterLOGSY spectra of 300 µM of **3** in buffer alone or with 10 µM of the model r(CUG) duplex. Spectra were acquired at 25 °C.

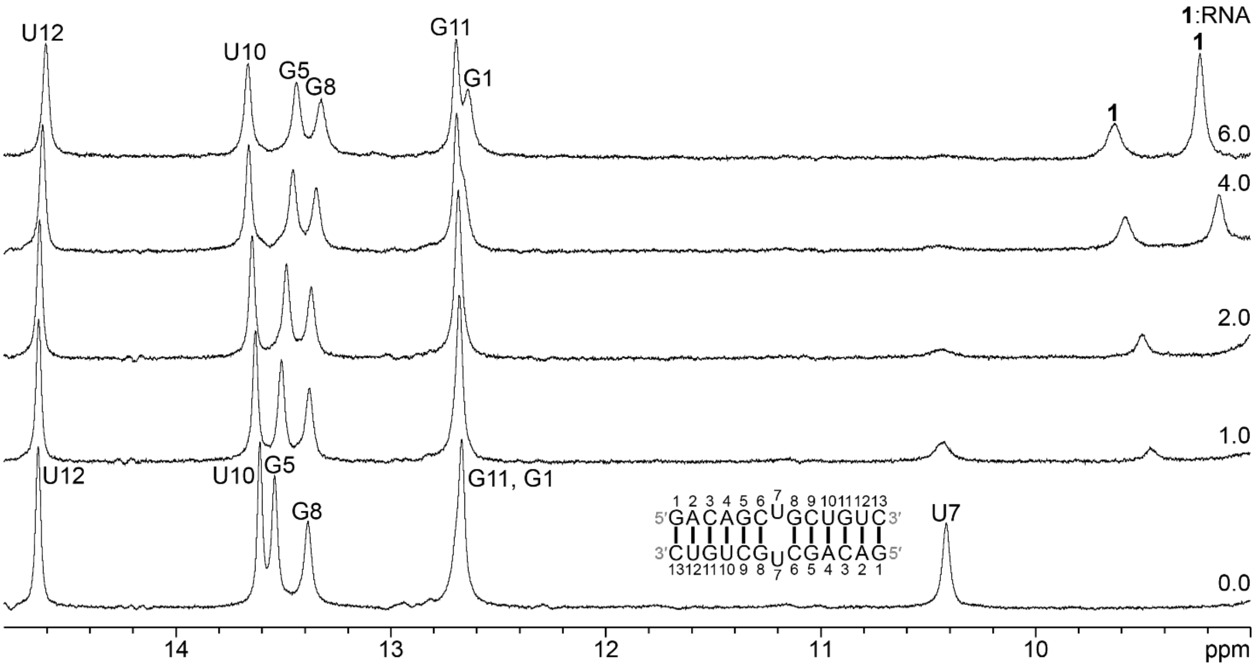


**Figure S8: 1D $^{1}$H imino region of the model r(CUG) duplex upon titration of 1.** The molar ratio of **1**:r(CUG) RNA is on the right. Spectra were acquired at 5 °C with 50 μM of RNA and 0 to 300 μM of **1**.

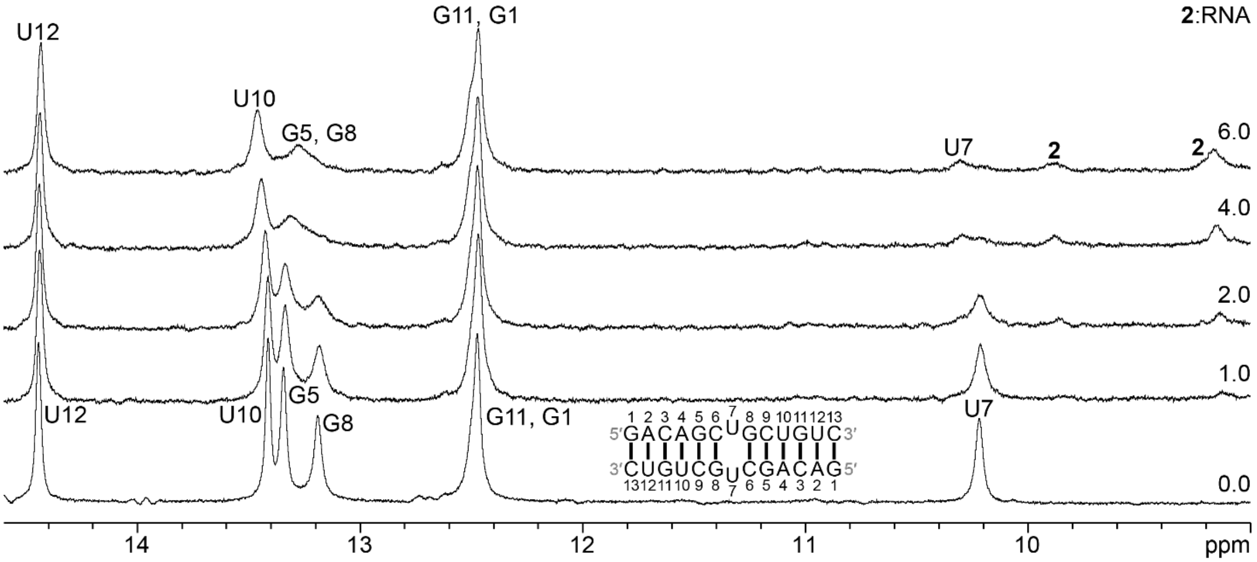


**Figure S9: 1D 1H imino region of the model r(CUG) duplex upon titration of 2.** The molar ratio of **2**:r(CUG) RNA is indicated on the right. Spectra were acquired at 5 °C with 50 μM of RNA and 0 to 300 μM of **2**.

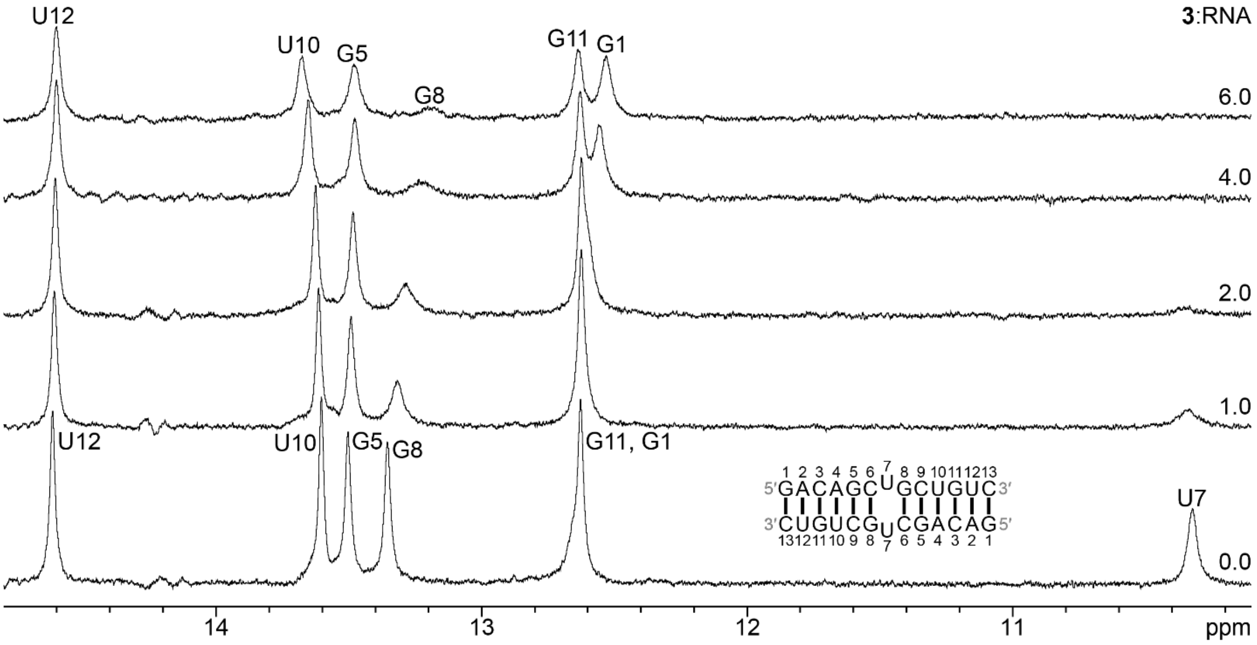


**Figure S10: 1D 1H imino region of the model r(CUG) duplex upon titration of 3.** The molar ratio of **3**:r(CUG) RNA is indicated on the right. Spectra were acquired at 5 °C with 50 μM of RNA and 0 to 300 μM of **3**.

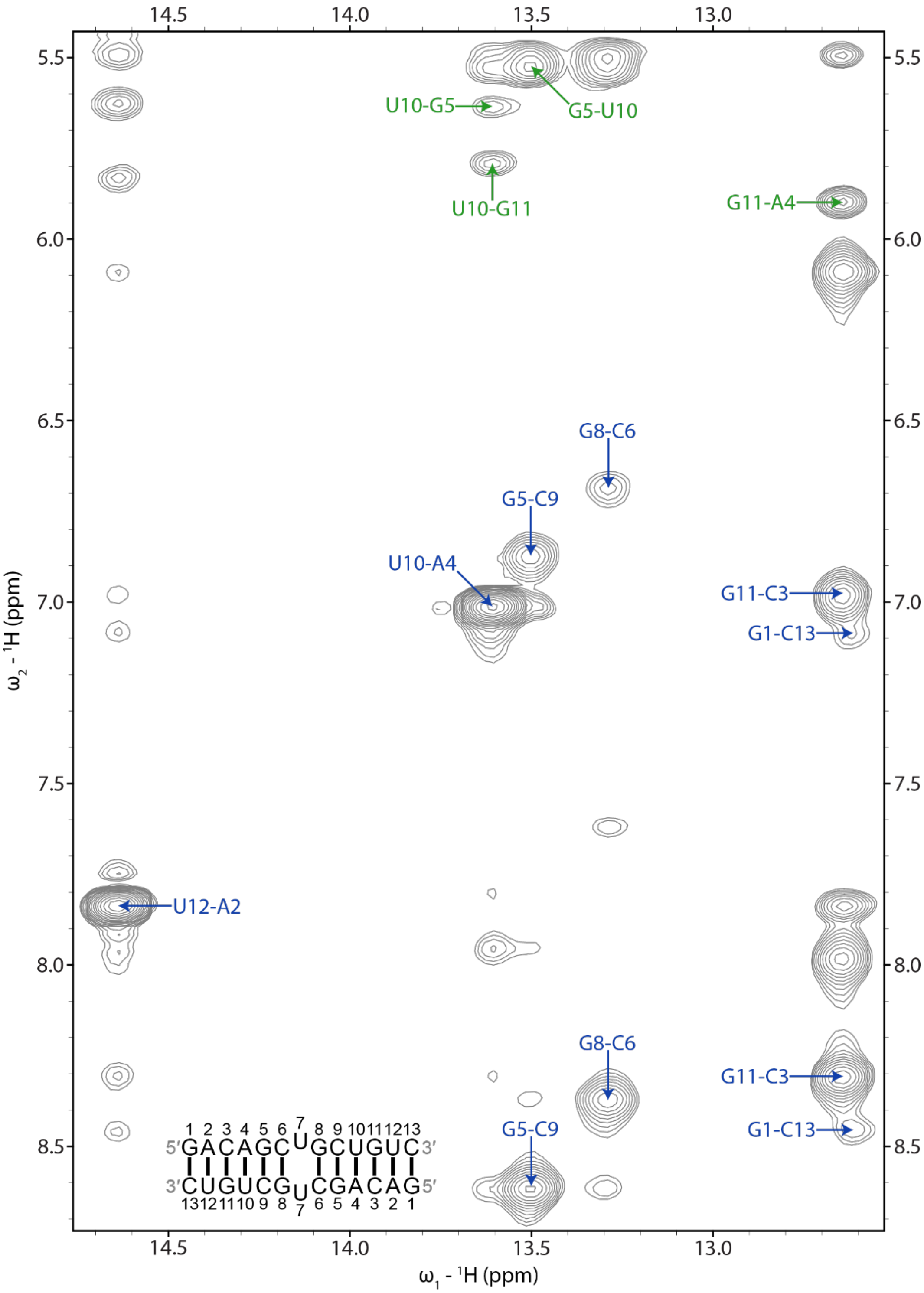


**Figure S11: Imino proton region of a 2D $^{1}$H NOESY spectrum of the r(CUG)-1 complex.** Blue labels represent UH3-AH2 and GH1 to C amino NOEs within base pairs. Green labels represent NOEs between UH3 or GH1 and the H1′ of a 3′ adjacent or cross-strand residue. In each label, the first residue is UH3 or GH1 and the second label is adenine H2, a cytosine amino proton, or H1′ of any residue. The spectrum was acquired at 6 °C with 125 ms mixing time and 0.3 mM of RNA and 0.6 mM of **1**.

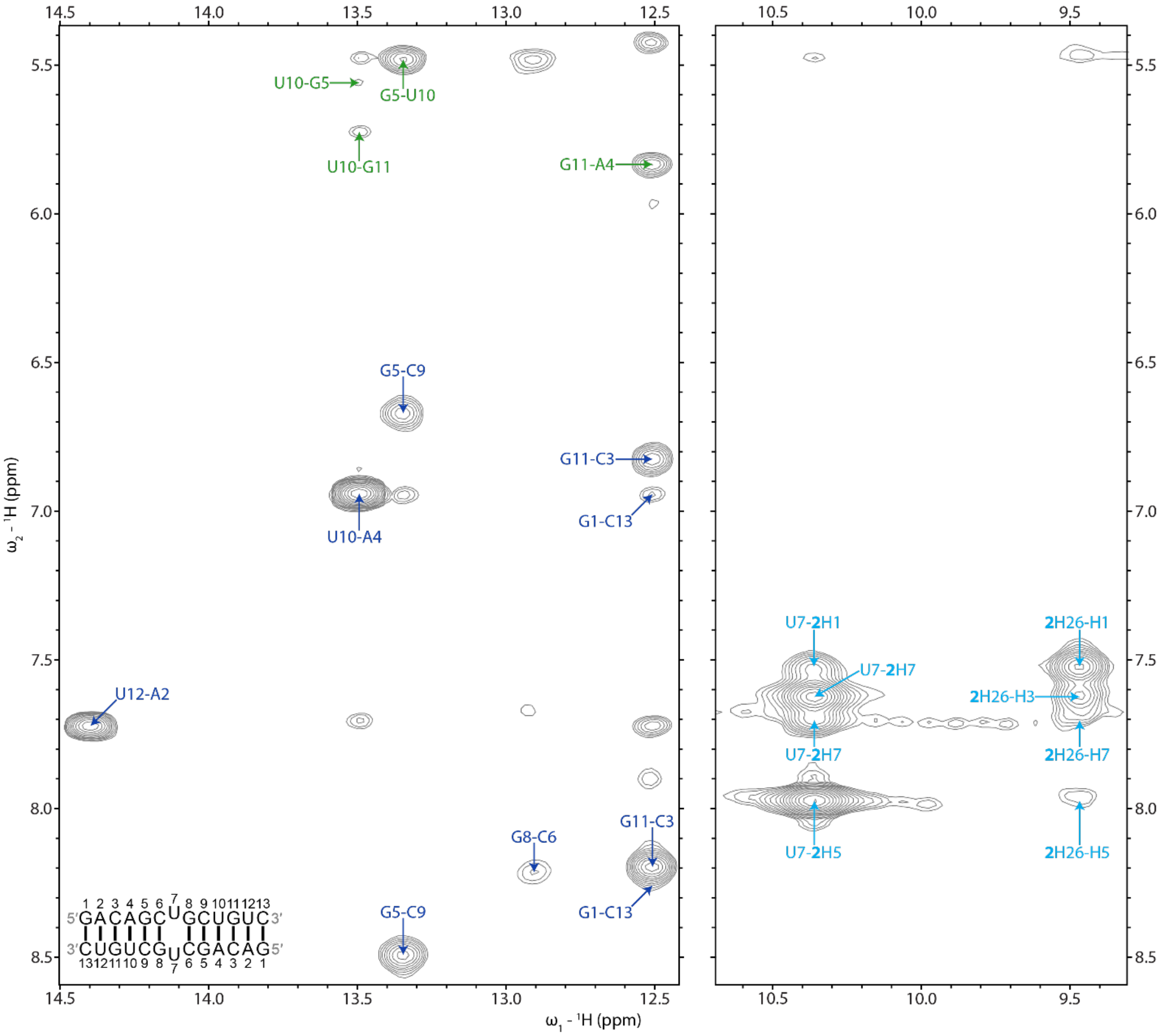


**Figure S12: Imino proton region of a 2D $^1$H NOESY spectrum of the r(CUG)-2 complex.** Blue labels represent UH3-AH2 and GH1 to C amino NOEs within base pairs. Green labels represent NOEs between UH3 or GH1 and the H1′ of a 3′ adjacent or cross-strand residue. In each label, the first residue is UH3 or GH1 and the second label is adenine H2, a cytosine amino proton, or H1′ of any residue. Light blue labels represent NOEs between exchangeable U7H3 or **2**-H26 protons and nonexchangeable protons of **2**. The spectrum was acquired at 15 °C with 125 ms mixing time and 0.3 mM of RNA and 0.6 mM of **2**.

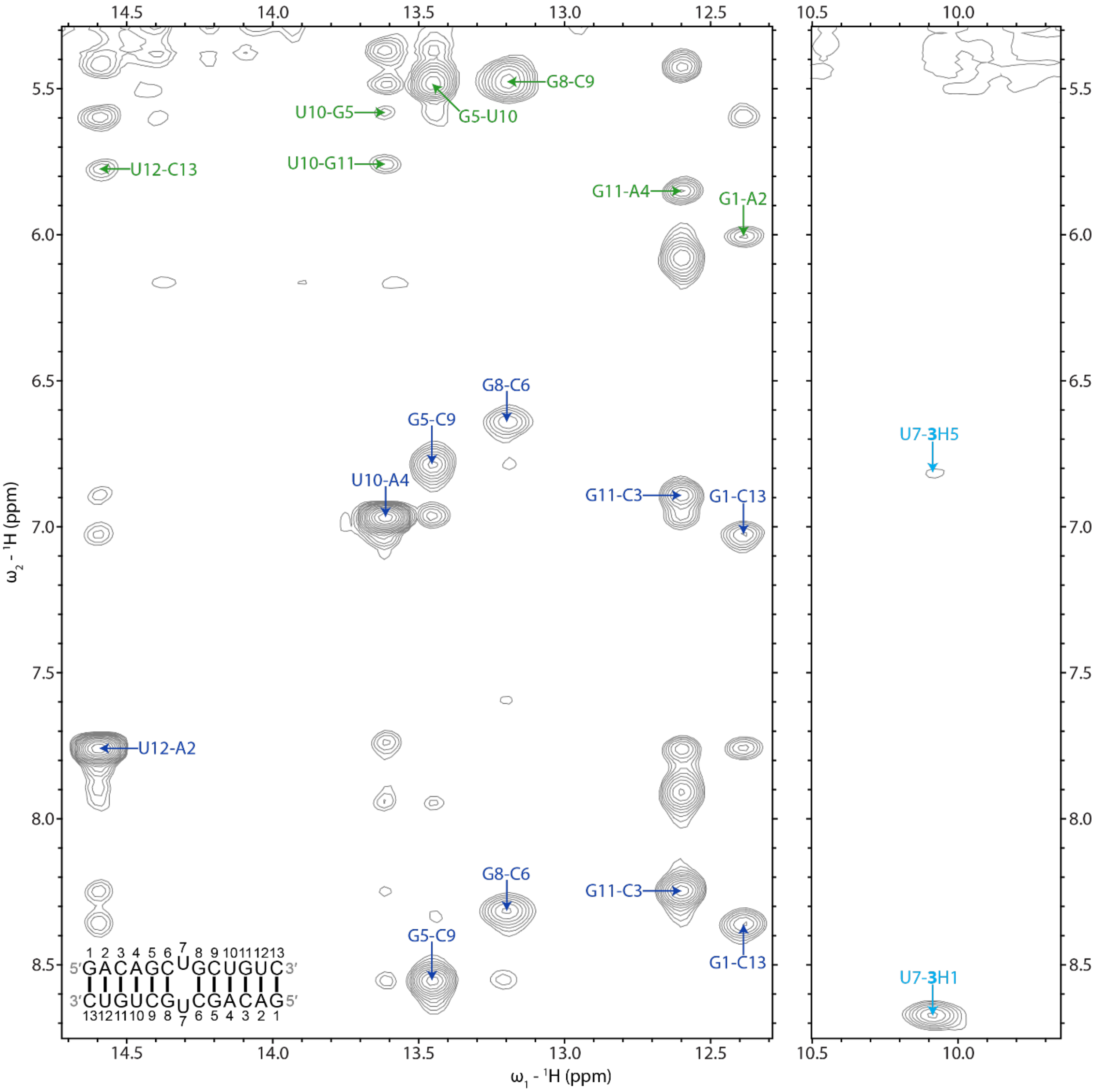


**Figure S13: Imino proton region of a 2D $^{1}$H NOESY spectrum of the r(CUG)-3 complex.** Blue labels represent UH3-AH2 and GH1 to C amino NOEs within base pairs. Green labels represent NOEs between UH3 or GH1 and the H1′ of a 3′ adjacent or cross-strand residue. In each label, the first residue is UH3 or GH1 and the second label is adenine H2, a cytosine amino proton, or H1′ of any residue. Light blue labels represent NOEs between U7H3 and **3**-H5 or -H1. The spectrum was acquired at 5 °C with 125 ms mixing time and 0.4 mM of RNA and 0.6 mM of **3**.

**Table S1:** Physicochemical properties of **1**, **2**, and **3**. Abbreviations are defined as follows: MW, molecular weight; NHAT, number of heavy atoms; NAHA, number of aromatic heavy atoms; Nrot, number of rotatable bonds; NHA, number of hydrogen bond acceptors; NHB, number of hydrogen bond donors; MF, molar refractivity; TPSA, topological polar surface area; PAINS, pan-assay interference compounds; DL, druglikeness as defined by Lipinski's Rule of 5;[2] QED: quantitative estimate of druglikeness.

| Compound | MW (g/mol) | NHAT | NAHA | Fraction C $sp^3$ | Nrot | NHA | NHB | MF | TPSA ($Å^2$) | PAINS alert | DL (Lipinski) | QED |
|---|---|---|---|---|---|---|---|---|---|---|---|---|
| **1** | 314.34 | 23 | 12 | 0.0 | 7 | 2 | 6 | 90.98 | 154.44 | 0 | Yes | 0.22 |
| **2** | 381.49 | 28 | 12 | 0.38 | 7 | 5 | 5 | 125.83 | 71.67 | 0 | Yes | 0.52 |
| **3** | 309.39 | 23 | 12 | 0.17 | 4 | 1 | 4 | 95.41 | 89.09 | 0 | Yes | 0.50 |

**Table S2:** NOE restraints used for modeling of the unbound r(CUG) duplex.

| | | | | | | | |
|---|---|---|---|---|---|---|---|
| 1 | G | H1 | 26 | C | N3 | 1.8 | 2.4 |
| 1 | G | H1' | 1 | G | H2' | 2.17 | 3.45 |
| 1 | G | H1' | 1 | G | H3' | 3.09 | 4.93 |
| 1 | G | H22 | 26 | C | O2 | 1.8 | 2.4 |
| 1 | G | H8 | 1 | G | H1' | 2.61 | 4.16 |
| 1 | G | H8 | 1 | G | H2' | 2.48 | 3.96 |
| 1 | G | H8 | 1 | G | H3' | 2.32 | 3.7 |
| 1 | G | O6 | 26 | C | H41 | 1.8 | 2.4 |
| 2 | A | H1' | 1 | G | H2' | 3.55 | 5.66 |
| 2 | A | H1' | 2 | A | H2' | 2.15 | 3.43 |
| 2 | A | H2 | 2 | A | H1' | 3 | 6 |
| 2 | A | H2 | 3 | C | H1' | 2.71 | 4.32 |
| 2 | A | H2 | 26 | C | H1' | 2.88 | 4.59 |
| 2 | A | H61 | 25 | U | O4 | 1.8 | 2.4 |
| 2 | A | H8 | 1 | G | H2' | 2.09 | 3.33 |
| 2 | A | H8 | 1 | G | H3' | 2.28 | 3.64 |
| 2 | A | H8 | 2 | A | H1' | 3 | 4.79 |
| 2 | A | H8 | 2 | A | H2' | 3.25 | 5.18 |
| 2 | A | H8 | 3 | C | H5 | 3 | 6 |
| 2 | A | N1 | 25 | U | H3 | 1.8 | 2.4 |
| 3 | C | H1' | 3 | C | H2' | 2.2 | 3.5 |
| 3 | C | H2' | 3 | C | H5 | 3 | 6 |
| 3 | C | H41 | 24 | G | O6 | 1.8 | 2.4 |
| 3 | C | H6 | 3 | C | H1' | 2.68 | 4.27 |
| 3 | C | H6 | 3 | C | H2' | 1.8 | 4.5 |
| 3 | C | N3 | 24 | G | H1 | 1.8 | 2.4 |
| 3 | C | O2 | 24 | G | H22 | 1.8 | 2.4 |
| 4 | A | H1' | 3 | C | H2' | 3.7 | 5.91 |
| 4 | A | H1' | 4 | A | H2' | 2.17 | 3.45 |
| 4 | A | H2 | 4 | A | H1' | 3 | 6 |
| 4 | A | H2 | 5 | G | H1' | 2.84 | 4.52 |
| 4 | A | H2 | 23 | U | H1' | 3 | 6 |
| 4 | A | H2 | 24 | G | H1' | 2.65 | 4.22 |
| 4 | A | H2' | 5 | G | H1' | 3.47 | 5.53 |
| 4 | A | H61 | 23 | U | O4 | 1.8 | 2.4 |
| 4 | A | H8 | 3 | C | H2' | 1.86 | 2.97 |
| 4 | A | H8 | 3 | C | H3' | 2.28 | 3.64 |
| 4 | A | H8 | 3 | C | H5 | 3 | 6 |
| 4 | A | H8 | 4 | A | H1' | 3.44 | 5.48 |
| 4 | A | H8 | 4 | A | H2' | 2.51 | 4.01 |
| 4 | A | N1 | 23 | U | H3 | 1.8 | 2.4 |
| 5 | G | H1 | 22 | C | N3 | 1.8 | 2.4 |
| 5 | G | H1' | 5 | G | H2' | 2.1 | 3.35 |
| 5 | G | H1' | 5 | G | H3' | 2.95 | 4.7 |
| 5 | G | H2' | 6 | C | H5 | 2.65 | 4.23 |
| 5 | G | H22 | 22 | C | O2 | 1.8 | 2.4 |
| 5 | G | H3' | 6 | C | H5 | 2.72 | 4.35 |
| 5 | G | H8 | 4 | A | H2' | 1.97 | 3.14 |

| | | | | | | | |
|---|---|---|---|---|---|---|---|
| 5 | G | H8 | 5 | G | H1' | 3.11 | 4.95 |
| 5 | G | H8 | 5 | G | H2' | 3.26 | 5.2 |
| 5 | G | H8 | 5 | G | H3' | 2.2 | 3.5 |
| 5 | G | H8 | 6 | C | H5 | 3 | 6 |
| 5 | G | O6 | 22 | C | H41 | 1.8 | 2.4 |
| 6 | C | H2' | 6 | C | H5 | 3.27 | 5.22 |
| 6 | C | H41 | 21 | G | O6 | 1.8 | 2.4 |
| 6 | C | H6 | 5 | G | H2' | 1.79 | 2.85 |
| 6 | C | H6 | 5 | G | H3' | 1.8 | 4.5 |
| 6 | C | H6 | 6 | C | H1' | 2.63 | 4.19 |
| 6 | C | H6 | 6 | C | H2' | 1.8 | 4.5 |
| 6 | C | N3 | 21 | G | H1 | 1.8 | 2.4 |
| 6 | C | O2 | 21 | G | H22 | 1.8 | 2.4 |
| 7 | U | H3 | 8 | G | H1' | 3 | 7 |
| 7 | U | H3 | 21 | G | H1' | 3 | 7 |
| 7 | U | H6 | 6 | C | H2' | 1.8 | 3 |
| 7 | U | H6 | 6 | C | H5 | 3 | 6 |
| 7 | U | H6 | 6 | C | H6 | 3 | 6 |
| 8 | G | H1 | 19 | C | N3 | 1.8 | 2.4 |
| 8 | G | H1' | 7 | U | H2' | 3.21 | 5.12 |
| 8 | G | H1' | 8 | G | H2' | 2.13 | 3.4 |
| 8 | G | H2' | 9 | C | H5 | 2.62 | 4.17 |
| 8 | G | H22 | 19 | C | O2 | 1.8 | 2.4 |
| 8 | G | H3' | 9 | C | H5 | 3.22 | 5.14 |
| 8 | G | H8 | 7 | U | H1' | 3 | 6 |
| 8 | G | H8 | 7 | U | H2' | 1.89 | 3.01 |
| 8 | G | H8 | 7 | U | H6 | 3 | 6 |
| 8 | G | H8 | 8 | G | H1' | 3.09 | 4.93 |
| 8 | G | H8 | 8 | G | H2' | 2.79 | 4.45 |
| 8 | G | H8 | 8 | G | H3' | 2.08 | 3.31 |
| 8 | G | H8 | 9 | C | H5 | 3.55 | 5.66 |
| 8 | G | O6 | 19 | C | H41 | 1.8 | 2.4 |
| 9 | C | H1' | 9 | C | H3' | 2.4 | 3.83 |
| 9 | C | H2' | 9 | C | H5 | 3 | 6 |
| 9 | C | H3' | 9 | C | H5 | 2.98 | 4.75 |
| 9 | C | H41 | 18 | G | O6 | 1.8 | 2.4 |
| 9 | C | H6 | 8 | G | H2' | 1.88 | 3 |
| 9 | C | H6 | 9 | C | H1' | 2.75 | 4.38 |
| 9 | C | H6 | 9 | C | H2' | 2.66 | 4.25 |
| 9 | C | H6 | 9 | C | H3' | 1.89 | 3.01 |
| 9 | C | H6 | 10 | U | H5 | 3.32 | 5.29 |
| 9 | C | N3 | 18 | G | H1 | 1.8 | 2.4 |
| 9 | C | O2 | 18 | G | H22 | 1.8 | 2.4 |
| 10 | U | H1' | 10 | U | H2' | 2.31 | 3.68 |
| 10 | U | H1' | 10 | U | H3' | 2.69 | 4.28 |
| 10 | U | H2' | 10 | U | H5 | 3 | 6 |
| 10 | U | H3 | 17 | A | N1 | 1.8 | 2.4 |
| 10 | U | H3' | 10 | U | H5 | 3 | 6 |
| 10 | U | H6 | 9 | C | H2' | 1.76 | 2.81 |
| 10 | U | H6 | 9 | C | H3' | 2.17 | 3.46 |
| 10 | U | H6 | 9 | C | H5 | 3 | 6 |

| | | | | | | | |
|---|---|---|---|---|---|---|---|
| 10 | U | H6 | 10 | U | H2' | 2.79 | 4.45 |
| 10 | U | H6 | 10 | U | H3' | 1.93 | 3.09 |
| 10 | U | O4 | 17 | A | H61 | 1.8 | 2.4 |
| 11 | G | H1 | 16 | C | N3 | 1.8 | 2.4 |
| 11 | G | H1' | 11 | G | H2' | 2.04 | 3.25 |
| 11 | G | H1' | 11 | G | H3' | 2.66 | 4.25 |
| 11 | G | H2' | 12 | U | H5 | 2.5 | 3.99 |
| 11 | G | H22 | 16 | C | O2 | 1.8 | 2.4 |
| 11 | G | H3' | 12 | U | H5 | 2.65 | 4.22 |
| 11 | G | H8 | 10 | U | H2' | 1.88 | 3 |
| 11 | G | H8 | 11 | G | H1' | 2.86 | 4.56 |
| 11 | G | H8 | 11 | G | H3' | 2.11 | 3.37 |
| 11 | G | H8 | 12 | U | H5 | 3.56 | 5.67 |
| 11 | G | O6 | 16 | C | H41 | 1.8 | 2.4 |
| 12 | U | H1' | 12 | U | H2' | 1.98 | 3.15 |
| 12 | U | H1' | 12 | U | H3' | 2.66 | 4.25 |
| 12 | U | H2' | 12 | U | H5 | 3 | 6 |
| 12 | U | H3 | 15 | A | N1 | 1.8 | 2.4 |
| 12 | U | H3' | 12 | U | H5 | 3.05 | 4.86 |
| 12 | U | H6 | 11 | G | H2' | 1.67 | 2.66 |
| 12 | U | H6 | 11 | G | H3' | 2.12 | 3.39 |
| 12 | U | H6 | 12 | U | H1' | 2.53 | 4.03 |
| 12 | U | H6 | 12 | U | H2' | 2.55 | 4.07 |
| 12 | U | H6 | 12 | U | H3' | 1.91 | 3.05 |
| 12 | U | H6 | 13 | C | H5 | 3.19 | 5.09 |
| 12 | U | O4 | 15 | A | H61 | 1.8 | 2.4 |
| 13 | C | H1' | 13 | C | H2' | 2.01 | 3.2 |
| 13 | C | H1' | 13 | C | H3' | 2.65 | 4.23 |
| 13 | C | H2' | 13 | C | H5 | 3 | 6 |
| 13 | C | H3' | 13 | C | H5 | 3.03 | 4.83 |
| 13 | C | H41 | 14 | G | O6 | 1.8 | 2.4 |
| 13 | C | H5 | 12 | U | H2' | 2.8 | 4.46 |
| 13 | C | H5 | 12 | U | H3' | 3.03 | 4.84 |
| 13 | C | H6 | 12 | U | H2' | 1.84 | 2.94 |
| 13 | C | H6 | 12 | U | H3' | 2.2 | 3.52 |
| 13 | C | H6 | 13 | C | H1' | 2.56 | 4.08 |
| 13 | C | H6 | 13 | C | H2' | 2.28 | 3.63 |
| 13 | C | H6 | 13 | C | H3' | 1.85 | 2.95 |
| 13 | C | N3 | 14 | G | H1 | 1.8 | 2.4 |
| 13 | C | O2 | 14 | G | H22 | 1.8 | 2.4 |
| 14 | G | H1' | 14 | G | H2' | 2.17 | 3.45 |
| 14 | G | H1' | 14 | G | H3' | 3.09 | 4.93 |
| 14 | G | H8 | 14 | G | H1' | 2.61 | 4.16 |
| 14 | G | H8 | 14 | G | H2' | 2.48 | 3.96 |
| 14 | G | H8 | 14 | G | H3' | 2.32 | 3.7 |
| 15 | A | H1' | 14 | G | H2' | 3.55 | 5.66 |
| 15 | A | H1' | 15 | A | H2' | 2.15 | 3.43 |
| 15 | A | H2 | 13 | C | H1' | 2.88 | 4.59 |
| 15 | A | H2 | 15 | A | H1' | 3 | 6 |
| 15 | A | H2 | 16 | C | H1' | 2.71 | 4.32 |
| 15 | A | H8 | 14 | G | H2' | 2.09 | 3.33 |

| | | | | | | | |
|---|---|---|---|---|---|---|---|
| 15 | A | H8 | 14 | G | H3' | 2.28 | 3.64 |
| 15 | A | H8 | 15 | A | H1' | 3 | 4.79 |
| 15 | A | H8 | 15 | A | H2' | 3.25 | 5.18 |
| 15 | A | H8 | 16 | C | H5 | 3 | 6 |
| 16 | C | H1' | 16 | C | H2' | 2.2 | 3.5 |
| 16 | C | H2' | 16 | C | H5 | 3 | 6 |
| 16 | C | H6 | 16 | C | H1' | 2.68 | 4.27 |
| 16 | C | H6 | 16 | C | H2' | 1.8 | 4.5 |
| 17 | A | H1' | 16 | C | H2' | 3.7 | 5.91 |
| 17 | A | H1' | 17 | A | H2' | 2.17 | 3.45 |
| 17 | A | H2 | 10 | U | H1' | 3 | 6 |
| 17 | A | H2 | 11 | G | H1' | 2.65 | 4.22 |
| 17 | A | H2 | 17 | A | H1' | 3 | 6 |
| 17 | A | H2 | 18 | G | H1' | 2.84 | 4.52 |
| 17 | A | H2' | 18 | G | H1' | 3.47 | 5.53 |
| 17 | A | H8 | 16 | C | H2' | 1.86 | 2.97 |
| 17 | A | H8 | 16 | C | H3' | 2.28 | 3.64 |
| 17 | A | H8 | 16 | C | H5 | 3 | 6 |
| 17 | A | H8 | 17 | A | H1' | 3.44 | 5.48 |
| 17 | A | H8 | 17 | A | H2' | 2.51 | 4.01 |
| 18 | G | H1' | 18 | G | H2' | 2.1 | 3.35 |
| 18 | G | H1' | 18 | G | H3' | 2.95 | 4.7 |
| 18 | G | H2' | 19 | C | H5 | 2.65 | 4.23 |
| 18 | G | H3' | 19 | C | H5 | 2.72 | 4.35 |
| 18 | G | H8 | 17 | A | H2' | 1.97 | 3.14 |
| 18 | G | H8 | 18 | G | H1' | 3.11 | 4.95 |
| 18 | G | H8 | 18 | G | H2' | 3.26 | 5.2 |
| 18 | G | H8 | 18 | G | H3' | 2.2 | 3.5 |
| 18 | G | H8 | 19 | C | H5 | 3 | 6 |
| 19 | C | H2' | 19 | C | H5 | 3.27 | 5.22 |
| 19 | C | H6 | 18 | G | H2' | 1.79 | 2.85 |
| 19 | C | H6 | 18 | G | H3' | 1.8 | 4.5 |
| 19 | C | H6 | 19 | C | H1' | 2.63 | 4.19 |
| 19 | C | H6 | 19 | C | H2' | 1.8 | 4.5 |
| 20 | U | H3 | 8 | G | H1' | 3 | 7 |
| 20 | U | H3 | 21 | G | H1' | 3 | 7 |
| 20 | U | H6 | 19 | C | H2' | 1.8 | 3 |
| 20 | U | H6 | 19 | C | H5 | 3 | 6 |
| 20 | U | H6 | 19 | C | H6 | 3 | 6 |
| 21 | G | H1' | 20 | U | H2' | 3.21 | 5.12 |
| 21 | G | H1' | 21 | G | H2' | 2.13 | 3.4 |
| 21 | G | H2' | 22 | C | H5 | 2.62 | 4.17 |
| 21 | G | H3' | 22 | C | H5 | 3.22 | 5.14 |
| 21 | G | H8 | 20 | U | H1' | 3 | 6 |
| 21 | G | H8 | 20 | U | H2' | 1.89 | 3.01 |
| 21 | G | H8 | 20 | U | H6 | 3 | 6 |
| 21 | G | H8 | 21 | G | H1' | 3.09 | 4.93 |
| 21 | G | H8 | 21 | G | H2' | 2.79 | 4.45 |
| 21 | G | H8 | 21 | G | H3' | 2.08 | 3.31 |
| 21 | G | H8 | 22 | C | H5 | 3.55 | 5.66 |
| 22 | C | H1' | 22 | C | H3' | 2.4 | 3.83 |

| | | | | | | | |
|---|---|---|---|---|---|---|---|
| 22 | C | H2' | 22 | C | H5 | 3 | 6 |
| 22 | C | H3' | 22 | C | H5 | 2.98 | 4.75 |
| 22 | C | H6 | 21 | G | H2' | 1.88 | 3 |
| 22 | C | H6 | 22 | C | H1' | 2.75 | 4.38 |
| 22 | C | H6 | 22 | C | H2' | 2.66 | 4.25 |
| 22 | C | H6 | 22 | C | H3' | 1.89 | 3.01 |
| 22 | C | H6 | 23 | U | H5 | 3.32 | 5.29 |
| 23 | U | H1' | 23 | U | H2' | 2.31 | 3.68 |
| 23 | U | H1' | 23 | U | H3' | 2.69 | 4.28 |
| 23 | U | H2' | 23 | U | H5 | 3 | 6 |
| 23 | U | H3' | 23 | U | H5 | 3 | 6 |
| 23 | U | H6 | 22 | C | H2' | 1.76 | 2.81 |
| 23 | U | H6 | 22 | C | H3' | 2.17 | 3.46 |
| 23 | U | H6 | 22 | C | H5 | 3 | 6 |
| 23 | U | H6 | 23 | U | H2' | 2.79 | 4.45 |
| 23 | U | H6 | 23 | U | H3' | 1.93 | 3.09 |
| 24 | G | H1' | 24 | G | H2' | 2.04 | 3.25 |
| 24 | G | H1' | 24 | G | H3' | 2.66 | 4.25 |
| 24 | G | H2' | 25 | U | H5 | 2.5 | 3.99 |
| 24 | G | H3' | 25 | U | H5 | 2.65 | 4.22 |
| 24 | G | H8 | 23 | U | H2' | 1.88 | 3 |
| 24 | G | H8 | 24 | G | H1' | 2.86 | 4.56 |
| 24 | G | H8 | 24 | G | H3' | 2.11 | 3.37 |
| 24 | G | H8 | 25 | U | H5 | 3.56 | 5.67 |
| 25 | U | H1' | 25 | U | H2' | 1.98 | 3.15 |
| 25 | U | H1' | 25 | U | H3' | 2.66 | 4.25 |
| 25 | U | H2' | 25 | U | H5 | 3 | 6 |
| 25 | U | H3' | 25 | U | H5 | 3.05 | 4.86 |
| 25 | U | H6 | 24 | G | H2' | 1.67 | 2.66 |
| 25 | U | H6 | 24 | G | H3' | 2.12 | 3.39 |
| 25 | U | H6 | 25 | U | H1' | 2.53 | 4.03 |
| 25 | U | H6 | 25 | U | H2' | 2.55 | 4.07 |
| 25 | U | H6 | 25 | U | H3' | 1.91 | 3.05 |
| 25 | U | H6 | 26 | C | H5 | 3.19 | 5.09 |
| 26 | C | H1' | 26 | C | H2' | 2.01 | 3.2 |
| 26 | C | H1' | 26 | C | H3' | 2.65 | 4.23 |
| 26 | C | H2' | 26 | C | H5 | 3 | 6 |
| 26 | C | H3' | 26 | C | H5 | 3.03 | 4.83 |
| 26 | C | H5 | 25 | U | H2' | 2.8 | 4.46 |
| 26 | C | H5 | 25 | U | H3' | 3.03 | 4.84 |
| 26 | C | H6 | 25 | U | H2' | 1.84 | 2.94 |
| 26 | C | H6 | 25 | U | H3' | 2.2 | 3.52 |
| 26 | C | H6 | 26 | C | H1' | 2.56 | 4.08 |
| 26 | C | H6 | 26 | C | H2' | 2.28 | 3.63 |
| 26 | C | H6 | 26 | C | H3' | 1.85 | 2.95 |

Table S3: Dihedral restraints used for modeling of the unbound r(CUG) duplex.

| | | | |
|---|---|---|---|
| ALPHA | (1 RG5 O3')-(2 RA P)-(2 RA O5')-(2 RA C5') | -155.0 | 25.0 |
| ALPHA | (2 RA O3')-(3 RC P)-(3 RC O5')-(3 RC C5') | -155.0 | 25.0 |
| ALPHA | (3 RC O3')-(4 RA P)-(4 RA O5')-(4 RA C5') | -155.0 | 25.0 |
| ALPHA | (4 RA O3')-(5 RG P)-(5 RG O5')-(5 RG C5') | -155.0 | 25.0 |
| ALPHA | (5 RG O3')-(6 RC P)-(6 RC O5')-(6 RC C5') | -155.0 | 25.0 |
| ALPHA | (7 RU O3')-(8 RG P)-(8 RG O5')-(8 RG C5') | -155.0 | 25.0 |
| ALPHA | (8 RG O3')-(9 RC P)-(9 RC O5')-(9 RC C5') | -155.0 | 25.0 |
| ALPHA | (9 RC O3')-(10 RU P)-(10 RU O5')-(10 RU C5') | -155.0 | 25.0 |
| ALPHA | (10 RU O3')-(11 RG P)-(11 RG O5')-(11 RG C5') | -155.0 | 25.0 |
| ALPHA | (11 RG O3')-(12 RU P)-(12 RU O5')-(12 RU C5') | -155.0 | 25.0 |
| ALPHA | (14 RG5 O3')-(15 RA P)-(15 RA O5')-(15 RA C5') | -155.0 | 25.0 |
| ALPHA | (15 RA O3')-(16 RC P)-(16 RC O5')-(16 RC C5') | -155.0 | 25.0 |
| ALPHA | (16 RC O3')-(17 RA P)-(17 RA O5')-(17 RA C5') | -155.0 | 25.0 |
| ALPHA | (17 RA O3')-(18 RG P)-(18 RG O5')-(18 RG C5') | -155.0 | 25.0 |
| ALPHA | (18 RG O3')-(19 RC P)-(19 RC O5')-(19 RC C5') | -155.0 | 25.0 |
| ALPHA | (20 RU O3')-(21 RG P)-(21 RG O5')-(21 RG C5') | -155.0 | 25.0 |
| ALPHA | (21 RG O3')-(22 RC P)-(22 RC O5')-(22 RC C5') | -155.0 | 25.0 |
| ALPHA | (22 RC O3')-(23 RU P)-(23 RU O5')-(23 RU C5') | -155.0 | 25.0 |
| ALPHA | (23 RU O3')-(24 RG P)-(24 RG O5')-(24 RG C5') | -155.0 | 25.0 |
| ALPHA | (24 RG O3')-(25 RU P)-(25 RU O5')-(25 RU C5') | -155.0 | 25.0 |
| BETA | (2 RA P)-(2 RA O5')-(2 RA C5')-(2 RA C4') | 90.0 | 240.0 |
| BETA | (3 RC P)-(3 RC O5')-(3 RC C5')-(3 RC C4') | 90.0 | 240.0 |
| BETA | (4 RA P)-(4 RA O5')-(4 RA C5')-(4 RA C4') | 90.0 | 240.0 |
| BETA | (5 RG P)-(5 RG O5')-(5 RG C5')-(5 RG C4') | 90.0 | 240.0 |
| BETA | (6 RC P)-(6 RC O5')-(6 RC C5')-(6 RC C4') | 90.0 | 240.0 |
| BETA | (8 RG P)-(8 RG O5')-(8 RG C5')-(8 RG C4') | 90.0 | 240.0 |
| BETA | (9 RC P)-(9 RC O5')-(9 RC C5')-(9 RC C4') | 90.0 | 240.0 |
| BETA | (10 RU P)-(10 RU O5')-(10 RU C5')-(10 RU C4') | 90.0 | 240.0 |
| BETA | (11 RG P)-(11 RG O5')-(11 RG C5')-(11 RG C4') | 90.0 | 240.0 |
| BETA | (12 RU P)-(12 RU O5')-(12 RU C5')-(12 RU C4') | 90.0 | 240.0 |
| BETA | (15 RA P)-(15 RA O5')-(15 RA C5')-(15 RA C4') | 90.0 | 240.0 |
| BETA | (16 RC P)-(16 RC O5')-(16 RC C5')-(16 RC C4') | 90.0 | 240.0 |
| BETA | (17 RA P)-(17 RA O5')-(17 RA C5')-(17 RA C4') | 90.0 | 240.0 |
| BETA | (18 RG P)-(18 RG O5')-(18 RG C5')-(18 RG C4') | 90.0 | 240.0 |
| BETA | (19 RC P)-(19 RC O5')-(19 RC C5')-(19 RC C4') | 90.0 | 240.0 |
| BETA | (21 RG P)-(21 RG O5')-(21 RG C5')-(21 RG C4') | 90.0 | 240.0 |
| BETA | (22 RC P)-(22 RC O5')-(22 RC C5')-(22 RC C4') | 90.0 | 240.0 |
| BETA | (23 RU P)-(23 RU O5')-(23 RU C5')-(23 RU C4') | 90.0 | 240.0 |
| BETA | (24 RG P)-(24 RG O5')-(24 RG C5')-(24 RG C4') | 90.0 | 240.0 |
| BETA | (25 RU P)-(25 RU O5')-(25 RU C5')-(25 RU C4') | 90.0 | 240.0 |
| GAMMA | (2 RA O5')-(2 RA C5')-(2 RA C4')-(2 RA C3') | 0.0 | 120.0 |
| GAMMA | (3 RC O5')-(3 RC C5')-(3 RC C4')-(3 RC C3') | 0.0 | 120.0 |
| GAMMA | (4 RA O5')-(4 RA C5')-(4 RA C4')-(4 RA C3') | 0.0 | 120.0 |
| GAMMA | (5 RG O5')-(5 RG C5')-(5 RG C4')-(5 RG C3') | 0.0 | 120.0 |
| GAMMA | (6 RC O5')-(6 RC C5')-(6 RC C4')-(6 RC C3') | 0.0 | 120.0 |
| GAMMA | (8 RG O5')-(8 RG C5')-(8 RG C4')-(8 RG C3') | 0.0 | 120.0 |
| GAMMA | (9 RC O5')-(9 RC C5')-(9 RC C4')-(9 RC C3') | 0.0 | 120.0 |
| GAMMA | (10 RU O5')-(10 RU C5')-(10 RU C4')-(10 RU C3') | 0.0 | 120.0 |
| GAMMA | (11 RG O5')-(11 RG C5')-(11 RG C4')-(11 RG C3') | 0.0 | 120.0 |

| | | | |
|---|---|---|---|
| GAMMA | (12 RU O5')-(12 RU C5')-(12 RU C4')-(12 RU C3') | 0.0 | 120.0 |
| GAMMA | (15 RA O5')-(15 RA C5')-(15 RA C4')-(15 RA C3') | 0.0 | 120.0 |
| GAMMA | (16 RC O5')-(16 RC C5')-(16 RC C4')-(16 RC C3') | 0.0 | 120.0 |
| GAMMA | (17 RA O5')-(17 RA C5')-(17 RA C4')-(17 RA C3') | 0.0 | 120.0 |
| GAMMA | (18 RG O5')-(18 RG C5')-(18 RG C4')-(18 RG C3') | 0.0 | 120.0 |
| GAMMA | (19 RC O5')-(19 RC C5')-(19 RC C4')-(19 RC C3') | 0.0 | 120.0 |
| GAMMA | (21 RG O5')-(21 RG C5')-(21 RG C4')-(21 RG C3') | 0.0 | 120.0 |
| GAMMA | (22 RC O5')-(22 RC C5')-(22 RC C4')-(22 RC C3') | 0.0 | 120.0 |
| GAMMA | (23 RU O5')-(23 RU C5')-(23 RU C4')-(23 RU C3') | 0.0 | 120.0 |
| GAMMA | (24 RG O5')-(24 RG C5')-(24 RG C4')-(24 RG C3') | 0.0 | 120.0 |
| GAMMA | (25 RU O5')-(25 RU C5')-(25 RU C4')-(25 RU C3') | 0.0 | 120.0 |
| DELTA | (2 RA C5')-(2 RA C4')-(2 RA C3')-(2 RA O3') | 45.0 | 115.0 |
| DELTA | (3 RC C5')-(3 RC C4')-(3 RC C3')-(3 RC O3') | 45.0 | 115.0 |
| DELTA | (4 RA C5')-(4 RA C4')-(4 RA C3')-(4 RA O3') | 45.0 | 115.0 |
| DELTA | (5 RG C5')-(5 RG C4')-(5 RG C3')-(5 RG O3') | 45.0 | 115.0 |
| DELTA | (6 RC C5')-(6 RC C4')-(6 RC C3')-(6 RC O3') | 45.0 | 115.0 |
| DELTA | (8 RG C5')-(8 RG C4')-(8 RG C3')-(8 RG O3') | 45.0 | 115.0 |
| DELTA | (9 RC C5')-(9 RC C4')-(9 RC C3')-(9 RC O3') | 45.0 | 115.0 |
| DELTA | (10 RU C5')-(10 RU C4')-(10 RU C3')-(10 RU O3') | 45.0 | 115.0 |
| DELTA | (11 RG C5')-(11 RG C4')-(11 RG C3')-(11 RG O3') | 45.0 | 115.0 |
| DELTA | (12 RU C5')-(12 RU C4')-(12 RU C3')-(12 RU O3') | 45.0 | 115.0 |
| DELTA | (15 RA C5')-(15 RA C4')-(15 RA C3')-(15 RA O3') | 45.0 | 115.0 |
| DELTA | (16 RC C5')-(16 RC C4')-(16 RC C3')-(16 RC O3') | 45.0 | 115.0 |
| DELTA | (17 RA C5')-(17 RA C4')-(17 RA C3')-(17 RA O3') | 45.0 | 115.0 |
| DELTA | (18 RG C5')-(18 RG C4')-(18 RG C3')-(18 RG O3') | 45.0 | 115.0 |
| DELTA | (19 RC C5')-(19 RC C4')-(19 RC C3')-(19 RC O3') | 45.0 | 115.0 |
| DELTA | (21 RG C5')-(21 RG C4')-(21 RG C3')-(21 RG O3') | 45.0 | 115.0 |
| DELTA | (22 RC C5')-(22 RC C4')-(22 RC C3')-(22 RC O3') | 45.0 | 115.0 |
| DELTA | (23 RU C5')-(23 RU C4')-(23 RU C3')-(23 RU O3') | 45.0 | 115.0 |
| DELTA | (24 RG C5')-(24 RG C4')-(24 RG C3')-(24 RG O3') | 45.0 | 115.0 |
| DELTA | (25 RU C5')-(25 RU C4')-(25 RU C3')-(25 RU O3') | 45.0 | 115.0 |
| EPSILON | (2 RA C4')-(2 RA C3')-(2 RA O3')-(3 RC P) | -240.0 | 10.0 |
| EPSILON | (3 RC C4')-(3 RC C3')-(3 RC O3')-(4 RA P) | -240.0 | 10.0 |
| EPSILON | (4 RA C4')-(4 RA C3')-(4 RA O3')-(5 RG P) | -240.0 | 10.0 |
| EPSILON | (5 RG C4')-(5 RG C3')-(5 RG O3')-(6 RC P) | -240.0 | 10.0 |
| EPSILON | (6 RC C4')-(6 RC C3')-(6 RC O3')-(7 RU P) | -240.0 | 10.0 |
| EPSILON | (8 RG C4')-(8 RG C3')-(8 RG O3')-(9 RC P) | -240.0 | 10.0 |
| EPSILON | (9 RC C4')-(9 RC C3')-(9 RC O3')-(10 RU P) | -240.0 | 10.0 |
| EPSILON | (10 RU C4')-(10 RU C3')-(10 RU O3')-(11 RG P) | -240.0 | 10.0 |
| EPSILON | (11 RG C4')-(11 RG C3')-(11 RG O3')-(12 RU P) | -240.0 | 10.0 |
| EPSILON | (12 RU C4')-(12 RU C3')-(12 RU O3')-(13 RC3 P) | -240.0 | 10.0 |
| EPSILON | (15 RA C4')-(15 RA C3')-(15 RA O3')-(16 RC P) | -240.0 | 10.0 |
| EPSILON | (16 RC C4')-(16 RC C3')-(16 RC O3')-(17 RA P) | -240.0 | 10.0 |
| EPSILON | (17 RA C4')-(17 RA C3')-(17 RA O3')-(18 RG P) | -240.0 | 10.0 |
| EPSILON | (18 RG C4')-(18 RG C3')-(18 RG O3')-(19 RC P) | -240.0 | 10.0 |
| EPSILON | (19 RC C4')-(19 RC C3')-(19 RC O3')-(20 RU P) | -240.0 | 10.0 |
| EPSILON | (21 RG C4')-(21 RG C3')-(21 RG O3')-(22 RC P) | -240.0 | 10.0 |
| EPSILON | (22 RC C4')-(22 RC C3')-(22 RC O3')-(23 RU P) | -240.0 | 10.0 |
| EPSILON | (23 RU C4')-(23 RU C3')-(23 RU O3')-(24 RG P) | -240.0 | 10.0 |
| EPSILON | (24 RG C4')-(24 RG C3')-(24 RG O3')-(25 RU P) | -240.0 | 10.0 |
| EPSILON | (25 RU C4')-(25 RU C3')-(25 RU O3')-(26 RC3 P) | -240.0 | 10.0 |

| | | | |
|---|---|---|---|
| ZETA | (2 RA C3')-(2 RA O3')-(3 RC P)-(3 RC O5') | -160.0 | 20.0 |
| ZETA | (3 RC C3')-(3 RC O3')-(4 RA P)-(4 RA O5') | -160.0 | 20.0 |
| ZETA | (4 RA C3')-(4 RA O3')-(5 RG P)-(5 RG O5') | -160.0 | 20.0 |
| ZETA | (5 RG C3')-(5 RG O3')-(6 RC P)-(6 RC O5') | -160.0 | 20.0 |
| ZETA | (6 RC C3')-(6 RC O3')-(7 RU P)-(7 RU O5') | -160.0 | 20.0 |
| ZETA | (8 RG C3')-(8 RG O3')-(9 RC P)-(9 RC O5') | -160.0 | 20.0 |
| ZETA | (9 RC C3')-(9 RC O3')-(10 RU P)-(10 RU O5') | -160.0 | 20.0 |
| ZETA | (10 RU C3')-(10 RU O3')-(11 RG P)-(11 RG O5') | -160.0 | 20.0 |
| ZETA | (11 RG C3')-(11 RG O3')-(12 RU P)-(12 RU O5') | -160.0 | 20.0 |
| ZETA | (12 RU C3')-(12 RU O3')-(13 RC3 P)-(13 RC3 O5') | -160.0 | 20.0 |
| ZETA | (15 RA C3')-(15 RA O3')-(16 RC P)-(16 RC O5') | -160.0 | 20.0 |
| ZETA | (16 RC C3')-(16 RC O3')-(17 RA P)-(17 RA O5') | -160.0 | 20.0 |
| ZETA | (17 RA C3')-(17 RA O3')-(18 RG P)-(18 RG O5') | -160.0 | 20.0 |
| ZETA | (18 RG C3')-(18 RG O3')-(19 RC P)-(19 RC O5') | -160.0 | 20.0 |
| ZETA | (19 RC C3')-(19 RC O3')-(20 RU P)-(20 RU O5') | -160.0 | 20.0 |
| ZETA | (21 RG C3')-(21 RG O3')-(22 RC P)-(22 RC O5') | -160.0 | 20.0 |
| ZETA | (22 RC C3')-(22 RC O3')-(23 RU P)-(23 RU O5') | -160.0 | 20.0 |
| ZETA | (23 RU C3')-(23 RU O3')-(24 RG P)-(24 RG O5') | -160.0 | 20.0 |
| ZETA | (24 RG C3')-(24 RG O3')-(25 RU P)-(25 RU O5') | -160.0 | 20.0 |
| ZETA | (25 RU C3')-(25 RU O3')-(26 RC3 P)-(26 RC3 O5') | -160.0 | 20.0 |
| CHI | (2 RA O4')-(2 RA C1')-(2 RA N9)-(2 RA C4) | 170.0 | 340.0 |
| CHI | (3 RC O4')-(3 RC C1')-(3 RC N1)-(3 RC C2) | 170.0 | 340.0 |
| CHI | (4 RA O4')-(4 RA C1')-(4 RA N9)-(4 RA C4) | 170.0 | 340.0 |
| CHI | (5 RG O4')-(5 RG C1')-(5 RG N9)-(5 RG C4) | 170.0 | 340.0 |
| CHI | (6 RC O4')-(6 RC C1')-(6 RC N1)-(6 RC C2) | 170.0 | 340.0 |
| CHI | (8 RG O4')-(8 RG C1')-(8 RG N9)-(8 RG C4) | 170.0 | 340.0 |
| CHI | (9 RC O4')-(9 RC C1')-(9 RC N1)-(9 RC C2) | 170.0 | 340.0 |
| CHI | (10 RU O4')-(10 RU C1')-(10 RU N1)-(10 RU C2) | 170.0 | 340.0 |
| CHI | (11 RG O4')-(11 RG C1')-(11 RG N9)-(11 RG C4) | 170.0 | 340.0 |
| CHI | (12 RU O4')-(12 RU C1')-(12 RU N1)-(12 RU C2) | 170.0 | 340.0 |
| CHI | (15 RA O4')-(15 RA C1')-(15 RA N9)-(15 RA C4) | 170.0 | 340.0 |
| CHI | (16 RC O4')-(16 RC C1')-(16 RC N1)-(16 RC C2) | 170.0 | 340.0 |
| CHI | (17 RA O4')-(17 RA C1')-(17 RA N9)-(17 RA C4) | 170.0 | 340.0 |
| CHI | (18 RG O4')-(18 RG C1')-(18 RG N9)-(18 RG C4) | 170.0 | 340.0 |
| CHI | (19 RC O4')-(19 RC C1')-(19 RC N1)-(19 RC C2) | 170.0 | 340.0 |
| CHI | (21 RG O4')-(21 RG C1')-(21 RG N9)-(21 RG C4) | 170.0 | 340.0 |
| CHI | (22 RC O4')-(22 RC C1')-(22 RC N1)-(22 RC C2) | 170.0 | 340.0 |
| CHI | (23 RU O4')-(23 RU C1')-(23 RU N1)-(23 RU C2) | 170.0 | 340.0 |
| CHI | (24 RG O4')-(24 RG C1')-(24 RG N9)-(24 RG C4) | 170.0 | 340.0 |
| CHI | (25 RU O4')-(25 RU C1')-(25 RU N1)-(25 RU C2) | 170.0 | 340.0 |

**Table S4:** NOE restraints used for modeling of the r(CUG)-**1** complex.

| | | | | | | | |
|---|---|---|---|---|---|---|---|
| 1 | G | H1 | 26 | C | N3 | 1.84 | 2.04 |
| 1 | G | H1' | 1 | G | H4' | 2.51 | 3.76 |
| 1 | G | H1' | 1 | G | H8 | 2.66 | 3.99 |
| 1 | G | H2' | 1 | G | H1' | 2.36 | 3.53 |
| 1 | G | H2' | 1 | G | H8 | 2.82 | 4.23 |
| 1 | G | H2' | 2 | A | H1' | 3.48 | 5.22 |
| 1 | G | H2' | 2 | A | H8 | 2.26 | 3.39 |
| 1 | G | H21 | 26 | C | O2 | 1.75 | 1.95 |
| 1 | G | H4' | 1 | G | H2' | 2.61 | 3.92 |
| 1 | G | N1 | 26 | C | N3 | 2.85 | 3.05 |
| 1 | G | O6 | 26 | C | H41 | 1.8 | 2 |
| 1 | G | O6 | 26 | C | N4 | 2.81 | 3.01 |
| 2 | A | H1' | 2 | A | H2' | 2.24 | 3.36 |
| 2 | A | H1' | 2 | A | H8 | 2.98 | 4.47 |
| 2 | A | H1' | 3 | C | H1' | 4 | 6.01 |
| 2 | A | H2 | 2 | A | H1' | 3.69 | 5.54 |
| 2 | A | H2 | 2 | A | H2' | 4.29 | 6.43 |
| 2 | A | H2' | 3 | C | H5 | 2.7 | 4.06 |
| 2 | A | H3' | 2 | A | H1' | 2.93 | 4.4 |
| 2 | A | H3' | 2 | A | H8 | 2.62 | 3.93 |
| 2 | A | H3' | 3 | C | H6 | 3.24 | 4.86 |
| 2 | A | H61 | 25 | U | O4 | 1.84 | 2.04 |
| 2 | A | H8 | 2 | A | H2' | 3.09 | 4.64 |
| 2 | A | H8 | 3 | C | H5 | 3.46 | 5.18 |
| 2 | A | H8 | 3 | C | H6 | 3.5 | 5.24 |
| 2 | A | N1 | 25 | U | H3 | 1.71 | 1.91 |
| 2 | A | N1 | 25 | U | N3 | 2.72 | 2.92 |
| 3 | C | H1' | 2 | A | H2 | 2.69 | 4.04 |
| 3 | C | H1' | 3 | C | H5 | 4.09 | 6.13 |
| 3 | C | H1' | 4 | A | H1' | 4.23 | 6.35 |
| 3 | C | H2' | 3 | C | H1' | 2.17 | 3.25 |
| 3 | C | H2' | 3 | C | H5 | 3.62 | 5.43 |
| 3 | C | H2' | 3 | C | H6 | 3.47 | 5.21 |
| 3 | C | H2' | 4 | A | H8 | 2.02 | 3.03 |
| 3 | C | H3' | 4 | A | H8 | 2.39 | 3.58 |
| 3 | C | H41 | 24 | G | O6 | 1.8 | 2 |
| 3 | C | H5 | 2 | A | H3' | 2.83 | 4.25 |
| 3 | C | H5 | 3 | C | H6 | 2.04 | 3.06 |
| 3 | C | H6 | 2 | A | H2' | 1.91 | 2.86 |
| 3 | C | H6 | 3 | C | H1' | 2.58 | 3.87 |
| 3 | C | N3 | 24 | G | H1 | 1.84 | 2.04 |
| 3 | C | N3 | 24 | G | N1 | 2.85 | 3.05 |
| 3 | C | N4 | 24 | G | O6 | 2.81 | 3.01 |
| 3 | C | O2 | 24 | G | H21 | 1.75 | 1.95 |
| 4 | A | H1' | 3 | C | H2' | 3.17 | 4.76 |
| 4 | A | H1' | 4 | A | H3' | 3.12 | 4.68 |
| 4 | A | H1' | 4 | A | H8 | 3.01 | 4.51 |
| 4 | A | H2 | 4 | A | H1' | 3.84 | 5.76 |
| 4 | A | H2 | 5 | G | H1' | 2.68 | 4.01 |

| | | | | | | | |
|---|---|---|---|---|---|---|---|
| 4 | A | H2' | 4 | A | H1' | 2.32 | 3.48 |
| 4 | A | H2' | 4 | A | H8 | 2.87 | 4.31 |
| 4 | A | H2' | 5 | G | H1' | 3.45 | 5.18 |
| 4 | A | H2' | 5 | G | H8 | 2.25 | 3.37 |
| 4 | A | H3' | 4 | A | H8 | 2.69 | 4.04 |
| 4 | A | H3' | 5 | G | H8 | 3.01 | 4.52 |
| 4 | A | H4' | 4 | A | H1' | 2.61 | 3.91 |
| 4 | A | H61 | 23 | U | O4 | 1.84 | 2.04 |
| 4 | A | H8 | 3 | C | H5 | 3.73 | 5.59 |
| 4 | A | H8 | 3 | C | H6 | 3.27 | 4.9 |
| 4 | A | H8 | 5 | G | H8 | 3.57 | 5.35 |
| 4 | A | N1 | 23 | U | H3 | 1.71 | 1.91 |
| 4 | A | N1 | 23 | U | N3 | 2.72 | 2.92 |
| 5 | G | H1 | 22 | C | N3 | 1.84 | 2.04 |
| 5 | G | H1' | 5 | G | H2' | 2.12 | 3.19 |
| 5 | G | H1' | 5 | G | H8 | 2.87 | 4.31 |
| 5 | G | H2' | 5 | G | H8 | 2.87 | 4.3 |
| 5 | G | H2' | 6 | C | H1' | 3.2 | 4.8 |
| 5 | G | H2' | 6 | C | H5 | 2.7 | 4.07 |
| 5 | G | H2' | 6 | C | H6 | 2.17 | 3.25 |
| 5 | G | H21 | 22 | C | O2 | 1.75 | 1.95 |
| 5 | G | H3' | 6 | C | H6 | 2.3 | 3.45 |
| 5 | G | H8 | 5 | G | H3' | 2.24 | 3.36 |
| 5 | G | N1 | 22 | C | N3 | 2.85 | 3.05 |
| 5 | G | O6 | 22 | C | H41 | 1.8 | 2 |
| 5 | G | O6 | 22 | C | N4 | 2.81 | 3.01 |
| 6 | C | H1' | 6 | C | H5 | 3.84 | 5.76 |
| 6 | C | H1' | 6 | C | H6 | 2.72 | 4.08 |
| 6 | C | H2' | 6 | C | H1' | 2.13 | 3.2 |
| 6 | C | H2' | 6 | C | H6 | 2.9 | 4.35 |
| 6 | C | H2' | 7 | U | H1' | 2.25 | 3.38 |
| 6 | C | H3' | 7 | U | H6 | 2.37 | 3.56 |
| 6 | C | H41 | 21 | G | O6 | 1.8 | 2 |
| 6 | C | H5 | 5 | G | H8 | 3.15 | 4.72 |
| 6 | C | H5 | 6 | C | H2' | 3.79 | 5.69 |
| 6 | C | H5 | 6 | C | H6 | 2.03 | 3.04 |
| 6 | C | H5 | 7 | U | H6 | 3.28 | 4.92 |
| 6 | C | H6 | 5 | G | H8 | 3.59 | 5.38 |
| 6 | C | H6 | 6 | C | H3' | 1.93 | 2.9 |
| 6 | C | N3 | 21 | G | H1 | 1.84 | 2.04 |
| 6 | C | N3 | 21 | G | N1 | 2.85 | 3.05 |
| 6 | C | N4 | 21 | G | O6 | 2.81 | 3.01 |
| 6 | C | O2 | 21 | G | H21 | 1.75 | 1.95 |
| 7 | U | H1' | 7 | U | H2' | 2.19 | 3.28 |
| 7 | U | H1' | 7 | U | H6 | 2.42 | 3.63 |
| 7 | U | H5 | 6 | C | H2' | 2.74 | 4.11 |
| 7 | U | H5 | 6 | C | H5 | 2.92 | 4.38 |
| 7 | U | H5 | 6 | C | H6 | 3.37 | 5.06 |
| 7 | U | H5 | 7 | U | H6 | 1.95 | 2.92 |
| 7 | U | H6 | 6 | C | H2' | 2.2 | 3.3 |
| 7 | U | H6 | 6 | C | H6 | 3.74 | 5.6 |

| | | | | | | | |
|---|---|---|---|---|---|---|---|
| 7 | U | H6 | 8 | G | H8 | 2.88 | 5.05 |
| 8 | G | H1 | 19 | C | N3 | 1.84 | 2.04 |
| 8 | G | H1' | 7 | U | H2' | 3.02 | 4.52 |
| 8 | G | H1' | 8 | G | H8 | 2.87 | 4.31 |
| 8 | G | H2' | 8 | G | H1' | 2.5 | 3.76 |
| 8 | G | H2' | 8 | G | H8 | 3.44 | 5.15 |
| 8 | G | H2' | 9 | C | H1' | 3.34 | 5.01 |
| 8 | G | H2' | 9 | C | H5 | 2.86 | 4.29 |
| 8 | G | H21 | 19 | C | O2 | 1.75 | 1.95 |
| 8 | G | H3' | 8 | G | H8 | 3.23 | 4.85 |
| 8 | G | H8 | 7 | U | H2' | 2.14 | 3.21 |
| 8 | G | H8 | 7 | U | H6 | 3.34 | 5.01 |
| 8 | G | N1 | 19 | C | N3 | 2.85 | 3.05 |
| 8 | G | O6 | 19 | C | H41 | 1.8 | 2 |
| 8 | G | O6 | 19 | C | N4 | 2.81 | 3.01 |
| 9 | C | H1' | 9 | C | H2' | 2.19 | 3.28 |
| 9 | C | H1' | 9 | C | H5 | 3.93 | 5.89 |
| 9 | C | H1' | 9 | C | H6 | 2.93 | 4.39 |
| 9 | C | H2' | 10 | U | H5 | 2.71 | 4.06 |
| 9 | C | H2' | 10 | U | H6 | 2.16 | 3.25 |
| 9 | C | H41 | 18 | G | O6 | 1.8 | 2 |
| 9 | C | H5 | 8 | G | H8 | 3.34 | 5.01 |
| 9 | C | H5 | 9 | C | H3' | 3.14 | 4.71 |
| 9 | C | H5 | 9 | C | H6 | 2.15 | 3.23 |
| 9 | C | H5 | 10 | U | H5 | 2.97 | 4.45 |
| 9 | C | H6 | 9 | C | H2' | 3.26 | 4.89 |
| 9 | C | H6 | 10 | U | H5 | 3.6 | 5.4 |
| 9 | C | N3 | 18 | G | H1 | 1.84 | 2.04 |
| 9 | C | N3 | 18 | G | N1 | 2.85 | 3.05 |
| 9 | C | N4 | 18 | G | O6 | 2.81 | 3.01 |
| 9 | C | O2 | 18 | G | H21 | 1.75 | 1.95 |
| 10 | U | H3 | 17 | A | N1 | 1.71 | 1.91 |
| 10 | U | H5 | 9 | C | H3' | 3.35 | 5.02 |
| 10 | U | H6 | 9 | C | H3' | 2.96 | 4.44 |
| 10 | U | H6 | 9 | C | H6 | 3.4 | 5.1 |
| 10 | U | H6 | 10 | U | H1' | 2.68 | 4.03 |
| 10 | U | H6 | 10 | U | H5 | 2.15 | 3.22 |
| 10 | U | N3 | 17 | A | N1 | 2.72 | 2.92 |
| 10 | U | O4 | 17 | A | H61 | 1.84 | 2.04 |
| 11 | G | H1 | 16 | C | N3 | 1.84 | 2.04 |
| 11 | G | H1' | 11 | G | H3' | 2.6 | 3.89 |
| 11 | G | H1' | 12 | U | H5 | 3.94 | 5.91 |
| 11 | G | H2' | 11 | G | H1' | 2.21 | 3.32 |
| 11 | G | H21 | 16 | C | O2 | 1.75 | 1.95 |
| 11 | G | H3' | 12 | U | H5 | 2.92 | 4.39 |
| 11 | G | H8 | 10 | U | H1' | 3.4 | 5.1 |
| 11 | G | H8 | 11 | G | H1' | 3.39 | 5.08 |
| 11 | G | H8 | 11 | G | H2' | 3.82 | 5.72 |
| 11 | G | H8 | 11 | G | H3' | 2.67 | 4.01 |
| 11 | G | H8 | 12 | U | H5 | 3.84 | 5.75 |
| 11 | G | N1 | 16 | C | N3 | 2.85 | 3.05 |

| | | | | | | | |
|---|---|---|---|---|---|---|---|
| 11 | G | O6 | 16 | C | H41 | 1.8 | 2 |
| 11 | G | O6 | 16 | C | N4 | 2.81 | 3.01 |
| 12 | U | H1' | 11 | G | H2' | 3.07 | 4.61 |
| 12 | U | H1' | 12 | U | H2' | 2.07 | 3.1 |
| 12 | U | H1' | 12 | U | H6 | 2.77 | 4.15 |
| 12 | U | H2' | 13 | C | H5 | 2.81 | 4.22 |
| 12 | U | H2' | 13 | C | H6 | 2 | 3 |
| 12 | U | H3 | 15 | A | N1 | 1.71 | 1.91 |
| 12 | U | H3' | 12 | U | H1' | 2.67 | 4.01 |
| 12 | U | H3' | 13 | C | H6 | 3.34 | 5.02 |
| 12 | U | H5 | 11 | G | H2' | 2.52 | 3.79 |
| 12 | U | H5 | 12 | U | H2' | 3.76 | 5.64 |
| 12 | U | H5 | 12 | U | H6 | 2.01 | 3.02 |
| 12 | U | H5 | 13 | C | H5 | 2.86 | 4.29 |
| 12 | U | H6 | 11 | G | H2' | 1.99 | 2.99 |
| 12 | U | H6 | 11 | G | H3' | 2.3 | 3.45 |
| 12 | U | H6 | 12 | U | H2' | 2.75 | 4.12 |
| 12 | U | H6 | 12 | U | H3' | 2.04 | 3.06 |
| 12 | U | H6 | 13 | C | H5 | 3.39 | 5.09 |
| 12 | U | N3 | 15 | A | N1 | 2.72 | 2.92 |
| 12 | U | O4 | 15 | A | H61 | 1.84 | 2.04 |
| 13 | C | H1' | 12 | U | H2' | 2.85 | 4.27 |
| 13 | C | H1' | 13 | C | H2' | 1.98 | 2.97 |
| 13 | C | H1' | 13 | C | H6 | 2.98 | 4.47 |
| 13 | C | H2' | 13 | C | H6 | 2.67 | 4.01 |
| 13 | C | H6 | 13 | C | H3' | 1.86 | 2.79 |
| 14 | G | H1 | 13 | C | N3 | 1.84 | 2.04 |
| 14 | G | H1' | 14 | G | H4' | 2.51 | 3.76 |
| 14 | G | H1' | 14 | G | H8 | 2.66 | 3.99 |
| 14 | G | H2' | 14 | G | H1' | 2.36 | 3.53 |
| 14 | G | H2' | 14 | G | H8 | 2.82 | 4.23 |
| 14 | G | H2' | 15 | A | H1' | 3.48 | 5.22 |
| 14 | G | H2' | 15 | A | H8 | 2.26 | 3.39 |
| 14 | G | H21 | 13 | C | O2 | 1.75 | 1.95 |
| 14 | G | H4' | 14 | G | H2' | 2.61 | 3.92 |
| 14 | G | N1 | 13 | C | N3 | 2.85 | 3.05 |
| 14 | G | O6 | 13 | C | H41 | 1.8 | 2 |
| 14 | G | O6 | 13 | C | N4 | 2.81 | 3.01 |
| 15 | A | H1' | 15 | A | H2' | 2.24 | 3.36 |
| 15 | A | H1' | 15 | A | H8 | 2.98 | 4.47 |
| 15 | A | H1' | 16 | C | H1' | 4 | 6.01 |
| 15 | A | H2 | 15 | A | H1' | 3.69 | 5.54 |
| 15 | A | H2 | 15 | A | H2' | 4.29 | 6.43 |
| 15 | A | H2' | 16 | C | H5 | 2.7 | 4.06 |
| 15 | A | H3' | 15 | A | H1' | 2.93 | 4.4 |
| 15 | A | H3' | 15 | A | H8 | 2.62 | 3.93 |
| 15 | A | H3' | 16 | C | H6 | 3.24 | 4.86 |
| 15 | A | H8 | 15 | A | H2' | 3.09 | 4.64 |
| 15 | A | H8 | 16 | C | H5 | 3.46 | 5.18 |
| 15 | A | H8 | 16 | C | H6 | 3.5 | 5.24 |
| 16 | C | H1' | 15 | A | H2 | 2.69 | 4.04 |

| | | | | | | | |
|---|---|---|---|---|---|---|---|
| 16 | C | H1' | 16 | C | H5 | 4.09 | 6.13 |
| 16 | C | H1' | 17 | A | H1' | 4.23 | 6.35 |
| 16 | C | H2' | 16 | C | H1' | 2.17 | 3.25 |
| 16 | C | H2' | 16 | C | H5 | 3.62 | 5.43 |
| 16 | C | H2' | 16 | C | H6 | 3.47 | 5.21 |
| 16 | C | H2' | 17 | A | H8 | 2.02 | 3.03 |
| 16 | C | H3' | 17 | A | H8 | 2.39 | 3.58 |
| 16 | C | H5 | 15 | A | H3' | 2.83 | 4.25 |
| 16 | C | H5 | 16 | C | H6 | 2.04 | 3.06 |
| 16 | C | H6 | 15 | A | H2' | 1.91 | 2.86 |
| 16 | C | H6 | 16 | C | H1' | 2.58 | 3.87 |
| 17 | A | H1' | 16 | C | H2' | 3.17 | 4.76 |
| 17 | A | H1' | 17 | A | H3' | 3.12 | 4.68 |
| 17 | A | H1' | 17 | A | H8 | 3.01 | 4.51 |
| 17 | A | H2 | 17 | A | H1' | 3.84 | 5.76 |
| 17 | A | H2 | 18 | G | H1' | 2.68 | 4.01 |
| 17 | A | H2' | 17 | A | H1' | 2.32 | 3.48 |
| 17 | A | H2' | 17 | A | H8 | 2.87 | 4.31 |
| 17 | A | H2' | 18 | G | H1' | 3.45 | 5.18 |
| 17 | A | H2' | 18 | G | H8 | 2.25 | 3.37 |
| 17 | A | H3' | 17 | A | H8 | 2.69 | 4.04 |
| 17 | A | H3' | 18 | G | H8 | 3.01 | 4.52 |
| 17 | A | H4' | 17 | A | H1' | 2.61 | 3.91 |
| 17 | A | H8 | 16 | C | H5 | 3.73 | 5.59 |
| 17 | A | H8 | 16 | C | H6 | 3.27 | 4.9 |
| 17 | A | H8 | 18 | G | H8 | 3.57 | 5.35 |
| 18 | G | H1' | 18 | G | H2' | 2.12 | 3.19 |
| 18 | G | H1' | 18 | G | H8 | 2.87 | 4.31 |
| 18 | G | H2' | 18 | G | H8 | 2.87 | 4.3 |
| 18 | G | H2' | 19 | C | H1' | 3.2 | 4.8 |
| 18 | G | H2' | 19 | C | H5 | 2.7 | 4.06 |
| 18 | G | H2' | 19 | C | H6 | 2.17 | 3.25 |
| 18 | G | H3' | 19 | C | H6 | 2.3 | 3.45 |
| 18 | G | H8 | 18 | G | H3' | 2.24 | 3.36 |
| 19 | C | H1' | 19 | C | H5 | 3.84 | 5.76 |
| 19 | C | H1' | 19 | C | H6 | 2.72 | 4.08 |
| 19 | C | H2' | 19 | C | H1' | 2.13 | 3.2 |
| 19 | C | H2' | 19 | C | H6 | 2.9 | 4.35 |
| 19 | C | H2' | 20 | U | H1' | 2.25 | 3.38 |
| 19 | C | H3' | 20 | U | H6 | 2.37 | 3.56 |
| 19 | C | H5 | 18 | G | H8 | 3.15 | 4.72 |
| 19 | C | H5 | 19 | C | H2' | 3.79 | 5.69 |
| 19 | C | H5 | 19 | C | H6 | 2.03 | 3.04 |
| 19 | C | H5 | 20 | U | H6 | 3.28 | 4.92 |
| 19 | C | H6 | 18 | G | H8 | 3.59 | 5.38 |
| 19 | C | H6 | 19 | C | H3' | 1.93 | 2.9 |
| 20 | U | H1' | 20 | U | H2' | 2.19 | 3.28 |
| 20 | U | H1' | 20 | U | H6 | 2.42 | 3.63 |
| 20 | U | H5 | 19 | C | H2' | 2.74 | 4.11 |
| 20 | U | H5 | 19 | C | H5 | 2.92 | 4.38 |
| 20 | U | H5 | 19 | C | H6 | 3.37 | 5.06 |

| | | | | | | | |
|---|---|---|---|---|---|---|---|
| 20 | U | H5 | 20 | U | H6 | 1.95 | 2.92 |
| 20 | U | H6 | 19 | C | H2' | 2.2 | 3.3 |
| 20 | U | H6 | 19 | C | H6 | 3.74 | 5.6 |
| 20 | U | H6 | 21 | G | H8 | 2.88 | 5.05 |
| 21 | G | H1' | 20 | U | H2' | 3.02 | 4.52 |
| 21 | G | H1' | 21 | G | H8 | 2.87 | 4.31 |
| 21 | G | H2' | 21 | G | H1' | 2.5 | 3.76 |
| 21 | G | H2' | 21 | G | H8 | 3.44 | 5.15 |
| 21 | G | H2' | 22 | C | H1' | 3.34 | 5.01 |
| 21 | G | H2' | 22 | C | H6 | 2.91 | 4.36 |
| 21 | G | H3' | 21 | G | H8 | 3.23 | 4.85 |
| 21 | G | H8 | 20 | U | H2' | 2.14 | 3.21 |
| 21 | G | H8 | 20 | U | H6 | 3.34 | 5.01 |
| 22 | C | H1' | 22 | C | H2' | 2.19 | 3.28 |
| 22 | C | H1' | 22 | C | H5 | 3.93 | 5.89 |
| 22 | C | H1' | 22 | C | H6 | 2.93 | 4.39 |
| 22 | C | H2' | 23 | U | H5 | 2.71 | 4.06 |
| 22 | C | H2' | 23 | U | H6 | 2.16 | 3.25 |
| 22 | C | H5 | 21 | G | H8 | 3.34 | 5.01 |
| 22 | C | H5 | 22 | C | H3' | 3.14 | 4.71 |
| 22 | C | H5 | 22 | C | H6 | 2.15 | 3.23 |
| 22 | C | H5 | 23 | U | H5 | 2.97 | 4.45 |
| 22 | C | H6 | 22 | C | H2' | 3.26 | 4.89 |
| 22 | C | H6 | 23 | U | H5 | 3.6 | 5.4 |
| 23 | U | H5 | 22 | C | H3' | 3.35 | 5.02 |
| 23 | U | H6 | 22 | C | H3' | 2.96 | 4.44 |
| 23 | U | H6 | 22 | C | H6 | 3.4 | 5.1 |
| 23 | U | H6 | 23 | U | H1' | 2.68 | 4.03 |
| 23 | U | H6 | 23 | U | H5 | 2.15 | 3.22 |
| 24 | G | H1' | 24 | G | H3' | 2.6 | 3.89 |
| 24 | G | H1' | 25 | U | H5 | 3.94 | 5.91 |
| 24 | G | H2' | 24 | G | H1' | 2.21 | 3.32 |
| 24 | G | H3' | 25 | U | H5 | 2.92 | 4.39 |
| 24 | G | H8 | 23 | U | H1' | 3.4 | 5.1 |
| 24 | G | H8 | 24 | G | H1' | 3.39 | 5.08 |
| 24 | G | H8 | 24 | G | H2' | 3.82 | 5.72 |
| 24 | G | H8 | 24 | G | H3' | 2.67 | 4.01 |
| 24 | G | H8 | 25 | U | H5 | 3.84 | 5.75 |
| 25 | U | H1' | 24 | G | H2' | 3.07 | 4.61 |
| 25 | U | H1' | 25 | U | H2' | 2.07 | 3.1 |
| 25 | U | H1' | 25 | U | H6 | 2.77 | 4.15 |
| 25 | U | H2' | 26 | C | H5 | 2.81 | 4.22 |
| 25 | U | H2' | 26 | C | H6 | 2 | 3 |
| 25 | U | H3' | 25 | U | H1' | 2.67 | 4.01 |
| 25 | U | H3' | 26 | C | H6 | 3.34 | 5.02 |
| 25 | U | H5 | 24 | G | H2' | 2.52 | 3.79 |
| 25 | U | H5 | 25 | U | H2' | 3.76 | 5.64 |
| 25 | U | H5 | 25 | U | H6 | 2.01 | 3.02 |
| 25 | U | H5 | 26 | C | H5 | 2.86 | 4.29 |
| 25 | U | H6 | 24 | G | H2' | 1.99 | 2.99 |
| 25 | U | H6 | 24 | G | H3' | 2.3 | 3.45 |

| | | | | | | | |
|---|---|---|---|---|---|---|---|
| 25 | U | H6 | 25 | U | H2' | 2.75 | 4.12 |
| 25 | U | H6 | 25 | U | H3' | 2.04 | 3.06 |
| 25 | U | H6 | 26 | C | H5 | 3.39 | 5.09 |
| 26 | C | H1' | 25 | U | H2' | 2.85 | 4.27 |
| 26 | C | H1' | 26 | C | H2' | 1.98 | 2.97 |
| 26 | C | H1' | 26 | C | H6 | 2.98 | 4.47 |
| 26 | C | H2' | 26 | C | H6 | 2.67 | 4.01 |
| 26 | C | H6 | 26 | C | H3' | 1.86 | 2.79 |
| 27 | **1** | H1 | 6 | C | H2' | 3 | 6 |
| 27 | **1** | H1 | 6 | C | H5 | 3 | 6 |
| 27 | **1** | H1 | 7 | U | H1' | 3 | 6 |
| 27 | **1** | H1 | 7 | U | H5 | 3 | 6 |
| 27 | **1** | H1 | 7 | U | H6 | 3 | 6 |
| 27 | **1** | H1 | 8 | G | H1' | 3 | 6 |
| 27 | **1** | H1 | 8 | G | H8 | 3 | 6 |
| 27 | **1** | H3 | 6 | C | H2' | 3 | 6 |
| 27 | **1** | H3 | 6 | C | H5 | 3 | 6 |
| 27 | **1** | H3 | 7 | U | H5 | 3 | 6 |
| 27 | **1** | H3 | 7 | U | H6 | 3 | 6 |
| 27 | **1** | H3 | 8 | G | H1' | 3 | 6 |
| 27 | **1** | H3 | 8 | G | H8 | 3 | 6 |
| 27 | **1** | H5 | 19 | C | H2' | 3 | 6 |
| 27 | **1** | H5 | 20 | U | H1' | 3 | 6 |
| 27 | **1** | H5 | 20 | U | H5 | 3 | 6 |
| 27 | **1** | H5 | 20 | U | H6 | 3 | 6 |
| 27 | **1** | H5 | 21 | G | H1' | 3 | 6 |
| 27 | **1** | H5 | 21 | G | H8 | 3 | 6 |
| 27 | **1** | H7 | 20 | U | H1' | 3 | 6 |
| 27 | **1** | H7 | 20 | U | H5 | 3 | 6 |
| 27 | **1** | H7 | 20 | U | H6 | 3 | 6 |
| 27 | **1** | H7 | 21 | G | H1' | 3 | 6 |
| 27 | **1** | H7 | 21 | G | H8 | 3 | 6 |

**Table S5:** Distance restraint violations greater than 0.1 Å for the r(CUG)-**1** complex.

| No. of violations ≥ 0.1 Å | Mean violation† (Å) | Assignment | Minimum distance (Å) | Maximum distance (Å) |
|---|---|---|---|---|
| 10 | 0.095 | **1**27H3-C6H2′ | 3.00 | 6.00 |
| 1 | 0.052 | C19H2′-U20H1′ | 2.25 | 3.38 |

† Includes violations < 0.1 Å

**Table S6:** NOE restraints used for modeling of the r(CUG)-**2** complex.

| | | | | | | | |
|---|---|---|---|---|---|---|---|
| 1 | G | H1 | 2 | A | H1' | 3.32 | 5.19 |
| 1 | G | H1 | 2 | A | H2 | 3.42 | 5.34 |
| 1 | G | H1 | 26 | C | H42 | 3.4 | 5.31 |
| 1 | G | H1 | 26 | C | N3 | 1.8 | 2.4 |
| 1 | G | H1' | 1 | G | H2' | 2.12 | 3.46 |
| 1 | G | H1' | 1 | G | H8 | 2.7 | 4.22 |
| 1 | G | H22 | 26 | C | O2 | 1.8 | 2.4 |
| 1 | G | H3' | 1 | G | H1' | 2.58 | 3.86 |
| 1 | G | H8 | 1 | G | H2' | 2.8 | 4.38 |
| 1 | G | O6 | 26 | C | H41 | 1.8 | 2.4 |
| 2 | A | H1' | 2 | A | H2' | 2.15 | 3.23 |
| 2 | A | H1' | 2 | A | H3' | 2.51 | 3.99 |
| 2 | A | H1' | 2 | A | H4' | 2.77 | 4.41 |
| 2 | A | H1' | 2 | A | H4' | 2.77 | 4.41 |
| 2 | A | H2 | 2 | A | H1' | 3 | 6 |
| 2 | A | H2 | 2 | A | H1' | 3 | 6 |
| 2 | A | H2 | 3 | C | H1' | 2.57 | 3.88 |
| 2 | A | H2 | 3 | C | H1' | 2.84 | 4.52 |
| 2 | A | H2 | 26 | C | H1' | 3.12 | 4.88 |
| 2 | A | H61 | 25 | U | O4 | 1.8 | 2.4 |
| 2 | A | H8 | 1 | G | H1' | 3.04 | 4.75 |
| 2 | A | H8 | 1 | G | H2' | 1.6 | 2.94 |
| 2 | A | H8 | 2 | A | H1' | 2.88 | 4.48 |
| 2 | A | H8 | 2 | A | H2' | 2.75 | 4.25 |
| 2 | A | H8 | 2 | A | H3' | 2 | 3.29 |
| 2 | A | H8 | 3 | C | H5 | 2.96 | 4.63 |
| 2 | A | N1 | 25 | U | H3 | 1.8 | 2.4 |
| 3 | C | H1' | 3 | C | H2' | 2.07 | 3.39 |
| 3 | C | H1' | 3 | C | H4' | 2.83 | 9.22 |
| 3 | C | H41 | 24 | G | O6 | 1.8 | 2.4 |
| 3 | C | H5 | 2 | A | H2' | 3.04 | 4.75 |
| 3 | C | H5 | 3 | C | H3' | 2.88 | 4.48 |
| 3 | C | H6 | 2 | A | H1' | 3.84 | 5.25 |
| 3 | C | H6 | 2 | A | H2' | 1.7 | 3 |
| 3 | C | H6 | 2 | A | H3' | 2.53 | 8.85 |
| 3 | C | H6 | 3 | C | H1' | 2.74 | 4.17 |
| 3 | C | H6 | 3 | C | H2' | 2.75 | 4.1 |
| 3 | C | H6 | 3 | C | H4' | 2.78 | 4.33 |
| 3 | C | H6 | 4 | A | H8 | 2.8 | 4.375 |
| 3 | C | N3 | 24 | G | H1 | 1.8 | 2.4 |
| 3 | C | O2 | 24 | G | H22 | 1.8 | 2.4 |
| 4 | A | H1' | 4 | A | H2' | 2.26 | 3.63 |
| 4 | A | H2 | 4 | A | H1' | 3.04 | 4.75 |
| 4 | A | H2 | 5 | G | H1' | 2.96 | 4.63 |
| 4 | A | H2 | 24 | G | H1' | 2.72 | 3.89 |
| 4 | A | H61 | 23 | U | O4 | 1.8 | 2.4 |
| 4 | A | H8 | 3 | C | H1' | 2.88 | 4.5 |
| 4 | A | H8 | 3 | C | H2' | 1.88 | 3.34 |
| 4 | A | H8 | 3 | C | H3' | 2.88 | 4.5 |

| | | | | | | | |
|---|---|---|---|---|---|---|---|
| 4 | A | H8 | 4 | A | H1' | 2.75 | 4.1 |
| 4 | A | H8 | 4 | A | H2' | 3.04 | 4.75 |
| 4 | A | H8 | 4 | A | H3' | 2.72 | 4.25 |
| 4 | A | N1 | 23 | U | H3 | 1.8 | 2.4 |
| 5 | G | H1 | 22 | C | H41 | 2.78 | 4.34 |
| 5 | G | H1 | 22 | C | N3 | 1.8 | 2.4 |
| 5 | G | H1 | 23 | U | H1' | 2.74 | 4.28 |
| 5 | G | H1' | 5 | G | H2' | 2.11 | 3.43 |
| 5 | G | H1' | 5 | G | H3' | 2.57 | 3.78 |
| 5 | G | H2' | 5 | G | H1' | 2.14 | 3.51 |
| 5 | G | H2' | 6 | C | H5 | 2.94 | 4.55 |
| 5 | G | H22 | 22 | C | O2 | 1.8 | 2.4 |
| 5 | G | H3' | 6 | C | H5 | 2.65 | 3.94 |
| 5 | G | H3' | 6 | C | H5 | 2.65 | 3.99 |
| 5 | G | H4' | 5 | G | H1' | 2.56 | 4.06 |
| 5 | G | H8 | 4 | A | H1' | 3.12 | 4.875 |
| 5 | G | H8 | 4 | A | H2' | 1.95 | 3.36 |
| 5 | G | H8 | 4 | A | H3' | 1.92 | 3 |
| 5 | G | H8 | 5 | G | H1' | 2.77 | 4.09 |
| 5 | G | H8 | 5 | G | H2' | 2.74 | 4.42 |
| 5 | G | H8 | 5 | G | H3' | 2.08 | 3.25 |
| 5 | G | H8 | 6 | C | H5 | 2.75 | 3.91 |
| 5 | G | O6 | 22 | C | H41 | 1.8 | 2.4 |
| 6 | C | H1' | 5 | G | H2' | 3.21 | 4.78 |
| 6 | C | H1' | 6 | C | H3' | 2.8 | 4.38 |
| 6 | C | H3' | 6 | C | H5 | 2.96 | 4.63 |
| 6 | C | H4' | 6 | C | H1' | 2.18 | 3.91 |
| 6 | C | H6 | 5 | G | H1' | 3.2 | 5 |
| 6 | C | H6 | 5 | G | H2' | 1.83 | 2.9 |
| 6 | C | H6 | 5 | G | H3' | 2 | 3.06 |
| 6 | C | H6 | 5 | G | H8 | 3.84 | 5.25 |
| 6 | C | H6 | 6 | C | H1' | 2.96 | 4.03 |
| 6 | C | H6 | 6 | C | H2' | 2.75 | 4.1 |
| 6 | C | H6 | 6 | C | H3' | 2 | 3.13 |
| 6 | C | H6 | 6 | C | H4' | 2.8 | 4.38 |
| 7 | U | H1' | 7 | U | H6 | 2.8 | 4.38 |
| 7 | U | H6 | 6 | C | H1' | 4.4 | 5.5 |
| 7 | U | H6 | 6 | C | H2' | 2.48 | 3.88 |
| 7 | U | H6 | 6 | C | H3' | 3 | 5 |
| 7 | U | H6 | 7 | U | H1' | 2.8 | 4.17 |
| 7 | U | H6 | 7 | U | H2' | 2.8 | 4.17 |
| 8 | G | H4' | 8 | G | H1' | 2.64 | 4.13 |
| 8 | G | H8 | 7 | U | H2' | 2.32 | 3.63 |
| 8 | G | H8 | 8 | G | H1' | 2.8 | 4.45 |
| 8 | G | H8 | 8 | G | H2' | 2.22 | 3.71 |
| 9 | C | H1' | 9 | C | H2' | 2.1 | 3.55 |
| 9 | C | H1' | 9 | C | H3' | 2.22 | 4.4 |
| 9 | C | H2' | 10 | U | H5 | 2.91 | 4.28 |
| 9 | C | H3' | 10 | U | H5 | 2.72 | 4.1 |
| 9 | C | H41 | 18 | G | O6 | 1.8 | 2.4 |
| 9 | C | H6 | 8 | G | H1' | 2.72 | 4.25 |

| | | | | | | | |
|---|---|---|---|---|---|---|---|
| 9 | C | H6 | 9 | C | H2' | 2.8 | 4.63 |
| 9 | C | H6 | 9 | C | H3' | 2.09 | 3.49 |
| 9 | C | H6 | 10 | U | H5 | 3.12 | 4.88 |
| 9 | C | N3 | 18 | G | H1 | 1.8 | 2.4 |
| 9 | C | O2 | 18 | G | H22 | 1.8 | 2.4 |
| 10 | U | H3 | 11 | G | H1 | 3.69 | 5.77 |
| 10 | U | H3 | 17 | A | H2 | 2.28 | 3.56 |
| 10 | U | H3 | 17 | A | N1 | 1.8 | 2.4 |
| 10 | U | H3 | 18 | G | H1 | 2.28 | 3.56 |
| 10 | U | H6 | 9 | C | H1' | 3.04 | 5 |
| 10 | U | H6 | 9 | C | H2' | 1.85 | 3.19 |
| 10 | U | H6 | 9 | C | H3' | 2.19 | 3.36 |
| 10 | U | H6 | 10 | U | H1' | 3.6 | 5.57 |
| 10 | U | H6 | 11 | G | H8 | 3.04 | 4.75 |
| 10 | U | O4 | 17 | A | H61 | 1.8 | 2.4 |
| 11 | G | H1 | 15 | A | H2 | 2.64 | 4.13 |
| 11 | G | H1 | 16 | C | H41 | 2.65 | 4.14 |
| 11 | G | H1 | 16 | C | N3 | 1.8 | 2.4 |
| 11 | G | H1' | 11 | G | H2' | 2.22 | 3.71 |
| 11 | G | H1' | 11 | G | H4' | 2.64 | 3.93 |
| 11 | G | H22 | 16 | C | O2 | 1.8 | 2.4 |
| 11 | G | H8 | 10 | U | H1' | 3.69 | 5.77 |
| 11 | G | H8 | 11 | G | H1' | 2.96 | 4.03 |
| 11 | G | H8 | 11 | G | H2' | 2.64 | 4.13 |
| 11 | G | H8 | 11 | G | H3' | 2 | 3.29 |
| 11 | G | O6 | 16 | C | H41 | 1.8 | 2.4 |
| 12 | U | H1' | 11 | G | H2' | 3.41 | 5.41 |
| 12 | U | H1' | 12 | U | H2' | 2.09 | 3.51 |
| 12 | U | H1' | 12 | U | H3' | 2.99 | 4.53 |
| 12 | U | H3 | 11 | G | H1 | 2.92 | 4.56 |
| 12 | U | H3 | 12 | U | H1' | 3.44 | 5.38 |
| 12 | U | H3 | 12 | U | H2' | 3.55 | 5.55 |
| 12 | U | H3 | 15 | A | H2 | 2.12 | 3.31 |
| 12 | U | H3 | 15 | A | N1 | 1.8 | 2.4 |
| 12 | U | H3 | 16 | C | H41 | 3.26 | 5.1 |
| 12 | U | H4' | 12 | U | H1' | 2.79 | 4.41 |
| 12 | U | H6 | 11 | G | H2' | 1.75 | 2.92 |
| 12 | U | H6 | 11 | G | H3' | 2 | 3.53 |
| 12 | U | H6 | 12 | U | H1' | 3.6 | 5.57 |
| 12 | U | H6 | 12 | U | H3' | 2 | 3.18 |
| 12 | U | O4 | 15 | A | H61 | 1.8 | 2.4 |
| 13 | C | H1' | 12 | U | H2' | 3.29 | 4.73 |
| 13 | C | H41 | 14 | G | O6 | 1.8 | 2.4 |
| 13 | C | H5 | 12 | U | H2' | 3.68 | 4.6 |
| 13 | C | H5 | 12 | U | H3' | 2.96 | 3.7 |
| 13 | C | H5 | 12 | U | H5 | 2.99 | 4.53 |
| 13 | C | H6 | 12 | U | H1' | 3.12 | 4.875 |
| 13 | C | H6 | 12 | U | H2' | 1.75 | 2.92 |
| 13 | C | H6 | 12 | U | H3' | 2.72 | 4.25 |
| 13 | C | H6 | 13 | C | H1' | 2.55 | 3.96 |
| 13 | C | H6 | 13 | C | H2' | 2.75 | 4.1 |

| | | | | | | | |
|---|---|---|---|---|---|---|---|
| 13 | C | H6 | 13 | C | H3' | 2 | 2.5 |
| 13 | C | N3 | 14 | G | H1 | 1.8 | 2.4 |
| 13 | C | O2 | 14 | G | H22 | 1.8 | 2.4 |
| 14 | G | H1 | 15 | A | H1' | 3.32 | 5.19 |
| 14 | G | H1 | 15 | A | H2 | 3.42 | 5.34 |
| 14 | G | H1' | 14 | G | H2' | 2.12 | 3.46 |
| 14 | G | H1' | 14 | G | H8 | 2.7 | 4.22 |
| 14 | G | H3' | 14 | G | H1' | 2.58 | 3.86 |
| 14 | G | H8 | 14 | G | H2' | 2.8 | 4.38 |
| 15 | A | H1' | 15 | A | H2' | 2.15 | 3.23 |
| 15 | A | H1' | 15 | A | H3' | 2.51 | 3.99 |
| 15 | A | H2 | 13 | C | H1' | 3.12 | 4.88 |
| 15 | A | H2 | 16 | C | H1' | 2.8 | 4.38 |
| 15 | A | H2 | 16 | C | H1' | 2.84 | 4.52 |
| 15 | A | H8 | 14 | G | H1' | 3.04 | 4.75 |
| 15 | A | H8 | 14 | G | H2' | 1.6 | 2.94 |
| 15 | A | H8 | 15 | A | H1' | 2.88 | 4.48 |
| 15 | A | H8 | 15 | A | H2' | 2.75 | 4.1 |
| 15 | A | H8 | 15 | A | H3' | 2 | 3.29 |
| 15 | A | H8 | 16 | C | H5 | 2.96 | 4.63 |
| 16 | C | H1' | 16 | C | H2' | 2.07 | 3.39 |
| 16 | C | H1' | 16 | C | H4' | 2.83 | 9.22 |
| 16 | C | H5 | 15 | A | H2' | 3.04 | 4.75 |
| 16 | C | H5 | 16 | C | H3' | 2.88 | 4.48 |
| 16 | C | H6 | 15 | A | H1' | 3.84 | 5.25 |
| 16 | C | H6 | 15 | A | H2' | 1.7 | 3 |
| 16 | C | H6 | 15 | A | H3' | 2.53 | 3.85 |
| 16 | C | H6 | 16 | C | H1' | 2.74 | 4.17 |
| 16 | C | H6 | 16 | C | H2' | 2.75 | 4.1 |
| 16 | C | H6 | 16 | C | H4' | 2.78 | 4.33 |
| 16 | C | H6 | 17 | A | H8 | 2.8 | 4.375 |
| 17 | A | H1' | 17 | A | H2' | 2.26 | 3.63 |
| 17 | A | H2 | 11 | G | H1' | 2.88 | 4.5 |
| 17 | A | H2 | 17 | A | H1' | 3.04 | 4.75 |
| 17 | A | H2 | 18 | G | H1' | 2.96 | 4.63 |
| 17 | A | H8 | 16 | C | H1' | 2.88 | 4.5 |
| 17 | A | H8 | 16 | C | H2' | 1.88 | 3.34 |
| 17 | A | H8 | 16 | C | H3' | 2.88 | 4.5 |
| 17 | A | H8 | 17 | A | H1' | 2.75 | 4.1 |
| 17 | A | H8 | 17 | A | H2' | 3.04 | 4.75 |
| 17 | A | H8 | 17 | A | H3' | 2.72 | 4.25 |
| 18 | G | H1' | 18 | G | H2' | 2.11 | 3.43 |
| 18 | G | H1' | 18 | G | H3' | 2.57 | 3.78 |
| 18 | G | H2' | 18 | G | H1' | 2.14 | 3.51 |
| 18 | G | H2' | 19 | C | H5 | 2.94 | 4.55 |
| 18 | G | H3' | 19 | C | H5 | 2.65 | 3.94 |
| 18 | G | H3' | 19 | C | H5 | 2.65 | 3.99 |
| 18 | G | H4' | 18 | G | H1' | 2.56 | 4.06 |
| 18 | G | H8 | 17 | A | H1' | 3.12 | 4.875 |
| 18 | G | H8 | 17 | A | H2' | 1.95 | 3.36 |
| 18 | G | H8 | 17 | A | H3' | 1.92 | 3 |

| | | | | | | | |
|---|---|---|---|---|---|---|---|
| 18 | G | H8 | 18 | G | H1' | 2.77 | 4.09 |
| 18 | G | H8 | 18 | G | H2' | 2.74 | 4.42 |
| 18 | G | H8 | 18 | G | H3' | 2.08 | 3.25 |
| 18 | G | H8 | 19 | C | H5 | 2.75 | 3.91 |
| 19 | C | H1' | 18 | G | H2' | 3.21 | 4.78 |
| 19 | C | H1' | 19 | C | H3' | 2.8 | 4.38 |
| 19 | C | H3' | 19 | C | H5 | 2.96 | 4.63 |
| 19 | C | H4' | 19 | C | H1' | 2.18 | 3.91 |
| 19 | C | H6 | 18 | G | H1' | 3.2 | 5 |
| 19 | C | H6 | 18 | G | H2' | 1.83 | 2.9 |
| 19 | C | H6 | 18 | G | H3' | 2 | 3.06 |
| 19 | C | H6 | 18 | G | H8 | 3.84 | 5.25 |
| 19 | C | H6 | 19 | C | H1' | 2.96 | 4.03 |
| 19 | C | H6 | 19 | C | H2' | 2.75 | 4.1 |
| 19 | C | H6 | 19 | C | H3' | 2 | 3.13 |
| 19 | C | H6 | 19 | C | H4' | 2.8 | 4.38 |
| 20 | U | H1' | 20 | U | H6 | 2.8 | 4.38 |
| 20 | U | H6 | 19 | C | H1' | 4.4 | 5.5 |
| 20 | U | H6 | 19 | C | H2' | 2.48 | 3.88 |
| 20 | U | H6 | 19 | C | H3' | 3 | 5 |
| 20 | U | H6 | 20 | U | H1' | 2.8 | 4.17 |
| 20 | U | H6 | 20 | U | H2' | 2.8 | 4.17 |
| 21 | G | H4' | 21 | G | H1' | 2.64 | 4.13 |
| 21 | G | H8 | 20 | U | H2' | 2.32 | 3.63 |
| 21 | G | H8 | 21 | G | H1' | 2.8 | 4.45 |
| 21 | G | H8 | 21 | G | H2' | 2.22 | 3.71 |
| 22 | C | H1' | 22 | C | H2' | 2.1 | 3.55 |
| 22 | C | H1' | 22 | C | H3' | 2.22 | 4.4 |
| 22 | C | H2' | 23 | U | H5 | 2.91 | 4.28 |
| 22 | C | H3' | 23 | U | H5 | 2.72 | 4.1 |
| 22 | C | H6 | 21 | G | H1' | 2.72 | 4.25 |
| 22 | C | H6 | 22 | C | H2' | 2.8 | 4.63 |
| 22 | C | H6 | 22 | C | H3' | 2.09 | 3.49 |
| 22 | C | H6 | 23 | U | H5 | 3.12 | 4.88 |
| 23 | U | H3 | 4 | A | H2 | 2.28 | 3.56 |
| 23 | U | H3 | 5 | G | H1 | 3.26 | 5.1 |
| 23 | U | H6 | 22 | C | H1' | 3.04 | 5 |
| 23 | U | H6 | 22 | C | H2' | 1.85 | 3.19 |
| 23 | U | H6 | 22 | C | H3' | 2.19 | 3.36 |
| 23 | U | H6 | 23 | U | H1' | 3.6 | 5.57 |
| 23 | U | H6 | 24 | G | H8 | 3.04 | 4.75 |
| 24 | G | H1 | 2 | A | H2 | 2.64 | 4.13 |
| 24 | G | H1' | 24 | G | H2' | 2.22 | 3.71 |
| 24 | G | H8 | 23 | U | H1' | 3.69 | 5.77 |
| 24 | G | H8 | 24 | G | H1' | 2.96 | 4.03 |
| 24 | G | H8 | 24 | G | H2' | 2.64 | 4.13 |
| 24 | G | H8 | 24 | G | H3' | 2 | 3.29 |
| 25 | U | H1' | 24 | G | H2' | 3.41 | 5.41 |
| 25 | U | H1' | 25 | U | H2' | 2.09 | 3.51 |
| 25 | U | H1' | 25 | U | H3' | 2.99 | 4.53 |
| 25 | U | H3 | 2 | A | H2 | 2.12 | 3.31 |

| | | | | | | | |
|---|---|---|---|---|---|---|---|
| 25 | U | H4' | 25 | U | H1' | 2.79 | 4.41 |
| 25 | U | H6 | 24 | G | H2' | 1.75 | 2.92 |
| 25 | U | H6 | 24 | G | H3' | 2 | 3.53 |
| 25 | U | H6 | 25 | U | H1' | 3.6 | 5.57 |
| 25 | U | H6 | 25 | U | H3' | 2 | 3.18 |
| 26 | C | H1' | 25 | U | H2' | 3.29 | 4.73 |
| 26 | C | H5 | 25 | U | H2' | 3.68 | 4.6 |
| 26 | C | H5 | 25 | U | H3' | 2.96 | 3.7 |
| 26 | C | H5 | 25 | U | H5 | 2.89 | 3.61 |
| 26 | C | H6 | 25 | U | H1' | 3.12 | 4.875 |
| 26 | C | H6 | 25 | U | H2' | 1.75 | 2.92 |
| 26 | C | H6 | 25 | U | H3' | 2.72 | 4.25 |
| 26 | C | H6 | 26 | C | H1' | 2.55 | 3.96 |
| 26 | C | H6 | 26 | C | H2' | 2.75 | 4.1 |
| 27 | **2** | H1 | 6 | C | H1' | 3 | 6 |
| 27 | **2** | H1 | 7 | U | H5 | 3 | 6 |
| 27 | **2** | H1 | 7 | U | H6 | 3 | 5 |
| 27 | **2** | H13 | 20 | U | H6 | 3 | 5 |
| 27 | **2** | H13 | 21 | G | H5'' | 3 | 5 |
| 27 | **2** | H17 | 27 | D6L | H8 | 3 | 5 |
| 27 | **2** | H19 | 27 | D6L | H8 | 2.5 | 5 |
| 27 | **2** | H26 | 7 | U | H3 | 3 | 5 |
| 27 | **2** | H26 | 7 | U | H3 | 2 | 4 |
| 27 | **2** | H3 | 6 | C | H1' | 3 | 6.5 |
| 27 | **2** | H4 | 7 | U | H3 | 2 | 4 |
| 27 | **2** | H4 | 7 | U | H5 | 3 | 6 |
| 27 | **2** | H5 | 19 | C | H2' | 3 | 5 |
| 27 | **2** | H5 | 19 | C | H5 | 3 | 5 |
| 27 | **2** | H5 | 19 | C | H5 | 3 | 5 |
| 27 | **2** | H5 | 19 | C | H6 | 3 | 5 |
| 27 | **2** | H5 | 20 | U | H5 | 3 | 5 |
| 27 | **2** | H5 | 20 | U | H6 | 3 | 5 |
| 27 | **2** | H6 | 20 | U | H3 | 3 | 5 |
| 27 | **2** | H6 | 20 | U | H3 | 2 | 4 |
| 27 | **2** | H7 | 19 | C | H3' | 3 | 5 |
| 27 | **2** | H7 | 19 | C | H3' | 3 | 5 |
| 27 | **2** | H7 | 19 | C | H5 | 3 | 5 |
| 27 | **2** | H7 | 20 | U | H2' | 4 | 6 |
| 27 | **2** | H7 | 20 | U | H5 | 2.5 | 5 |

**Table S7:** NOE restraints used for modeling of the r(CUG)-**3** complex.

| | | | | | | | |
|---|---|---|---|---|---|---|---|
| 1 | G | H1 | 26 | C | N3 | 1.8 | 2.4 |
| 1 | G | H1' | 1 | G | H2' | 2.27 | 3.55 |
| 1 | G | H1' | 1 | G | H3' | 3.56 | 5.56 |
| 1 | G | H1' | 1 | G | H4' | 2.62 | 4.09 |
| 1 | G | H22 | 26 | C | O2 | 1.8 | 2.4 |
| 1 | G | H8 | 1 | G | H1' | 2.66 | 4.16 |
| 1 | G | H8 | 1 | G | H2' | 2.34 | 3.8 |
| 1 | G | H8 | 1 | G | H3' | 2.16 | 3.51 |
| 1 | G | H8 | 1 | G | H5' | 2.42 | 3.79 |
| 1 | G | H8 | 2 | A | H8 | 3.36 | 5.25 |
| 1 | G | O6 | 26 | C | H41 | 1.8 | 2.4 |
| 2 | A | H1' | 2 | A | H2 | 3.12 | 5.07 |
| 2 | A | H1' | 2 | A | H3' | 2.65 | 4.14 |
| 2 | A | H1' | 2 | A | H4' | 2.16 | 3.51 |
| 2 | A | H2 | 2 | A | H1' | 2.96 | 4.81 |
| 2 | A | H2 | 3 | C | H1' | 2.56 | 4.16 |
| 2 | A | H2 | 26 | C | H1' | 2.96 | 4.81 |
| 2 | A | H61 | 25 | U | O4 | 1.8 | 2.4 |
| 2 | A | H8 | 1 | G | H1' | 3.23 | 5.05 |
| 2 | A | H8 | 1 | G | H2' | 2.34 | 3.8 |
| 2 | A | H8 | 1 | G | H3' | 2.34 | 3.8 |
| 2 | A | H8 | 1 | G | H8 | 3.23 | 5.05 |
| 2 | A | H8 | 2 | A | H1' | 2.63 | 4.11 |
| 2 | A | H8 | 2 | A | H2' | 2.34 | 3.8 |
| 2 | A | H8 | 2 | A | H3' | 2.1 | 3.28 |
| 2 | A | H8 | 3 | C | H6 | 3.23 | 5.05 |
| 2 | A | N1 | 25 | U | H3 | 1.8 | 2.4 |
| 3 | C | H1' | 2 | A | H2' | 2.64 | 4.29 |
| 3 | C | H1' | 3 | C | H2' | 2.14 | 3.34 |
| 3 | C | H1' | 3 | C | H3' | 2.32 | 3.77 |
| 3 | C | H1' | 3 | C | H4' | 2.81 | 4.4 |
| 3 | C | H41 | 24 | G | O6 | 1.8 | 2.4 |
| 3 | C | H5 | 2 | A | H2' | 3.23 | 5.05 |
| 3 | C | H5 | 2 | A | H3' | 2.57 | 4.02 |
| 3 | C | H5 | 3 | C | H3' | 2.81 | 4.4 |
| 3 | C | H6 | 2 | A | H1' | 3.23 | 5.05 |
| 3 | C | H6 | 2 | A | H2' | 1.89 | 3.32 |
| 3 | C | H6 | 2 | A | H3' | 2.47 | 3.85 |
| 3 | C | H6 | 2 | A | H8 | 3.23 | 5.05 |
| 3 | C | H6 | 3 | C | H1' | 2.57 | 4.02 |
| 3 | C | H6 | 3 | C | H3' | 1.88 | 2.94 |
| 3 | C | H6 | 3 | C | H5'' | 3.23 | 5.05 |
| 3 | C | H6 | 4 | A | H8 | 3.23 | 5.05 |
| 3 | C | N3 | 24 | G | H1 | 1.8 | 2.4 |
| 3 | C | O2 | 24 | G | H22 | 1.8 | 2.4 |
| 4 | A | H1' | 3 | C | H2' | 2.48 | 4.03 |
| 4 | A | H1' | 4 | A | H4' | 2.93 | 4.57 |
| 4 | A | H1' | 5 | G | H8 | 2.88 | 4.68 |
| 4 | A | H2 | 4 | A | H1' | 2.96 | 4.81 |

| | | | | | | | |
|---|---|---|---|---|---|---|---|
| 4 | A | H2 | 5 | G | H1' | 2.72 | 4.42 |
| 4 | A | H2 | 24 | G | H1' | 2.74 | 4.28 |
| 4 | A | H61 | 23 | U | O4 | 1.8 | 2.4 |
| 4 | A | H8 | 3 | C | H1' | 3.12 | 5.07 |
| 4 | A | H8 | 3 | C | H2' | 1.61 | 2.52 |
| 4 | A | H8 | 3 | C | H3' | 2.16 | 3.38 |
| 4 | A | H8 | 3 | C | H5 | 3.81 | 5.95 |
| 4 | A | H8 | 3 | C | H6 | 3.23 | 5.05 |
| 4 | A | H8 | 4 | A | H1' | 2.74 | 4.29 |
| 4 | A | H8 | 4 | A | H2' | 2.57 | 4.02 |
| 4 | A | H8 | 4 | A | H3' | 2.57 | 4.02 |
| 4 | A | N1 | 23 | U | H3 | 1.8 | 2.4 |
| 5 | G | H1 | 22 | C | N3 | 1.8 | 2.4 |
| 5 | G | H1' | 4 | A | H2' | 4.09 | 6.38 |
| 5 | G | H1' | 5 | G | H2' | 2.11 | 3.29 |
| 5 | G | H1' | 5 | G | H3' | 3.56 | 5.56 |
| 5 | G | H1' | 5 | G | H4' | 2.95 | 4.61 |
| 5 | G | H22 | 22 | C | O2 | 1.8 | 2.4 |
| 5 | G | H8 | 4 | A | H1' | 2.93 | 4.58 |
| 5 | G | H8 | 4 | A | H2' | 2.07 | 3.36 |
| 5 | G | H8 | 4 | A | H3' | 3.17 | 4.96 |
| 5 | G | H8 | 4 | A | H8 | 4.28 | 6.95 |
| 5 | G | H8 | 5 | G | H1' | 2.95 | 4.62 |
| 5 | G | H8 | 5 | G | H2' | 3.04 | 4.75 |
| 5 | G | H8 | 5 | G | H3' | 2.33 | 3.64 |
| 5 | G | O6 | 22 | C | H41 | 1.8 | 2.4 |
| 6 | C | H1' | 5 | G | H2' | 2.72 | 4.42 |
| 6 | C | H1' | 6 | C | H2' | 2.06 | 3.22 |
| 6 | C | H1' | 6 | C | H3' | 2.5 | 4.38 |
| 6 | C | H1' | 6 | C | H4' | 2.5 | 4.38 |
| 6 | C | H3' | 7 | U | H5 | 2.5 | 4.38 |
| 6 | C | H41 | 21 | G | O6 | 1.8 | 2.4 |
| 6 | C | H5 | 5 | G | H2' | 2.9 | 4.53 |
| 6 | C | H5 | 5 | G | H8 | 2.75 | 4.3 |
| 6 | C | H6 | 5 | G | H1' | 3.12 | 5.07 |
| 6 | C | H6 | 5 | G | H2' | 1.77 | 3.09 |
| 6 | C | H6 | 5 | G | H3' | 2.27 | 3.54 |
| 6 | C | H6 | 5 | G | H8 | 3.48 | 6.09 |
| 6 | C | H6 | 6 | C | H1' | 2.64 | 4.12 |
| 6 | C | H6 | 6 | C | H2' | 2.3 | 3.74 |
| 6 | C | H6 | 6 | C | H3' | 2.5 | 4.38 |
| 6 | C | H6 | 6 | C | H5 | 2.01 | 3.15 |
| 6 | C | H6 | 6 | C | H5'' | 2.73 | 4.27 |
| 6 | C | N3 | 21 | G | H1 | 1.8 | 2.4 |
| 6 | C | O2 | 21 | G | H22 | 1.8 | 2.4 |
| 7 | U | H1' | 6 | C | H2' | 2.67 | 4.17 |
| 7 | U | H1' | 7 | U | H2' | 2.3 | 4.03 |
| 7 | U | H1' | 7 | U | H3' | 3.36 | 5.25 |
| 7 | U | H1' | 7 | U | H4' | 2.92 | 4.56 |
| 7 | U | H6 | 6 | C | H1' | 3.36 | 5.25 |
| 7 | U | H6 | 6 | C | H2' | 2.01 | 3.15 |

| | | | | | | | |
|---|---|---|---|---|---|---|---|
| 7 | U | H6 | 7 | U | H1' | 2.59 | 4.21 |
| 7 | U | H6 | 7 | U | H2' | 2.64 | 4.63 |
| 7 | U | H6 | 7 | U | H3' | 3.36 | 5.25 |
| 7 | U | H6 | 7 | U | H4' | 2.5 | 4.38 |
| 7 | U | H6 | 7 | U | H5'' | 2.44 | 3.81 |
| 8 | G | H1 | 19 | C | N3 | 1.8 | 2.4 |
| 8 | G | H22 | 19 | C | O2 | 1.8 | 2.4 |
| 8 | G | H8 | 7 | U | H1' | 3.2 | 5 |
| 8 | G | H8 | 7 | U | H2' | 2.05 | 3.2 |
| 8 | G | H8 | 7 | U | H3' | 3.36 | 5.25 |
| 8 | G | H8 | 8 | G | H1' | 2.59 | 4.04 |
| 8 | G | H8 | 8 | G | H2' | 2.98 | 4.65 |
| 8 | G | O6 | 19 | C | H41 | 1.8 | 2.4 |
| 9 | C | H1' | 8 | G | H2' | 3.89 | 6.08 |
| 9 | C | H1' | 9 | C | H2' | 2.25 | 3.51 |
| 9 | C | H1' | 9 | C | H3' | 2.98 | 4.65 |
| 9 | C | H41 | 18 | G | O6 | 1.8 | 2.4 |
| 9 | C | H5 | 8 | G | H2' | 2.61 | 4.07 |
| 9 | C | H5 | 9 | C | H3' | 2.8 | 4.37 |
| 9 | C | H6 | 8 | G | H1' | 2.98 | 5 |
| 9 | C | H6 | 8 | G | H2' | 1.95 | 3.16 |
| 9 | C | H6 | 9 | C | H1' | 2.56 | 4.01 |
| 9 | C | H6 | 9 | C | H2' | 2.98 | 4.65 |
| 9 | C | H6 | 9 | C | H3' | 2.1 | 3.68 |
| 9 | C | H6 | 9 | C | H5 | 2.05 | 3.21 |
| 9 | C | N3 | 18 | G | H1 | 1.8 | 2.4 |
| 9 | C | O2 | 18 | G | H22 | 1.8 | 2.4 |
| 10 | U | H3 | 17 | A | N1 | 1.8 | 2.4 |
| 10 | U | H5 | 9 | C | H2' | 2.61 | 4.07 |
| 10 | U | H5 | 9 | C | H3' | 2.68 | 4.18 |
| 10 | U | H5 | 10 | U | H2' | 4.65 | 7.26 |
| 10 | U | H6 | 9 | C | H1' | 3.12 | 5.07 |
| 10 | U | H6 | 9 | C | H2' | 1.97 | 3.08 |
| 10 | U | H6 | 9 | C | H3' | 2.45 | 3.82 |
| 10 | U | H6 | 9 | C | H5 | 3.81 | 5.95 |
| 10 | U | H6 | 10 | U | H1' | 2.6 | 4.07 |
| 10 | U | H6 | 10 | U | H2' | 2.39 | 3.73 |
| 10 | U | H6 | 10 | U | H5 | 2.06 | 3.22 |
| 10 | U | O4 | 17 | A | H61 | 1.8 | 2.4 |
| 11 | G | H1 | 16 | C | N3 | 1.8 | 2.4 |
| 11 | G | H1' | 11 | G | H2' | 2.27 | 3.55 |
| 11 | G | H1' | 11 | G | H3' | 3.38 | 5.28 |
| 11 | G | H22 | 16 | C | O2 | 1.8 | 2.4 |
| 11 | G | H8 | 10 | U | H1' | 2.79 | 4.88 |
| 11 | G | H8 | 10 | U | H2' | 1.97 | 3.08 |
| 11 | G | H8 | 10 | U | H5 | 3.01 | 4.71 |
| 11 | G | H8 | 11 | G | H1' | 3.16 | 4.93 |
| 11 | G | H8 | 11 | G | H3' | 2.51 | 3.93 |
| 11 | G | O6 | 16 | C | H41 | 1.8 | 2.4 |
| 12 | U | H1' | 12 | U | H2' | 2.07 | 3.23 |
| 12 | U | H1' | 12 | U | H3' | 2.72 | 4.25 |

| | | | | | | | |
|---|---|---|---|---|---|---|---|
| 12 | U | H1' | 12 | U | H4' | 2.72 | 4.25 |
| 12 | U | H3 | 15 | A | N1 | 1.8 | 2.4 |
| 12 | U | H3' | 12 | U | H5 | 3.12 | 5.07 |
| 12 | U | H5 | 11 | G | H2' | 3.65 | 5.93 |
| 12 | U | H5 | 11 | G | H3' | 3.65 | 5.93 |
| 12 | U | H6 | 11 | G | H1' | 3.03 | 4.73 |
| 12 | U | H6 | 11 | G | H2' | 1.87 | 2.92 |
| 12 | U | H6 | 11 | G | H3' | 2.39 | 3.73 |
| 12 | U | H6 | 12 | U | H1' | 2.7 | 4.72 |
| 12 | U | H6 | 12 | U | H2' | 2.8 | 4.37 |
| 12 | U | H6 | 12 | U | H3' | 1.93 | 3.39 |
| 12 | U | H6 | 12 | U | H5 | 2 | 3.12 |
| 12 | U | H6 | 12 | U | H5'' | 2.71 | 4.24 |
| 12 | U | O4 | 15 | A | H61 | 1.8 | 2.4 |
| 13 | C | H1' | 12 | U | H2' | 2.96 | 4.81 |
| 13 | C | H1' | 13 | C | H2' | 1.99 | 3.11 |
| 13 | C | H3' | 13 | C | H2' | 1.79 | 2.8 |
| 13 | C | H41 | 14 | G | O6 | 1.8 | 2.4 |
| 13 | C | H5 | 12 | U | H2' | 2.85 | 4.46 |
| 13 | C | H5 | 12 | U | H3' | 3.31 | 5.17 |
| 13 | C | H5 | 12 | U | H6 | 2.67 | 4.67 |
| 13 | C | H5 | 13 | C | H2' | 3.26 | 5.1 |
| 13 | C | H5 | 13 | C | H3' | 3.04 | 4.76 |
| 13 | C | H6 | 12 | U | H1' | 3.43 | 5.36 |
| 13 | C | H6 | 12 | U | H2' | 1.83 | 2.86 |
| 13 | C | H6 | 12 | U | H3' | 2.15 | 3.77 |
| 13 | C | H6 | 13 | C | H1' | 2.48 | 3.87 |
| 13 | C | H6 | 13 | C | H2' | 2.26 | 3.67 |
| 13 | C | H6 | 13 | C | H3' | 1.89 | 2.95 |
| 13 | C | H6 | 13 | C | H5'' | 2.79 | 4.35 |
| 13 | C | N3 | 14 | G | H1 | 1.8 | 2.4 |
| 13 | C | O2 | 14 | G | H22 | 1.8 | 2.4 |
| 14 | G | H1' | 14 | G | H2' | 2.27 | 3.55 |
| 14 | G | H1' | 14 | G | H3' | 3.56 | 5.56 |
| 14 | G | H1' | 14 | G | H4' | 2.62 | 4.09 |
| 14 | G | H4' | 14 | G | H5'' | 2 | 3.13 |
| 14 | G | H8 | 14 | G | H1' | 2.66 | 4.16 |
| 14 | G | H8 | 14 | G | H2' | 2.34 | 3.8 |
| 14 | G | H8 | 14 | G | H3' | 2.16 | 3.51 |
| 14 | G | H8 | 15 | A | H8 | 3.36 | 5.25 |
| 15 | A | H1' | 15 | A | H2 | 3.12 | 5.07 |
| 15 | A | H1' | 15 | A | H3' | 2.65 | 4.14 |
| 15 | A | H2 | 13 | C | H1' | 2.64 | 4.29 |
| 15 | A | H2 | 15 | A | H1' | 2.96 | 4.81 |
| 15 | A | H2 | 16 | C | H1' | 2.57 | 4.01 |
| 15 | A | H8 | 14 | G | H1' | 3.23 | 5.05 |
| 15 | A | H8 | 14 | G | H2' | 2.34 | 3.8 |
| 15 | A | H8 | 14 | G | H3' | 2.34 | 3.8 |
| 15 | A | H8 | 14 | G | H8 | 3.23 | 5.05 |
| 15 | A | H8 | 15 | A | H1' | 2.63 | 4.11 |
| 15 | A | H8 | 15 | A | H2' | 2.34 | 3.8 |

| | | | | | | | |
|---|---|---|---|---|---|---|---|
| 15 | A | H8 | 15 | A | H3' | 2.1 | 3.28 |
| 15 | A | H8 | 16 | C | H6 | 3.23 | 5.05 |
| 16 | C | H1' | 15 | A | H2' | 2.64 | 4.29 |
| 16 | C | H1' | 16 | C | H2' | 2.14 | 3.34 |
| 16 | C | H1' | 16 | C | H3' | 2.32 | 3.77 |
| 16 | C | H1' | 16 | C | H4' | 2.81 | 4.4 |
| 16 | C | H5 | 15 | A | H2' | 3.23 | 5.05 |
| 16 | C | H5 | 15 | A | H3' | 2.57 | 4.02 |
| 16 | C | H5 | 16 | C | H3' | 2.73 | 4.27 |
| 16 | C | H6 | 15 | A | H1' | 3.23 | 5.05 |
| 16 | C | H6 | 15 | A | H2' | 1.89 | 3.32 |
| 16 | C | H6 | 15 | A | H3' | 2.47 | 3.85 |
| 16 | C | H6 | 15 | A | H8 | 3.23 | 5.05 |
| 16 | C | H6 | 16 | C | H1' | 2.57 | 4.02 |
| 16 | C | H6 | 16 | C | H3' | 1.88 | 2.94 |
| 16 | C | H6 | 16 | C | H5'' | 3.23 | 5.05 |
| 16 | C | H6 | 17 | A | H8 | 3.23 | 5.05 |
| 17 | A | H1' | 16 | C | H2' | 2.93 | 4.58 |
| 17 | A | H1' | 18 | G | H8 | 2.93 | 4.58 |
| 17 | A | H2 | 10 | U | H1' | 3 | 6 |
| 17 | A | H2 | 11 | G | H1' | 2.74 | 4.28 |
| 17 | A | H2 | 17 | A | H1' | 2.96 | 4.81 |
| 17 | A | H2 | 18 | G | H1' | 2.9 | 4.53 |
| 17 | A | H8 | 16 | C | H1' | 3.12 | 5.07 |
| 17 | A | H8 | 16 | C | H2' | 1.61 | 2.52 |
| 17 | A | H8 | 16 | C | H3' | 2.16 | 3.38 |
| 17 | A | H8 | 16 | C | H5 | 3.81 | 5.95 |
| 17 | A | H8 | 16 | C | H6 | 3.23 | 5.05 |
| 17 | A | H8 | 17 | A | H1' | 2.74 | 4.29 |
| 17 | A | H8 | 17 | A | H2' | 2.57 | 4.02 |
| 17 | A | H8 | 17 | A | H3' | 2.57 | 4.02 |
| 18 | G | H1' | 17 | A | H2' | 4.09 | 6.38 |
| 18 | G | H1' | 18 | G | H2' | 2.11 | 3.29 |
| 18 | G | H1' | 18 | G | H3' | 3.56 | 5.56 |
| 18 | G | H1' | 18 | G | H4' | 2.95 | 4.61 |
| 18 | G | H8 | 17 | A | H1' | 2.93 | 4.58 |
| 18 | G | H8 | 17 | A | H2' | 2.07 | 3.36 |
| 18 | G | H8 | 17 | A | H3' | 3.17 | 4.96 |
| 18 | G | H8 | 17 | A | H8 | 4.4 | 6.87 |
| 18 | G | H8 | 18 | G | H1' | 2.95 | 4.62 |
| 18 | G | H8 | 18 | G | H2' | 3.04 | 4.75 |
| 18 | G | H8 | 18 | G | H3' | 2.33 | 3.64 |
| 18 | G | H8 | 18 | G | H5'' | 2.76 | 4.49 |
| 19 | C | H1' | 18 | G | H2' | 2.44 | 4.27 |
| 19 | C | H1' | 19 | C | H2' | 2.06 | 3.22 |
| 19 | C | H1' | 19 | C | H3' | 2.5 | 4.38 |
| 19 | C | H1' | 19 | C | H4' | 2.5 | 4.38 |
| 19 | C | H5 | 18 | G | H2' | 2.9 | 4.53 |
| 19 | C | H5 | 18 | G | H8 | 2.75 | 4.3 |
| 19 | C | H6 | 18 | G | H1' | 3.12 | 5.07 |
| 19 | C | H6 | 18 | G | H2' | 1.77 | 3.09 |

| | | | | | | | |
|---|---|---|---|---|---|---|---|
| 19 | C | H6 | 18 | G | H3' | 2.27 | 3.54 |
| 19 | C | H6 | 18 | G | H8 | 3.48 | 6.09 |
| 19 | C | H6 | 19 | C | H1' | 2.64 | 4.12 |
| 19 | C | H6 | 19 | C | H2' | 2.42 | 3.78 |
| 19 | C | H6 | 19 | C | H3' | 2.42 | 3.78 |
| 19 | C | H6 | 19 | C | H5'' | 2.73 | 4.27 |
| 20 | U | H1' | 19 | C | H2' | 2.5 | 4.38 |
| 20 | U | H1' | 20 | U | H2' | 2.3 | 4.03 |
| 20 | U | H1' | 20 | U | H3' | 3.36 | 5.25 |
| 20 | U | H6 | 19 | C | H1' | 3.36 | 5.25 |
| 20 | U | H6 | 19 | C | H2' | 2.01 | 3.15 |
| 20 | U | H6 | 20 | U | H1' | 2.71 | 4.24 |
| 20 | U | H6 | 20 | U | H2' | 2.84 | 4.62 |
| 20 | U | H6 | 20 | U | H3' | 2.5 | 4.38 |
| 20 | U | H6 | 20 | U | H4' | 2.5 | 4.38 |
| 21 | G | H8 | 20 | U | H1' | 3.36 | 6 |
| 21 | G | H8 | 20 | U | H2' | 2.05 | 4.5 |
| 21 | G | H8 | 20 | U | H3' | 1.93 | 3.39 |
| 21 | G | H8 | 21 | G | H1' | 3.12 | 5.07 |
| 21 | G | H8 | 21 | G | H2' | 2.98 | 4.65 |
| 21 | G | H8 | 21 | G | H5'' | 2.5 | 5 |
| 22 | C | H1' | 21 | G | H2' | 3.89 | 6.08 |
| 22 | C | H1' | 22 | C | H2' | 2.25 | 3.51 |
| 22 | C | H1' | 22 | C | H3' | 2.4 | 3.9 |
| 22 | C | H5 | 21 | G | H2' | 2.61 | 4.07 |
| 22 | C | H5 | 22 | C | H3' | 2.8 | 4.37 |
| 22 | C | H6 | 21 | G | H1' | 2.99 | 5 |
| 22 | C | H6 | 21 | G | H2' | 1.95 | 3.16 |
| 22 | C | H6 | 22 | C | H1' | 2.56 | 4.01 |
| 22 | C | H6 | 22 | C | H2' | 2.98 | 4.65 |
| 22 | C | H6 | 22 | C | H3' | 2.1 | 3.68 |
| 22 | C | H6 | 22 | C | H5 | 2.05 | 3.21 |
| 23 | U | H5 | 22 | C | H2' | 2.61 | 4.07 |
| 23 | U | H5 | 22 | C | H3' | 2.68 | 4.18 |
| 23 | U | H5 | 23 | U | H2' | 4.65 | 7.26 |
| 23 | U | H6 | 22 | C | H1' | 3.12 | 5.07 |
| 23 | U | H6 | 22 | C | H2' | 1.97 | 3.08 |
| 23 | U | H6 | 22 | C | H3' | 2.45 | 3.82 |
| 23 | U | H6 | 22 | C | H5 | 3.81 | 5.95 |
| 23 | U | H6 | 23 | U | H1' | 2.6 | 4.07 |
| 23 | U | H6 | 23 | U | H2' | 2.39 | 3.73 |
| 23 | U | H6 | 23 | U | H5 | 2.06 | 3.22 |
| 24 | G | H1' | 24 | G | H2' | 2.27 | 3.55 |
| 24 | G | H1' | 24 | G | H3' | 3.38 | 5.28 |
| 24 | G | H8 | 23 | U | H1' | 2.79 | 4.88 |
| 24 | G | H8 | 23 | U | H2' | 1.97 | 3.08 |
| 24 | G | H8 | 23 | U | H5 | 3.01 | 4.71 |
| 24 | G | H8 | 24 | G | H1' | 3.16 | 4.93 |
| 24 | G | H8 | 24 | G | H3' | 2.51 | 3.93 |
| 25 | U | H1' | 25 | U | H3' | 2.72 | 4.25 |
| 25 | U | H1' | 25 | U | H4' | 2.72 | 4.25 |

| | | | | | | | |
|---|---|---|---|---|---|---|---|
| 25 | U | H3' | 25 | U | H5 | 3.12 | 5.07 |
| 25 | U | H5 | 24 | G | H2' | 3.65 | 5.93 |
| 25 | U | H5 | 24 | G | H3' | 3.65 | 5.93 |
| 25 | U | H6 | 24 | G | H1' | 3.03 | 4.73 |
| 25 | U | H6 | 24 | G | H2' | 1.87 | 2.92 |
| 25 | U | H6 | 24 | G | H3' | 2.39 | 3.73 |
| 25 | U | H6 | 25 | U | H1' | 2.7 | 4.72 |
| 25 | U | H6 | 25 | U | H2' | 2.8 | 4.37 |
| 25 | U | H6 | 25 | U | H3' | 1.93 | 3.39 |
| 25 | U | H6 | 25 | U | H5 | 2 | 3.12 |
| 25 | U | H6 | 25 | U | H5'' | 2.71 | 4.24 |
| 26 | C | H1' | 25 | U | H2' | 2.9 | 4.53 |
| 26 | C | H1' | 26 | C | H2' | 1.99 | 3.11 |
| 26 | C | H1' | 26 | C | H3' | 2.4 | 3.9 |
| 26 | C | H3' | 26 | C | H2' | 1.79 | 2.8 |
| 26 | C | H5 | 25 | U | H2' | 2.85 | 4.46 |
| 26 | C | H5 | 25 | U | H3' | 3.31 | 5.17 |
| 26 | C | H5 | 25 | U | H6 | 2.67 | 4.67 |
| 26 | C | H5 | 26 | C | H2' | 3.26 | 5.1 |
| 26 | C | H5 | 26 | C | H3' | 3.04 | 4.76 |
| 26 | C | H6 | 25 | U | H1' | 3.43 | 5.36 |
| 26 | C | H6 | 25 | U | H2' | 1.83 | 2.86 |
| 26 | C | H6 | 25 | U | H3' | 2.15 | 3.77 |
| 26 | C | H6 | 26 | C | H1' | 2.48 | 3.87 |
| 26 | C | H6 | 26 | C | H2' | 2.38 | 3.72 |
| 26 | C | H6 | 26 | C | H3' | 1.89 | 2.95 |
| 26 | C | H6 | 26 | C | H5'' | 2.79 | 4.35 |
| 27 | **3** | C15 | 19 | C | H5 | 2.81 | 6 |
| 27 | **3** | C15 | 19 | C | H6 | 3 | 7 |
| 27 | **3** | C15 | 20 | U | H1' | 3 | 7 |
| 27 | **3** | C15 | 20 | U | H6 | 3 | 7 |
| 27 | **3** | C15 | 21 | G | H1' | 3 | 7 |
| 27 | **3** | C7 | 6 | C | H1' | 3 | 7 |
| 27 | **3** | C7 | 6 | C | H2' | 3 | 7 |
| 27 | **3** | C7 | 7 | U | H1' | 3 | 7 |
| 27 | **3** | C7 | 7 | U | H4' | 3 | 7 |
| 27 | **3** | C7 | 7 | U | H6 | 3 | 7 |
| 27 | **3** | H2 | 6 | C | H1' | 3 | 7 |
| 27 | **3** | H2 | 6 | C | H2' | 3 | 7 |
| 27 | **3** | H2 | 6 | C | H5 | 3 | 7 |
| 27 | **3** | H6 | 19 | C | H1' | 3 | 7 |
| 27 | **3** | H6 | 19 | C | H5 | 3 | 7 |
| 27 | **3** | H6 | 20 | U | H1' | 3 | 7 |
| 27 | **3** | H6 | 20 | U | H2' | 3 | 7 |
| 27 | **3** | H6 | 21 | G | H2' | 3 | 7 |
| 27 | **3** | H7 | 19 | C | H1' | 3 | 7 |
| 27 | **3** | H7 | 19 | C | H2' | 3 | 7 |
| 27 | **3** | H7 | 20 | U | H2' | 3 | 7 |
| 27 | **3** | H7 | 20 | U | H6 | 3 | 7 |
| 27 | **3** | H7 | 21 | G | H2' | 3 | 7 |
| 27 | **3** | H8 | 19 | C | H1' | 3 | 7 |

| | | | | | | | |
|---|---|---|---|---|---|---|---|
| 27 | **3** | H8 | 19 | C | H2' | 3 | 7 |
| 27 | **3** | H8 | 20 | U | H1' | 3 | 7 |
| 27 | **3** | H8 | 20 | U | H2' | 3 | 7 |

**Table S8:** $^{1}$H NMR chemical shifts of the unbound r(CUG) duplex.

| Residue | H1′ | H2′ | H3′ | H4′ | H2/H5 | H6/H8 | H1/H3 | H21/41/61, H22/42/62 amino |
|---|---|---|---|---|---|---|---|---|
| G1 | 5.709 | 4.831 | 4.654 | - | n/a | 8.052 | 12.59 | - |
| A2 | 6.073 | 4.518 | - | 4.636 | 7.832 | 8.188 | n/a | - |
| C3 | 5.424 | 4.321 | 4.515 | 4.413 | 5.209 | 7.605 | n/a | 8.272, 6.953 |
| A4 | 5.882 | 4.555 | 4.716 | 4.480 | 7.026 | 7.973 | n/a | - |
| G5 | 5.613 | 4.348 | 4.457 | - | n/a | 7.298 | 13.47 | - |
| C6 | 5.461 | 4.293 | - | - | 5.143 | 7.475 | n/a | 8.341, 6.649 |
| U7 | 5.475 | 4.294 | - | - | 5.462 | 7.606 | 10.3 | n/a |
| G8 | 5.770 | 4.532 | 4.636 | - | n/a | 7.826 | 13.24 | - |
| C9 | 5.513 | 4.352 | 4.435 | - | 5.231 | 7.761 | n/a | 8.586, 6.847 |
| U10 | 5.517 | 4.648 | 4.604 | - | 5.438 | 7.936 | 13.57 | n/a |
| G11 | 5.771 | 4.418 | 4.564 | 4.479 | n/a | 7.737 | 12.61 | - |
| U12 | 5.479 | 4.226 | 4.464 | - | 5.105 | 7.800 | 14.62 | n/a |
| C13 | 5.830 | 3.968 | 4.181 | - | 5.635 | 7.740 | n/a | 8.424, 7.035 |

Chemical shifts are reported in ppm units. Non-exchangeable protons were assigned at 25 °C. Exchangeable protons were assigned at 5 °C.

**Table S9:** $^1$H NMR chemical shifts of r(CUG)-**1** complex.

$^1$H NMR chemical shifts of RNA in the r(CUG)-**1** complex:

| Residue | H1′ | H2′ | H3′ | H4′ | H2/H5 | H6/H8 | H1/H3 | H21/41/61, H22/42/62 amino |
|---|---|---|---|---|---|---|---|---|
| G1 | 5.667 | 4.809 | - | 4.389 | n/a | 8.027 | 12.61 | - |
| A2 | 6.060 | 4.524 | 4.751 | - | 7.819 | 8.149 | n/a | - |
| C3 | 5.424 | 4.328 | 4.499 | - | 5.206 | 7.581 | n/a | 8.307, 6.975 |
| A4 | 5.890 | 4.556 | 4.709 | 4.485 | 7.050 | 7.960 | n/a | - |
| G5 | 5.608 | 4.337 | - | - | n/a | 7.266 | 13.50 | - |
| C6 | 5.492 | 4.273 | 4.308 | - | 5.131 | 7.442 | n/a | 8.372, 6.686 |
| U7 | 5.505 | 4.256 | - | - | 5.442 | 7.590 | - | n/a |
| G8 | 5.711 | 4.556 | 4.606 | - | n/a | 7.799 | 13.29 | - |
| C9 | 5.529 | 4.371 | 4.441 | - | 5.226 | 7.718 | n/a | 8.610, 6.873 |
| U10 | 5.534 | - | - | - | 5.412 | 7.921 | 13.61 | n/a |
| G11 | 5.779 | 4.427 | 4.563 | - | n/a | 7.751 | 12.64 | - |
| U12 | 5.486 | 4.211 | 4.462 | - | 5.106 | 7.781 | 14.64 | n/a |
| C13 | 5.826 | 3.948 | 4.146 | - | 5.603 | 7.707 | n/a | 8.455, 7.085 |

$^1$H NMR chemical shifts of **1** in the r(CUG)-**1** complex:

| Hydrogen | Chemical shift |
|---|---|
| H1/H2 | 7.236 |
| H3/H4 | 7.318 |
| H5/H6 | 8.010 |
| H7/H8 | 7.297 |

Chemical shifts are in ppm units. Non-exchangeable protons were assigned at 35 °C. Exchangeable protons were assigned at 6 °C.

**Table S10:** $^{1}$H NMR chemical shifts of r(CUG)-**2** complex.

$^{1}$H NMR chemical shifts of RNA in the r(CUG)-**2** complex:

| Residue | H1′ | H2′ | H3′ | H2/H5 | H6/H8 | H1/H3 | H21/41/61, H22/42/62 amino |
|---|---|---|---|---|---|---|---|
| G1 | 5.631 | 4.753 | - | n/a | 8.021 | - | - |
| A2 | 6.021 | 4.494 | 4.739 | 7.770 | 8.146 | n/a | - |
| C3 | 5.384 | 4.298 | 4.490 | 5.179 | 7.552 | n/a | 8.194, 6.873 |
| A4 | 5.854 | 4.542 | 4.674 | 6.996 | 7.928 | n/a | - |
| G5 | 5.565 | 4.244 | 4.394 | n/a | 7.217 | 13.37 | - |
| C6 | 5.525 | 4.199 | 4.344 | 4.986 | 7.358 | n/a | - |
| U7 | 5.534 | 4.181 | - | 5.531 | 7.715 | 10.41 | n/a |
| G8 | 5.658 | 4.324 | - | n/a | 7.725 | - | - |
| C9 | 5.498 | 4.333 | 4.415 | 5.175 | 7.707 | n/a | 8.510, 6.725 |
| U10 | 5.503 | - | - | 5.337 | 7.886 | 13.52 | n/a |
| G11 | 5.738 | 4.388 | 4.537 | n/a | 7.715 | 12.50 | - |
| U12 | 5.464 | 4.179 | 4.425 | 5.023 | 7.741 | 14.44 | n/a |
| C13 | 5.785 | 3.955 | 4.140 | 5.541 | 7.658 | n/a | - |

$^{1}$H NMR chemical shifts of **2** in the r(CUG)-**2** complex:

| Hydrogen | Chemical shift |
|---|---|
| H1/H2 | 7.555 |
| H3/H4 | 7.641 |
| H5/H6 | 7.997 |
| H7/H8 | 7.725 |

Chemical shifts are in ppm units. Non-exchangeable protons were assigned at 35 °C. Exchangeable protons were assigned at 6 °C.

**Table S11:** $^{1}$H NMR chemical shifts of r(CUG)-**3** complex.

$^{1}$H NMR chemical shifts of RNA in the r(CUG)-**3** complex:

| Residue | H1′ | H2′ | H3′ | H4′ | H2/H5 | H6/H8 | H1/H3 | H21/41/61, H22/42/62 amino |
|---|---|---|---|---|---|---|---|---|
| G1 | 5.627 | 4.755 | 4.627 | 4.377 | n/a | 7.962 | 12.39 | - |
| A2 | 6.038 | 4.485 | 4.767 | 4.531 | 7.811 | 8.161 | n/a | - |
| C3 | 5.425 | 4.318 | 4.521 | 4.402 | 5.179 | 7.627 | n/a | 8.247, 6.893 |
| A4 | 5.882 | 4.548 | 4.717 | 4.473 | 7.041 | 7.963 | n/a | - |
| G5 | 5.597 | 4.322 | 4.443 | 4.407 | n/a | 7.270 | 13.45 | - |
| C6 | 5.500 | 4.247 | 4.346 | 4.350 | 5.128 | 7.445 | n/a | 8.316, 6.642 |
| U7 | 5.561 | 4.278 | 4.467 | 4.382 | - | 7.642 | 10.09 | n/a |
| G8 | 5.633 | 4.529 | - | - | n/a | 7.802 | 13.20 | - |
| C9 | 5.526 | 4.354 | 4.443 | - | 5.192 | 7.764 | n/a | 8.555, 6.787 |
| U10 | 5.530 | 4.626 | - | - | 5.364 | 7.953 | 13.61 | n/a |
| G11 | 5.775 | 4.421 | 4.580 | 4.476 | n/a | 7.761 | 12.60 | - |
| U12 | 5.475 | 4.202 | 4.464 | 4.377 | 5.117 | 7.795 | 14.60 | n/a |
| C13 | 5.828 | 3.924 | 4.165 | 4.442 | 5.631 | 7.746 | n/a | 8.360, 7.027 |

$^{1}$H NMR chemical shifts of **3** in the r(CUG)-**3** complex:

| Hydrogen | Chemical shift |
|---|---|
| H1 | 8.718 |
| H2 | 8.060 |
| H3 | 7.814 |
| H4 | 6.735 |
| H5 | 6.886 |
| H6 | 6.919 |
| H7 | 7.383 |
| H8 | 6.950 |
| H15/16/17 | 2.489 |
| H18/19/20 | 3.770 |

Chemical shifts are in ppm units. Non-exchangeable protons were assigned at 35 °C. Exchangeable protons were assigned at 5 °C.

#### SYNTHETIC METHODS

*Abbreviations*. Boc, tert-butoxycarbonyl; DIPEA, diisopropylethylamine; DMF, *N*, *N*-dimethylformamide; Hex, hexanes; EtOAc, ethyl acetate; DMSO, dimethylsulfoxide; DCM, dichloromethane; DMAP, dimethylaminopyridine; EDCI, 1-Ethyl-3-(3-dimethylaminopropyl) carbodiimide;MeOH, methanol; NBS, *N*-bromosuccinimide; Rf, retention factor; TEA, triethylamine; TFA, trifluoroacetic acid; TLC, thin layer chromatography

*General*. Note the syntheses of compounds **1**[3] and **3**[4] were previously reported. The following were purchased from Acros Organics: 4-formylbenzoic acid, diethylenetriamine, and 4-nitrobenzaldehyde. *N*, *N*-dimethylformamide (DMF, anhydrous) was purchased from EMD and used without further purification. Claricep S-series prepacked silica columns were purchased from Agela-Technologies.

Flash chromatography was carried out on a Biotage Isolera One Automated Flash Chromatography System with prepacked Claricep S-series (40-60 μm). Normal-phase flash columns of various sizes were used.

Preparative HPLC was performed using a Waters 1525 Binary HPLC pump equipped with a Waters 2487 dual absorbance detector system and a Waters Sunfire C18 OBD 5 μm, 19 x 150 mm S-14 column. Absorbance was monitored at 254 nm and 345 nm. A linear gradient with a flowrate of 5 mL/min from 0-100% methanol in water with 0.1% (v/v) TFA over 100 min was used for small molecule purification. Purity was assessed by an analytical HPLC using a Waters Symmetry C18 5 μm, 4.6 x 150 mm column with a flow rate of 1 mL/min and a linear gradient from 0-100% methanol in water with 0.1% (v/v) TFA over 60 min. Absorbance was monitored at 254 nm and 345 nm.

$^1$H NMR spectra were collected on a Bruker 400 MHz NMR spectrometer. Mass spectra were recorded on a 4800 plus MALDI-TOF/TOF analyzer mass spectrometry an α-cyano-4-

hydroxycinnamic acid matrix. The concentration was determined in water using a molecular extinction coefficient (22501 M-1 cm-1 at 290 nm).

**Synthesis of compound 2:**

**Scheme 1:** Synthetic method for compound **2**.

***Tert*-butyl 2-(4-(4-formylbenzamido)phenyl)-4,5-dihydro-1*H*-imidazole-1-carboxylate (S3)**: 4-carboxybenzaldehyde (**S1**; 0.5 g, 3.33 mmol), tert-butyl 2-(4-aminophenyl)-4,5-dihydro-1H-imidazole-1-carboxylate (**S2**, synthetic methods below; 0.783 g, 3 mmol) , EDCI (0.958 g, 5 mmol) and DMAP (0.008 g, 2 mol%) were stirred at room temperature in DCM (15 mL) for 12 h. After the reaction was complete as monitored by TLC, the compound was purified by flash chromatography using Hex/EtOAc. In brief, the column was prepared with 50:50 Hex/EtOAc + 1% (v/v) TEA, and the solvent strength was increased to 10:90 Hex/EtOAc + 1% (v/v) TEA. Compound **S3** was obtained as a white colored solid in 22% yield (0.290 g, 0.737 mmol). $^1$H NMR, $CDCl_3$ (400 MHz) δ: 1.32 (s, 9H, 9H, $(CH_3)_3$OCO), 3.90-4.02 (m, 4H, $CH_2$), 7.52-7.54 (d, 2H, Ar*H*), 7.66-7.69 (d, 2H, Ar*H*), 7.98-8.04 (m, 4H, Ar*H*), 8.24 (s, 1H, C*H*O), 10.11 (s, 1H, CON*H*).

**4-(1-(2-aminoethyl)-4,5-dihydro-1*H*-imidazol-2-yl)-N-(4-(4,5-dihydro-1*H*-imidazol-2-yl)phenyl)benzamide (2)**: Compound **S3** (0.275 g, 0.699 mmol) was dissolved in DCM (15 mL) to which tert-butyl (2-((2-aminoethyl)amino)ethyl)carbamate (**S4** (synthetic methods below); 0.170 g, 0.839 mmol) was added, and the reaction was stirred at room temperature for 3 h. After 3 h, NBS (0.162 g, 0.909 mmol) was added and stirred overnight at room temperature. A new spot was observed by TLC (10:1 DCM/MeOH with 5% (v/v) TEA). The crude reaction mixture was then treated with 30% (v/v) TFA/DCM for 4 h. The solvent was evaporated, and compound was purified using reverse phase C18 column (solvent A: Water + 0.1% (v/v) TFA; solvent B: Methanol + 0.1% (v/v) TFA) to obtain **2** (TFA salt) as a pale white colored solid in 24% yield over 2 steps (0.120 g, 0.167 mmol). $^{1}$H NMR, DMSO-*d6* (400 MHz) δ: 4.07 (s, 8H, C$H_2$), 7.67-7.69 (d, 2H, Ar*H*), 8.01 (s, br, 2H, N$H_2$), 8.06-8.12 (m, 4H, Ar*H*), 8.21-8.23 (d, 2H, Ar*H*), 10.61 (s, 1H, N*H*), 10.77 (S, 2H, N*H*), 10.97 (S, 1H, CON*H*).

**Scheme 2**

**Scheme 2**: Synthetic method for intermediate **S4**.

**2-(4-nitrophenyl)-4,5-dihydro-1*H*-imidazole (S7)**: 4-nitrobenzaldehyde (**S6**; 1.5 g, 9.93 mmol) was added to a round bottom flask, to which ethylenediamine (0.716 g, 11.9 mmol) was added in *t*-BuOH (40 mL). The reaction mixture was stirred at room temperature for 30 min, after which $K_2CO_3$ (4.12 g, 29.78 mmol) and iodine (3.15 g, 12.41 mmol) were added. The reaction was stirred at 70 °C for 3 h and then brought to room temperature. Sodium sufite solution was added to quench excess iodine. An orange-colored compound precipitated, which was filtered, washed with water and ether, and dried overnight. Compound **S7** was obtained as a dark orange colored solid in 95% yield (1.81g, 9.47 mmol). $^1$H NMR, DMSO-*d6* (400 MHz) δ: 3.43-3.48 (t, 2H, C$H_2$), 3.84-3.89 (t, 2H, C$H_2$), 7.20 (s, 1H, N*H*), 8.06-8.08 (d, 2H, Ar*H*), 8.28-8.31 (d, 2H, Ar*H*).

***Tert*-butyl 2-(4-nitrophenyl)-4,5-dihydro-1*H*-imidazole-1-carboxylate (S8)**: Compound **S7** (1.2 g, 6.28 mmol), boc anhydride (2.05 g, 9.41 mmol), and TEA (0.953 g, 9.41 mmol) were added to DMF (20 mL) in a round bottom flask and stirred at room temperature for 4 h. Once the reaction was complete as monitored by TLC, water was added, which resulted in precipitation of a pale-yellow colored solid. The solid was filtered and dried overnight. Compound **S8** was obtained as a pale yellow colored solid in 84% yield (1.63g, 5.25 mmol). Rf = 0.7 (10:1 DCM/MeOH). $^1$H NMR, DMSO-*d6* (400 MHz) δ: 1.20 (s, 9H, $(CH_3)_3$OCO), 3.91 (s, 4H, C$H_2$), 7.73-7.76 (d, 2H, Ar*H*), 8.25-8.28 (d, 2H, Ar*H*).

***Tert*-butyl 2-(4-aminophenyl)-4,5-dihydro-1*H*-imidazole-1-carboxylate (S2)**: Pd/C (0.1 g) was added to the hydrogenation vessel followed by addition of EtOH (5 mL). Compound **S8** (1 g, 3.43 mmol) was added next to the reaction vessel followed by addition of 20 mL EtOH. The reaction was carried out at room temperature under 10 atm pressure for 3 h. The reaction mixture was filtered through a plug of celite and washed with methanol to ensure maximum recovery of product. Solvent was evaporated to give pale white colored solid in 92% yield. (0.823 g, 3.15 mmol). $^1$H NMR, DMSO-*d6* (400 MHz) δ: 1.26 (s, 9H, $(CH_3)_3$OCO), 3.68-3.73 (m, 2H, C$H_2$), 3.79-3.84 (m, 2H, C$H_2$), 5.43 (s, 2H, N$H_2$), 6.49-6.51 (d, 2H, Ar*H*), 7.15-7.17 (d, 2H, Ar*H*).

***Tert*-butyl (2-((2-aminoethyl)amino)ethyl)carbamate (S4)**: Diethylenetriamine (**S9**; 3 g, 29.08 mmol) was dissolved in dioxane (40 mL) and stirred at room temperature for 10 min. Boc anhydride (0.825 g, 3.78 mmol) was dissolved in 20 mL dioxane and slowly added to the solution of diethylenetriamine and stirred at room temperature for 24 h. After the completion of reaction, the dioxane was evaporated, and the residue was dissolved in water and extracted with DCM (20 mL× 3). The pooled fractions were dried over $Na_2SO_4$ and evaporated to give crude intermediate **S4** (17% crude yield) as white oily liquid (1 g, 4.92 mmol). The compound was used without further purification. $^1$H NMR, $CDCl_3$ (400 MHz) δ: 1.45 (s, 9H, $(CH_3)_3OCO$), 2.66-2.69 (t, 2H, $CH_2NH_2$), 2.72-2.75 (t, 2H, $NH_2CH_2CH_2NH$), 2.79-2.81 (t, 2H, $NHCH_2CH_2NHBoc$), 3.21-3.25 (q, 2H, $NHCH_2CH_2NHBoc$).

**NMR Spectra for 2 and intermediates thereof:**

$^{1}H$ NMR spectrum of compound **S3:**

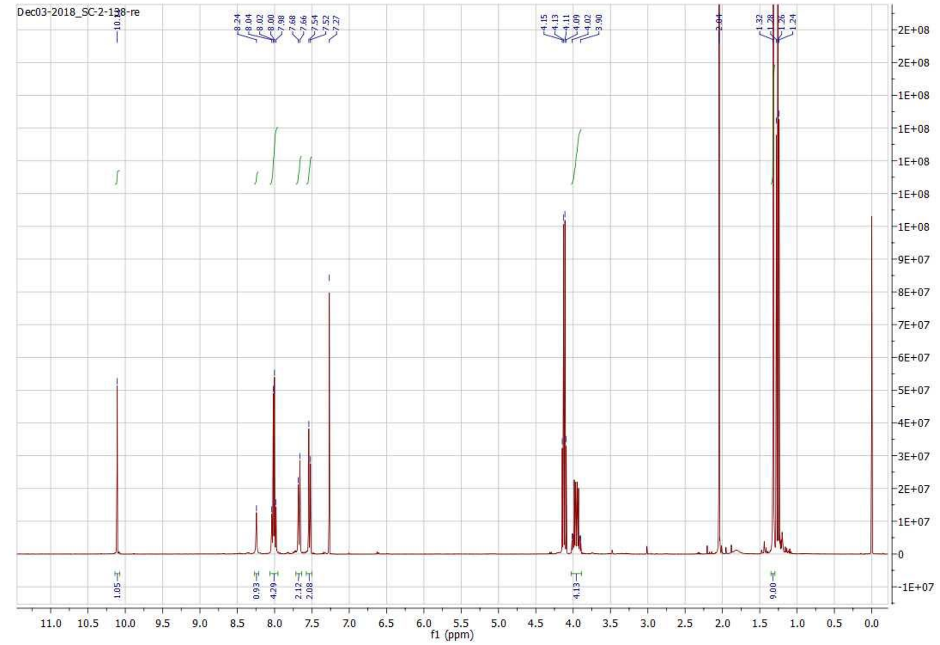


$^{1}H$ NMR spectrum of compound **2**:

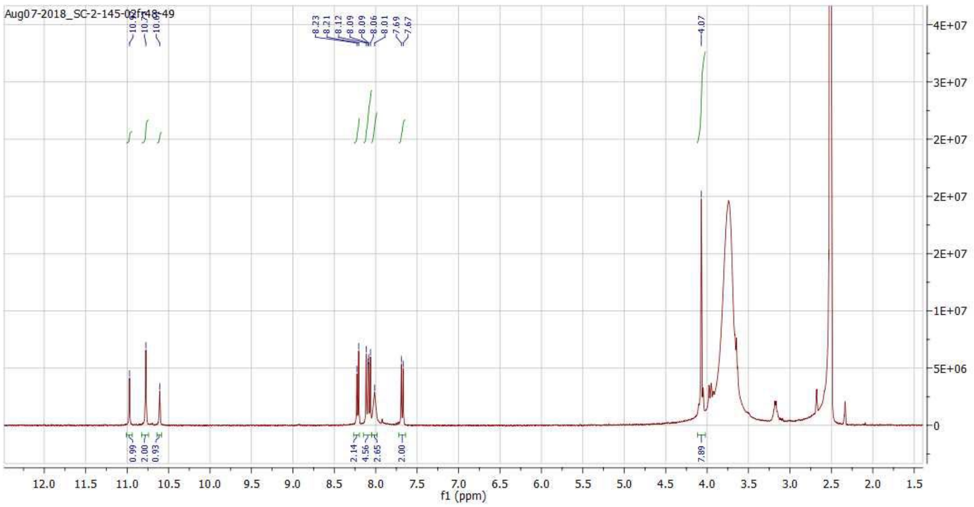

$^1$H NMR spectrum of compound **S7**:

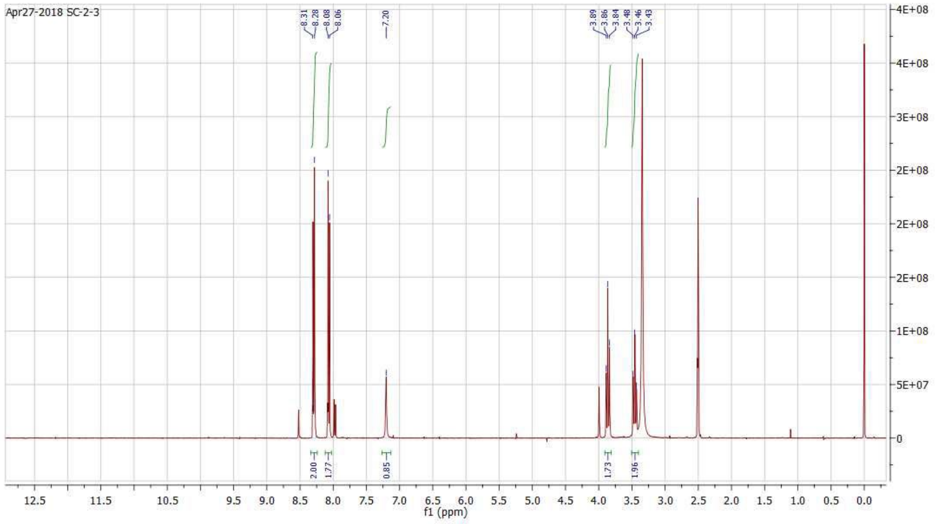


$^1$H NMR spectrum of compound **S8**:

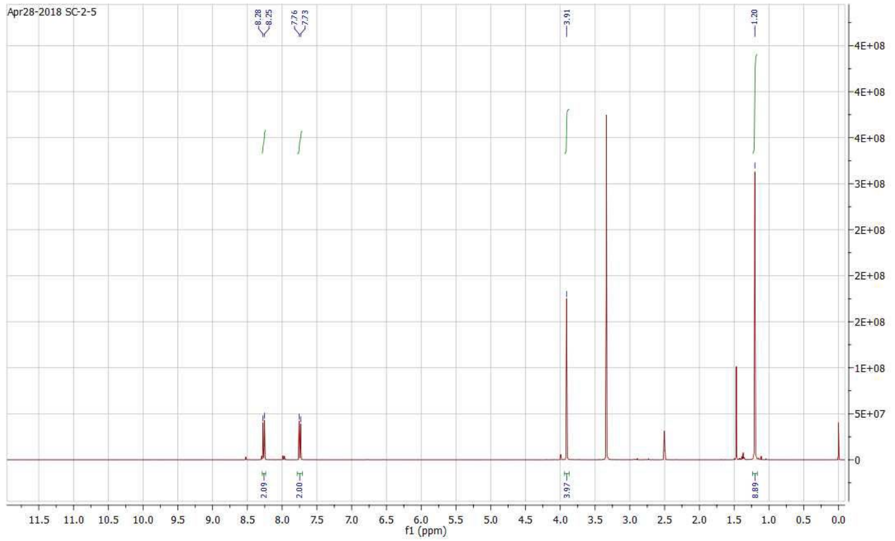

$^1H$ NMR spectrum of compound **S2**:

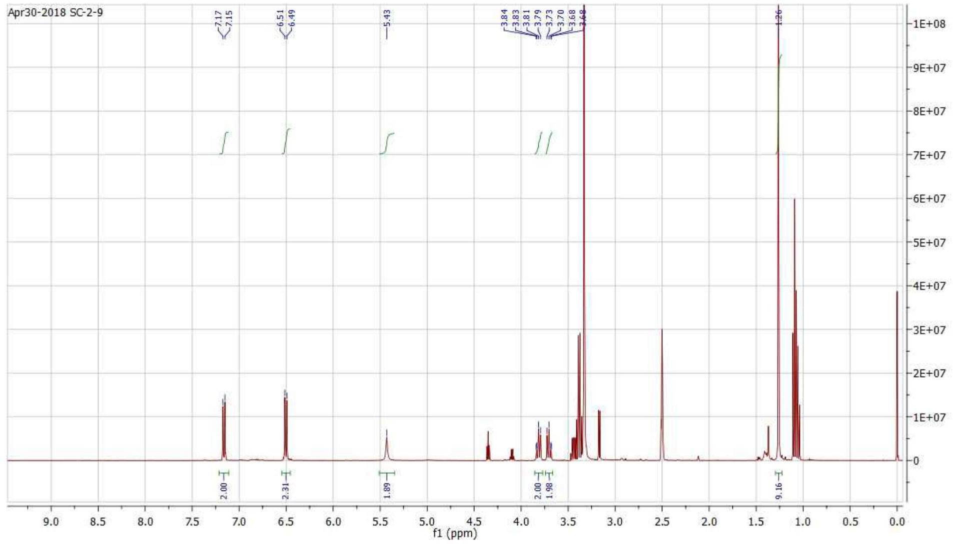


$^1H$ NMR spectrum of compound **S4**:

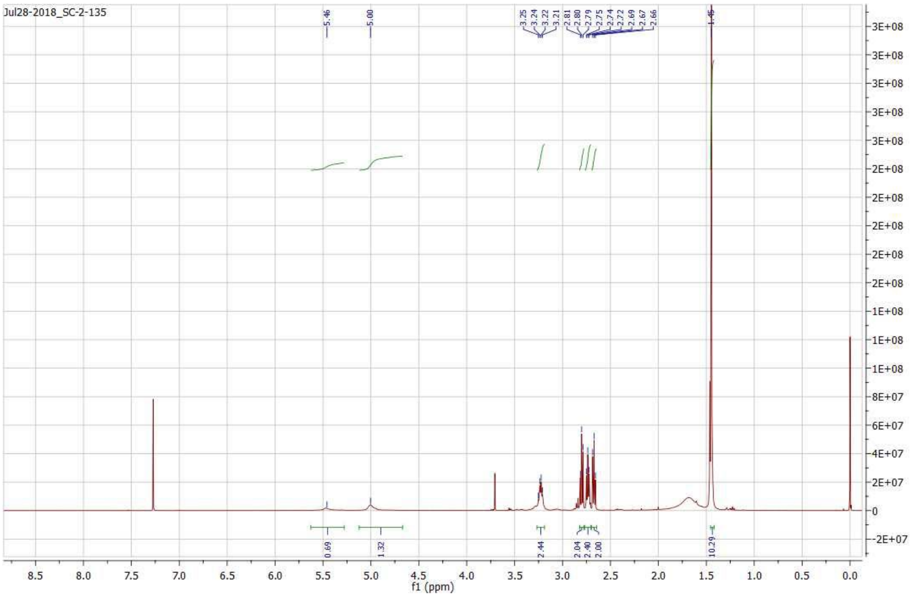
